# Rationally seeded computational protein design

**DOI:** 10.1101/2023.08.25.554789

**Authors:** Katherine I. Albanese, Rokas Petrenas, Fabio Pirro, Elise A. Naudin, Ufuk Borucu, William M. Dawson, D. Arne Scott, Graham J. Leggett, Orion D. Weiner, Thomas A. A. Oliver, Derek N. Woolfson

## Abstract

Computational protein design is advancing rapidly. Here we describe efficient routes to two families of α-helical-barrel proteins with central channels that bind small molecules. The designs are seeded by the sequences and structures of defined *de novo* oligomeric barrel-forming peptides. Adjacent helices are connected using computational loop building. For targets with antiparallel helices, short loops are sufficient. However, targets with parallel helices require longer connectors; namely, an outer layer of helix-turn-helix-turn-helix motifs that are packed onto the barrels computationally. Throughout these pipelines, residues that define open states of the barrels are maintained. This minimises sequence sampling and accelerates routes to successful designs. For each of 6 targets, just 2 – 6 synthetic genes are made for expression in *E. coli*. On average, 80% express to give soluble monomeric proteins that are characterized fully, including high-resolution structures for most targets that match the seed structures and design models with high accuracy.

## Introduction

Approaches to *de novo* protein design have developed considerably over the past four decades^1–5^. Early in the field, *minimal design* used straightforward chemical principles, particularly the patterning of hydrophobic and polar residues, to deliver peptide assemblies and relatively simple protein architectures. Largely, this gave way to *rational design,* in which sequence design was augmented by understood sequence-to-structure relationships garnered from bioinformatics and biochemical experiments. This delivered more-varied and more-robust designs. In parallel, *computational design* emerged allowing the realisation of concepts such as fragment-based and parametric backbone design, and methods for fitting *de novo* sequences onto these scaffolds^2,6,7^. In turn, this has led to increasingly complex designs of new structures and functions for both water-soluble and membrane-spanning proteins^3^. Currently, the field is undergoing another step-change with the application of data-driven/deep-learning methods to generate *de novo* protein sequences, structures, and functions^5,8–16^. These methods have the potential to democratise protein design^11,17^ and to promote its application in biotechnology^18,19^, cell biology^20^, materials science^21,22^, and medicine^23–25^.

Despite this progress, considerable challenges remain to realise the full promise of *de novo* protein design both in terms of advancing fundamental protein science and making it a robust and reliable alternative to engineering natural proteins for the application areas listed above. Current challenges include generating starting backbones that are designable^11,26,27^ towards a desired function, and increasing the success rates of converting *in silico* design into experimentally confirmed proteins^8,28–30^. In addition to these practical issues, we must address the concern that whilst deep-learning approaches will continue to advance our abilities to design protein structures and functions in new and unforeseen ways, it is less clear that they will necessarily improve our basic understanding of protein structure and function. Here, to bridge this gap, we advocate for and demonstrate the potential of combining rational and computational protein design. In this way, we deliver robust new protein sequences and structures—namely, barrel-like proteins with accessible and functionalisable central channels—rapidly and with high success rates.

Over the last decade, a range of oligomeric α-helical barrels (αHBs) have been designed based on self-assembling peptides that encode highly specific and stable coiled-coil interactions^31,32^. These *αHB peptides* are interesting *de novo* scaffolds because of their stability, robustness to mutation, and the potential to functionalise their internal lumens^18,33–35^. However, the scope for developing these is limited because they are peptide-based and largely homo-oligomeric. One solution to increasing the utility of αHBs is to connect the helices into single polypeptide chains that can be produced by the expression of synthetic genes. However, this is not straightforward, as the majority of αHBs have parallel helices. Here we describe two routes to *αHB proteins*. In the first, we design new antiparallel αHB peptides, and then connect adjacent helices into single chains using short loops (Fig. 1b). Second, for the existing all-parallel αHB peptides the helices are connected by longer structured loops (Fig. 1c). In both cases, we test several approaches to computational loop building. A key aspect of our design process is that it uses well-understood sequence-to-structure relationships garnered from the oligomeric peptides as rules to seed the designs rather than designing entirely new sequences. This speeds up the design process, produces robust *in silico* models, limits the number of constructs that have to be tested, and gives high success rates of experimentally validated targets (Fig. 1d).

**Fig. 1:**
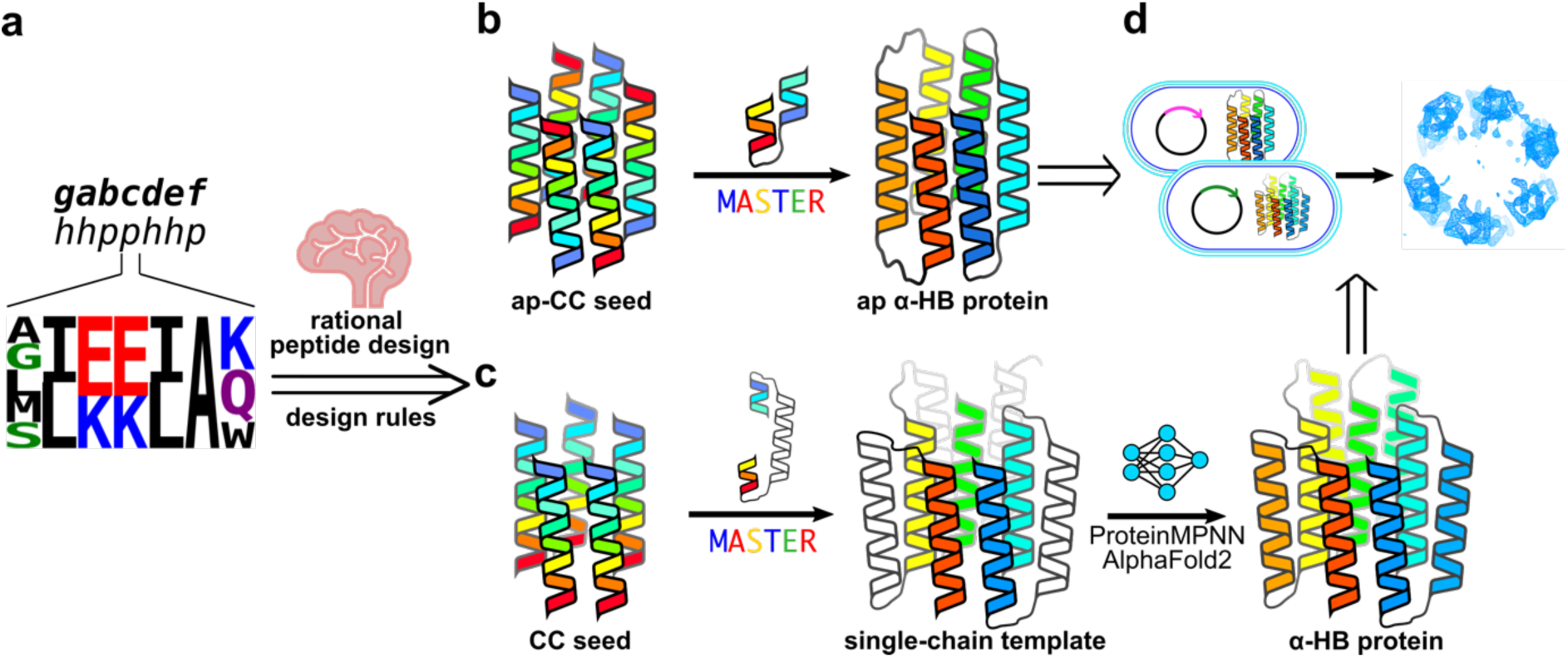
Pipeline for rationally seeded computational design of *de novo* protein folds. **a**, Robust sequence-to-structure relationships for coiled-coil oligomers are used as rules to seed the design of new protein scaffolds. **b, c,** Antiparallel (**b**) and parallel (**c**) α-helical barrel protein (αHB) design targets. For both targets, MASTER^36,37^ is used to search known experimental protein structures for segments with the potential to connect adjacent helices and generate single-chain models. For the antiparallel designs (**b**), the sequences and structures of identified short connectors can be used directly. However, the parallel targets require longer structured loops (**c**), for which we target helix-turn-helix-turn-helix motifs. ProteinMPNN^8^ and AlphaFold2^38,39^ are then used iteratively to optimise the sequences and models of these 3-helix bundle motifs. **d**, For each design, a small number of synthetic genes are made and expressed in *E. coli* for biophysical and structural characterisation. Colour scheme: peptide and protein chains are shown in chainbows from the *N* to the *C* termini (blue to red), except for the initially placed central helices of the helix-turn-helix-turn-helix motifs in the parallel designs, which are shown in white.

## Results

### New peptide rules deliver rarer antiparallel aHBs

To date, most αHB peptides have all-parallel arrangements of helices^32^. Because of the extended connections required (Fig. 1c), turning these into single-chain αHB proteins is not trivial. Therefore, first, we addressed the challenge of making less-common antiparallel αHB peptides^40–42^, which could be converted to αHB proteins using short linkers between adjacent helices (Fig. 1b). Based on the collective understanding of coiled coils^32^, we tested an informed subset of synthetic peptides that could potentially form antiparallel hexamers. Our designs focussed on the g, a, d, and e sites of the classical coiled-coil, heptad sequence repeat, gabcdef, as these contribute most to the helix-helix interfaces (Fig. 2a). Specifically, we made 20 sequence combinations with g = Ala, Gly, Leu, Met, or Ser, and a and d = Ile or Leu. These were installed into 4-heptad peptide sequences with a common background comprising: e = Ala^42–44^; ‘bar-magnet’ charge patterning of Glu and Lys at b and c to favour antiparallel coiled-coil assemblies^42,44,45^; and f = Gln, Lys, and Trp to aid helicity and solubility, and to add a chromophore. The 20 sequences (Supplementary Table 1) were made by solid phase peptide synthesis, purified by HPLC and confirmed by mass spectrometry (Supplementary Fig. 1). Each peptide was tested for α-helicity and thermal stability by circular dichroism (CD) spectroscopy (Fig. 2e,f and Supplementary Figs. 2 and 3); oligomeric state by analytical ultracentrifugation (AUC), (Fig. 2g, Supplementary Table 2, and Supplementary Figs. 4 and 5); and accessible central channels using a dye-binding assay^34^ (Fig. 2h and Supplementary Fig. 6). Most of these formed hyper-stable, helical hexamers that bound dye to varying degrees (Supplementary Table 3).

**Fig. 2:**
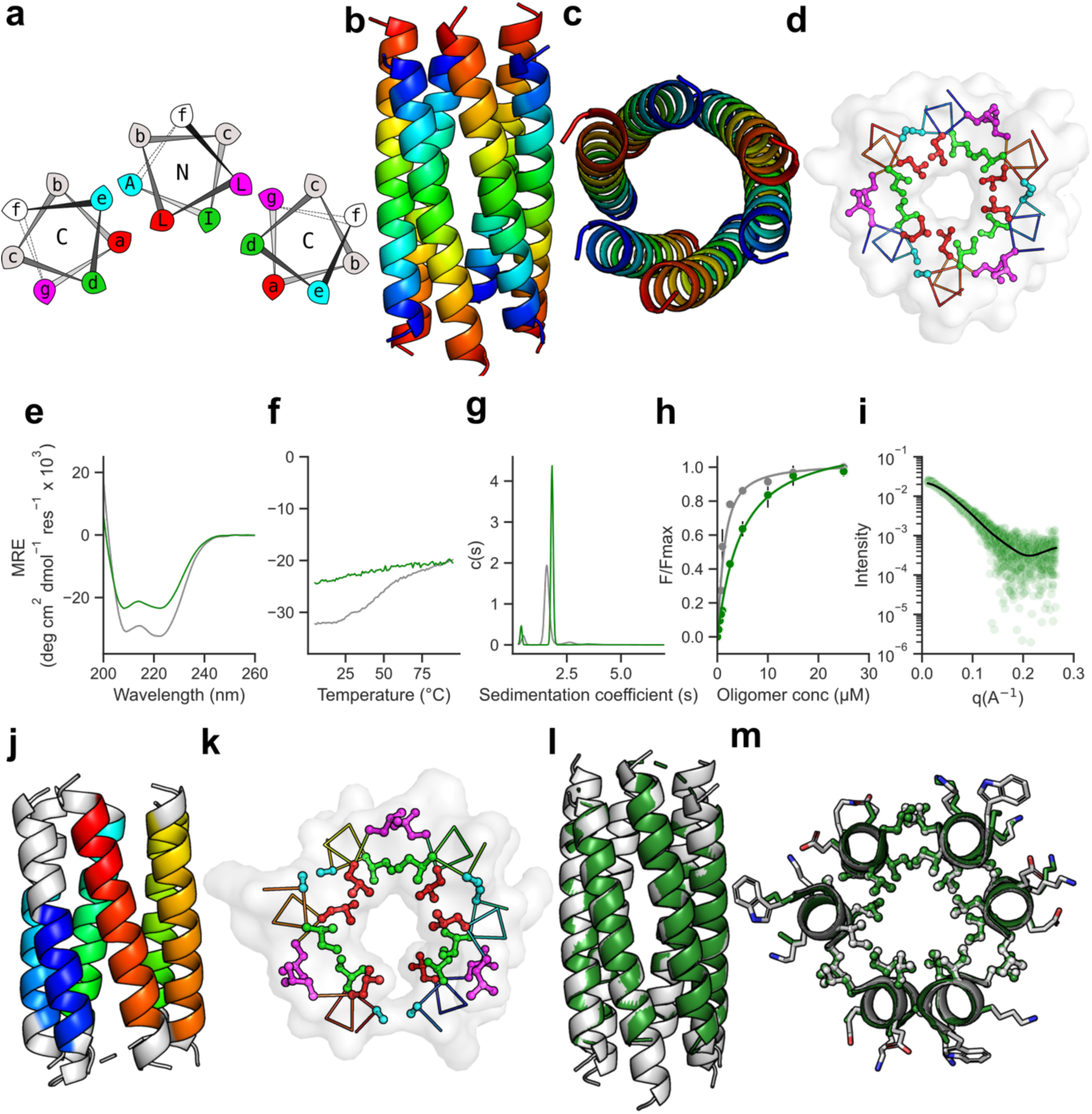
Biophysical and structural characterisation of the apCC-Hex peptide and the sc-apCC-6-LLIA protein. **a,** Helical-wheel representation of part of an antiparallel αHB highlighting the a – g heptad repeats: a sites, red; d, green; g, magenta; e, cyan; N and C labels refer to the termini of the helices closest to the viewer. **b** – **d**, 1.4 Å resolution X-ray crystal structure of apCC-Hex (PDB id, 8qab). **b, c,** Coiled-coil regions identified by Socket2^46^ (packing cutoff 7.0 Å) coloured as chainbows from *N* to *C* termini (blue to red). **d**, Slice through the structure for a heptad repeat with knobs-into-holes (KIH) packing coloured the same as in the helical wheel of panel **a**. **e – h**, Comparison of the biophysical data for the apCC-Hex αHB peptide (grey) and the sc-apCC-6-LLIA αHB protein (green). **e,** CD spectra recorded at 5 °C. **f,** Thermal responses of the α-helical CD signal at 222 nm. **g,** AUC sedimentation velocity data at 20 °C fitted to a single-species model; fits returned a peptide assembly of 18.7 kDa (hexamer), and a protein of 24.0 kDa (monomer), respectively. **h**, Fitted data for 1,6-diphenyl hexatriene (DPH) binding to the peptide and protein; fits returned K_D_ values of 0.8 ± 0.3 µM and 4.0 ± 0.4 µM, respectively. **i,** SEC-SAXS data for sc-apCC-6-LLIA fitted using FoXS^47,48^ to an AlphaFold2 model of the design, *x*^2^ = 1.50. **j**, 2.25 Å X-ray crystal structure of sc-apCC-6-LLIA (PDB id, 8qad) with coiled-coil regions identified by Socket2^46^ (packing cutoff 7.0 Å) coloured chainbow. **k**, Slice through the structure for a heptad repeat showing the KIH packing coloured as in panel **a**. **l**, **m,** Overlays of the experimental apCC-Hex (grey) and sc-apCC-6-LLIA protein (green) structures RMSD_bb_ = 1.177 Å. Conditions: CD spectroscopy, 50 µM peptide, 10 µM protein in PBS, pH 7.4; AUC, 100 µM peptide, 15 µM protein in PBS, pH 7.4; DPH binding, oligomer concentration 0 – 30 µM peptide, 0 – 30 µM protein in PBS, pH 7.4, 20 °C, final concentration 1 µM DPH (5% v/v DMSO); SEC-SAXS, 10mg/mL protein in PBS, pH 7.4.

From this set, we solved three high-resolution X-ray crystal structures. One, with g-a-d-e = Ala-Leu-Ile-Ala, revealed an antiparallel hexamer consistent with its solution-phase oligomer state (Supplementary Table 2). However, this was a collapsed bundle, conflicting with the solution-phase binding data that suggest this peptide can access an open αHB (Supplementary Table 3 and Supplementary Fig. 7). Another, with g-a-d-e = Gly-Leu-Ile-Ala, had promising solution-phase data for an open hexamer or heptamer (Supplementary Tables 2 and 3), but, interestingly, formed a collapsed antiparallel octamer in the crystal state (Supplementary Fig. 8). By contrast, g-a-d-e = Leu-Leu-Ile-Ala formed the targeted antiparallel hexameric open barrel with completely consistent solution-phase behaviour (Fig. 2b – d, Supplementary Table 3, and Supplementary Fig. 9)^42^. We named this peptide apCC-Hex-LLIA, and systematically as apCC-Hex. This process illustrates the importance of establishing robust rules for the next stage of the protein-design pipeline.

### Short loops yield an antiparallel aHB protein

Using the experimental apCC-Hex structure as a seed, we designed short loop sequences computationally to connect adjacent helices to generate an up-down α-meander structure (Fig. 1b). We tested three approaches: first, and most simply, we took loops from the literature and the PDB to span the distances between the *C* and *N* termini of the helices^42,49–51^; second, we applied MASTER^36,37^ (Supplementary Table 4) to find tertiary fragments to link the helices; and, third, we used the ColabPaint implementation of Protein Inpainting^9^ to bridge the gap (https://github.com/polizzilab/design_tools). The resulting single-chain templates were used in a computational screen to find the best-fitting combinations of residues at the g-a-d sites (with e fixed as Ala). This was guided by the privileged residue combinations from the experiments with synthetic peptides (Supplementary Table 3). Models for the remaining g-a-d combinations with different loop sequences were built using AlphaFold2^38,39^ in single-sequence mode (Supplementary Figs. 10 – 12) and assessed by pLDDT from AlphaFold2, and RMSD to the parent apCC-Hex starting scaffold. We generated 7 sequences with different g-a-d-e combinations and loop-building methods (Supplementary Tables 5 and 6).

Synthetic genes for all except 2 of the 7 expressed in *E. coli* (Supplementary Tables 6 – 8). As the peptide assemblies were hyper-thermally stable, we heat treated the cell lysate (75 °C for 10 mins) and subjected the soluble fraction to immobilised-metal-affinity (IMAC) and size-exclusion chromatography (SEC) to yield highly pure proteins in a minimal number of steps (Supplementary Fig. 13). CD spectroscopy showed that all 5 proteins were α-helical and hyper-thermally stable structures (Fig. 2e,f and Supplementary Figs. 14 and 15); AUC confirmed them as monomers (Fig. 2g, Supplementary Table 7, and Supplementary Fig. 16); and dye binding indicated they had accessible hydrophobic cavities (Fig. 2h and Supplementary Fig. 17). These data (Supplementary Table 8) were supported by size-exclusion chromatography coupled with small-angle X-ray scattering (SEC-SAXS) data, which fitted to their respective AlphaFold2 models with good χ^2^ values^47,48^ (Fig. 2i, Supplementary Table 9, and Supplementary Fig. 18). Finally, we obtained two high-resolution X-ray crystal structures for sequences generated using MASTER^36,37^: one directly derived from apCC-Hex, g-a-d-e = Leu-Leu-Ile-Ala (Fig. 2j-m and Supplementary Fig. 19); and another, g-a-d-e = Ser-Leu-Leu-Ala, one of the tighter dye-binding proteins characterised (Supplementary Fig. 20). The sequences and structures were named sc-apCC-6-LLIA and sc-apCC-6-SLLA, respectively, for single-chain antiparallel coiled-coil proteins with 6 central helices.

### Structured a-helical motifs link parallel helices

The parallel αHB proteins required a different design approach as sequence-to-structure relationships for the g-a-d-e positions were available to seed the designs^31,52,53^, but connecting adjacent parallel helices was not straightforward because of the need to span ≈40 Å along the structures (Fig. 1c). Indeed, previously we had made several unsuccessful attempts to link parallel helices using polyproline helix-based linkers^54^. Therefore, we tested whether MASTER^36,37^ could find better α-helical templates from the PDB to address this. We exploited the *C_n_* symmetry of the parallel αHB peptides to generate helix-turn-helix-turn-helix (h-t-h-t-h) units, which could be repeated about the *C_n_* axis to close structures with *n* central helices and *n-1* buttressing helices (Fig. 1c). To find h-t-h-t-h units, we restricted the MASTER searches to a non-redundant set of 3-helix coiled-coil bundles from the CC+ database^55,56^. This delivered several candidate backbones from which we chose the lowest RMSD hit for each target (Supplementary Table 4).

Adding sequences to the new backbones required optimisation of side-chain interactions in both the external 3-helix bundle and the internal barrel (Fig. 3a). For the latter, again, sequence-to-structure relationships from existing αHB peptides seeded and accelerated sequence design. This is best illustrated by example (Supplementary Fig. 22). For example, the g-a-d-e combination Ala-Leu-Ile-Ala defines the parallel heptamer, CC-Hept (PDB id, 4pna)^31^. Therefore, these positions were fixed in the 7 parallel inner helices of a 13-helix template derived from the backbone-generation procedure (Figs. 1c and 3b). Initially, the rest of the sequence was optimised using ProteinMPNN^8^. However, as others report^57^, we found that this placed hydrophobic residues on the solvent-exposed surface of the structure. To remedy this, as the outer helices were also based on coiled coils, we fixed the exposed b, c, and f sites to combinations of Glu, Lys, and Gln (Supplementary Fig. 23). Initially, 100 sequences were generated, filtered based on core packing, Rosetta energy, and charge, and modelled with AlphaFold2^38,39^ (Supplementary Fig. 22). The model with best pLDDT score was used to initiate another round of sequence design. At this point, we replaced the fixed constraint on the outermost b-c-f residues with a Lys/Glu bias in ProteinMPNN^8^, followed by a surface hydrophobicity filter within Rosetta. This gave similar charge distributions and exposed-hydrophobic scores, but allowed less-repetitive sequences to be generated (Supplementary Fig. 24). Iterations were repeated until the energies and the RMSDs between the ProteinMPNN^8^ inputs and AlphaFold2^38,39^ outputs converged (Supplementary Fig. 24). For the sc-CC-7 target, this occurred after 3 rounds to give helical sequences (Fig. 3b).

**Fig. 3:**
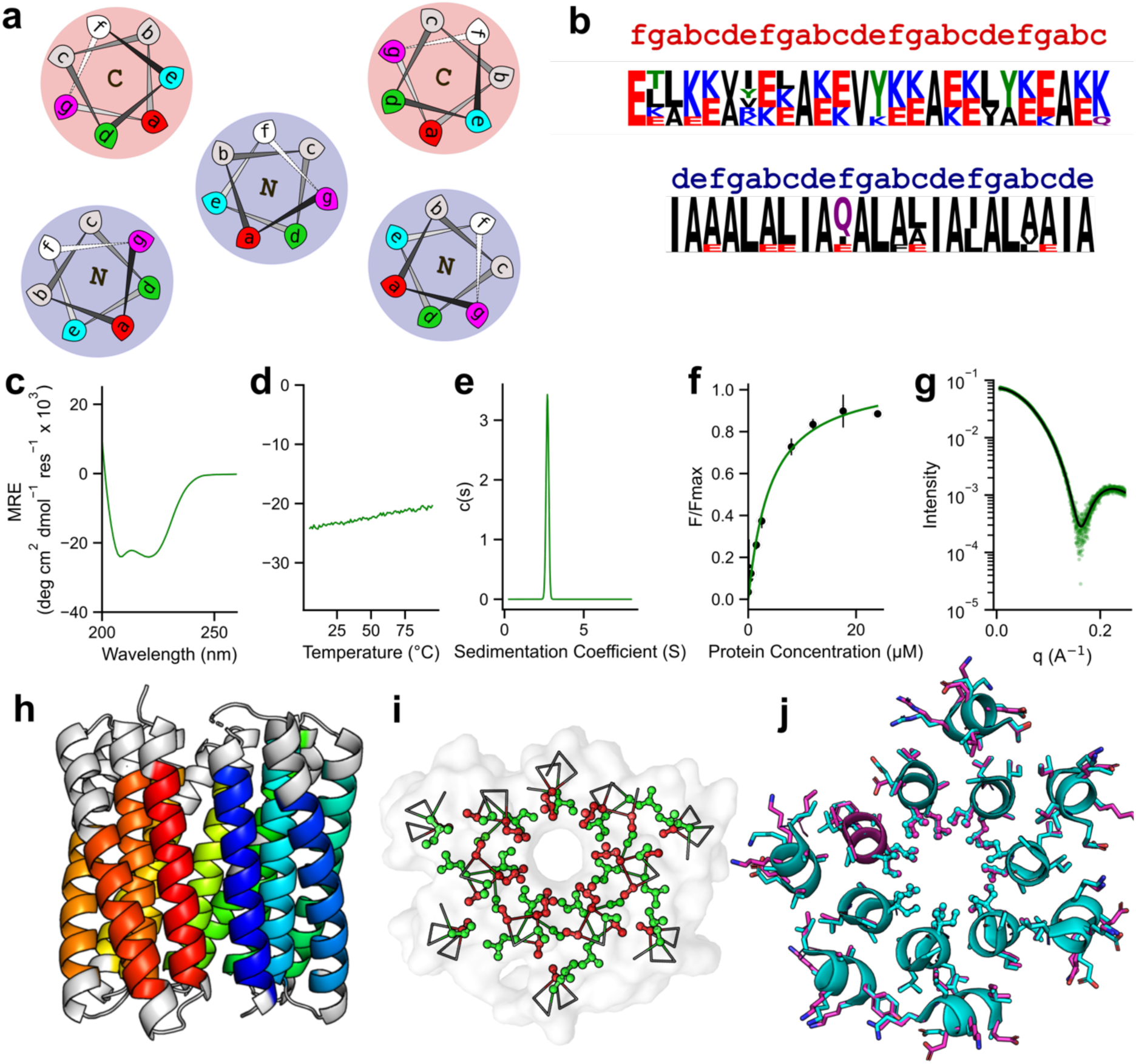
Biophysical and structural characterisation of sc-CC-7 *de novo* proteins. **a,** Helical-wheel representation for part of a parallel single-chain αHB showing the KIH packing for the buttressing helices (shaded red) and the inner barrel (shaded blue): *a* sites, red; *d*, green; *g*, magenta; *e*, cyan; N and C labels refer to the termini of the helices closest to the viewer. **b,** Sequence pileups and registers for the inner (blue register) and buttressing (red register) helices of sc-CC-7-LI. **c, d,** CD spectrum recorded at 5 °C **(c)** and thermal-response curve **(d)** for sc-CC-7-LI. **e,** AUC sedimentation velocity data for sc-CC-7-LI fitted to a single-species model, which returned M_W_ = 37.4 kDa (monomer). **f,** Fitted binding data of DPH to sc-CC-7-LI, which returned K_D_ = 3.8 ± 0.8 µM. **g,** SEC-SAXS data fitted using the final AlphaFold2 model and FoXS, *x*^2^ = 1.43^47,48^. **h,** X-ray crystal structure of sc-CC-7-LI at 2.5 Å resolution (PDB id, 8qai). Coiled-coil regions identified by Socket2^46^ (packing cutoff 7 Å) are coloured as chainbows from *N* to *C* termini (blue to red). **i,** Slice through the structure for a heptad repeat showing the KIH packing with a-type knobs in red and d*-*type in green. **j,** Overlay of the middle helical turns from the sc-CC-7-LI structure (cyan) and the final AlphaFold2 model (magenta), RMSD_bb_ = 0.433 Å. Conditions: CD spectroscopy, 5 µM protein in PBS, pH 7.4; AUC conditions, 25 µM protein in PBS, pH 7.4; DPH binding conditions, 0 – 24 µM protein in PBS, pH 7.4. final concentration 0.5 µM DPH (5% v/v DMSO); SEC-SAXS conditions, 10 mg/mL protein in PBS, pH 7.4.

We chose 4 protein sequences with <85% sequence identity, high pLDDT, and low Rosetta energies for gene synthesis and expression in *E. coli* (Supplementary Tables 10 and 11). 2 of these expressed. As for the antiparallel designs, these were purified by heat treatment, centrifugation, and IMAC and SEC to render highly pure protein (Supplementary Fig. 25). One of these (sc-CC-7-80) was oligomeric by AUC, which although helical and thermally stable, was not characterised further (Supplementary Tables 12 and 13 and Supplementary Figs. 26 – 30). The other protein, named sc-CC-7-LI because of its a = Leu d = Ile core, was helical and fully resistant to heat denaturation as judged by CD spectroscopy (Fig. 3c, d, Supplementary Table 13, and Supplementary Figs. 26 and 27); monomeric by AUC (Fig. 3e, Supplementary Table 12, and Supplementary Fig. 28); and bound dye consistent with an accessible channel (Fig. 3f, Supplementary Table 13, and Supplementary Fig. 29). This was supported by SEC-SAXS data fit to the AlphaFold2 model^47,48^ (Fig. 3g, Supplementary Table 14, and Supplementary Fig. 30). We solved an X-ray structure for sc-CC-7-LI to 2.5 Å resolution (Fig. 3h – j). Finally, to test the robustness of the design to mutation, we substituted all 49 a (Leu) and d (Ile) sites of the central αHB for alternative design rules for parallel heptameric αHBs, *i.e*., a = Ile and d = Val^53^. This protein, sc-CC-7-IV, expressed well and was also folded as shown by CD spectroscopy and SEC-SAXS, hyperstable, monomeric, and bound the reporter dye (Supplementary Tables 10 – 14 and Supplementary Figs. 25 – 30).

### Seeded design rapidly accesses more aHB proteins

Encouraged by the successful design of sc-apCC-6 and sc-CC-7, we extended the seeded-design approaches to target αHB proteins with 5, 6, and 8 central helices (Supplementary Tables 15 – 28 and Supplementary Figs. 31 – 63). To seed the antiparallel 8-helix αHB protein design, we had two starting sequences: the aforementioned peptide with g-a-d-e = Gly-Leu-Ile-Ala, which formed a collapsed antiparallel 8-helix bundle; and, from a previous study, g-a-d-e = Ala-Ile-Ile-Ala with a different b-c-f background forms an open parallel octamer by X-ray crystallography^52^. Therefore, we extended the peptide screen introduced above to explore around this sequence space (Supplementary Table 1). The resulting synthetic peptides formed stable, helical, higher-order oligomers with accessible channels (Supplementary Table 3 and Supplementary Figs. 1 – 6). Octameric AlphaFold2^38,39^ models for these were generated and used to seed the computational design of single-chain αHB proteins. We used MASTER^36,37^ to find backbones to connect the helices (Supplementary Table 4). Next, ProteinMPNN^8^ was used to generate loop sequences, keeping the helical residues fixed and iterating with AlphaFold2^38,39^ to find sequences and models that were open αHBs with the highest pLDDT. This led to two designs: g-a-d-e = Ala-Ile-Ile-Ala, and g-(a-d)_2_(a-d)_2_-e = Gly-(Ile-Leu)_2_(Leu-Ile)_2_-Ala (Supplementary Tables 15 and 16 and Supplementary Figs. 31 and 32). In the latter, two a-d combinations are repeated through the first two and last two heptads, respectively.

Both designs expressed well (Supplementary Fig. 33), and the purified proteins were soluble, folded, thermally stable, monomeric and monodisperse, with accessible cavities (Supplementary Tables 17 and 18 and Supplementary Figs. 34 – 37). This was confirmed by SEC-SAXS and X-ray crystallography (Fig. 4, Supplementary Table 19, and Supplementary Figs. 38 and 39). We solved a 2.0 Å X-ray crystal structure for g-a-d-e = Ala-Ile-Ile-Ala, which we called sc-apCC-8 (Fig. 4a; Supplementary Fig. 39).

**Fig. 4:**
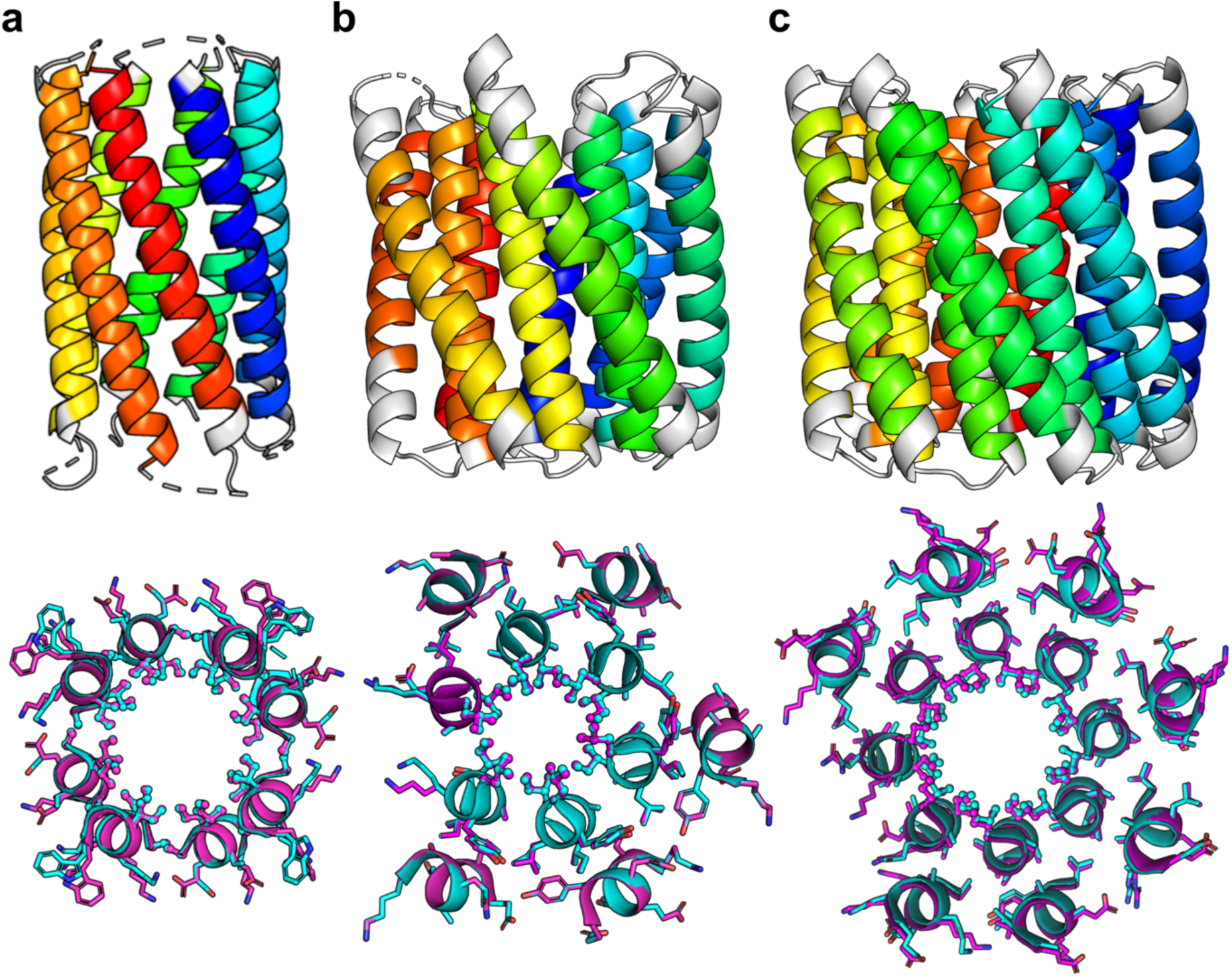
Structural characterisation of 6-helix and 8-helix targets. **(a – c, top),** X-ray crystal structures of **(a)** sc-apCC-8 at 2.0 Å resolution (PDB id, 8qaf), **(b)** sc-CC-6-95 at 2.8 Å resolution (PDB id, 8qag), and **(c)** sc-CC-8-58 at 2.35 Å resolution (PDB id, 8qah). Coiled-coil regions identified by Socket2^46^ (packing cutoff 7.5 Å for sc-apCC-8, sc-CC-6-95 and sc-CC-8-58 at 7.0 Å) are coloured as chainbows from *N* (blue) to *C* (red) termini. **(a – c, bottom),** Overlays for the middle helical turns of each crystal structure (cyan) and the corresponding AlphaFold2^38,39^ model (magenta); RMSD_bb_ = 0.413 Å **(a),** RMSD_bb_ = 0.300 Å **(b),** and RMSD_bb_ = 0.530 Å **(c)**.

For aHB proteins with inner barrels of 5, 6 and 8 parallel helices, we used seeds from existing peptide assemblies, with a slight modification of the 6-helix target CC-Hex2 (PDB id, 4pn9) to replace g = Ser in the peptide assembly with Ala, to avoid polar Ser at the h-t-h-t-h interface (Supplementary Tables 4, 20 – 28 and Supplementary Figs. 41 – 45)^31,52,53^. MASTER selected a similar right-handed h-t-h-t-h tertiary fragment to connect the helices of the 6- and 8-helix targets as it did for sc-CC-7 (Supplementary Table 4); specifically, from a *de novo* helical repeat protein (PDB id, 5cwq)^58^. However, and interestingly, for the 5-helix target, it returned a left-handed tertiary h-t-h-t-h template from the same design series (PDB id, 5cwi, Supplementary Table 4)^58^. This can be rationalised in that lower-order coiled-coil oligomers have left-handed, superhelical twists, whereas the larger helical assemblies have straighter superhelices^31,52,59^. For the three targets, 11 sequences were tested experimentally (Supplementary Tables 20 – 28 and Supplementary Figs. 46 – 63). Synthetic genes for all but 2 expressed in *E. coli*, and gave soluble proteins that were α-helical, monomeric, and thermally stable (Supplementary Figs. 46 – 63). The 5-helix-based proteins showed no dye binding suggesting collapsed bundles or simply that the cavities were too narrow to accommodate dye (Supplementary Table 27 and Supplementary Fig. 50). By contrast, the 6- and 8-helix-based targets bound dye consistent with accessible cavities, which were confirmed by SEC-SAXS and X-ray crystal structures (Fig. 4, Supplementary Tables 27 and 28, and Supplementary Figs. 52 – 63). Together, these additional designs delivered *de novo* proteins sc-CC-5, sc-CC-6, and sc-CC-8.

### The αHB proteins match the seeds and design models

We compared our experimental structures to the seed structures^31,52^, utilized tertiary fragments^58^, and final *in silico* design models generated by AlphaFold2^38,39^ (Supplementary Table 31). Because of changes from the full-sequence-design steps, we compared backbone atoms only. Apart from one structure, the backbone RMSD values for these comparisons are ≤1 Å (Supplementary Table 31). For the antiparallel αHB proteins, the seeds, models, and experimental structures for sc-apCC-6-LLIA and sc-apCC-8 are closely similar (Supplementary Table 31). The outlier is sc-apCC-6-SLLA (Supplementary Table 31), where the experimental structure and model differ at one of the Ser-Ser (g-g) helical interfaces (Supplementary Fig. 20e). Such polar contacts are notoriously difficult to model. For the parallel targets, the experimental structures show minor fraying at the *C* termini of the inner helices compared with the seeds and models, which appears to improve the packing of the external 3-helix bundles (Fig. 4b, Supplementary Table 31, and Supplementary Fig. 64). Along with the solution-phase data presented above, this high level of accuracy between the seeds, design models, and the experimental structures strongly supports the approach of rationally seeding computational design pipelines.

## Discussion

### αHB proteins are distinct from known and putative protein structures

In summary, our approach has delivered a set of *de novo* structures for antiparallel and parallel αHB proteins with 6 and 8, and 6, 7 and 8 central helices, respectively. We were interested how similar, if at all, these are to known protein structures and AlphaFold2 predicted models. Therefore, we used them as query structures in Foldseek^60^ to search the RSCB PDB^61,62^ and AlphaFold2-Swissprot databases^38,63^ (Fig. 5, Supplementary Tables 32 – 43, and Supplementary Fig. 65). This returned natural, *de novo*, and predicted α-helical bundles. However, most of the identified structures/models only partially overlapped with our queries, and the sequence identities of the overlapping regions and TM scores^64^ were generally low at <20% and of ≤0.5, respectively (Supplementary Tables 32 – 43). Moreover, most have spiralling and/or open structures rather than the cyclically closed structures that we targeted (Fig. 5).

**Fig. 5.**
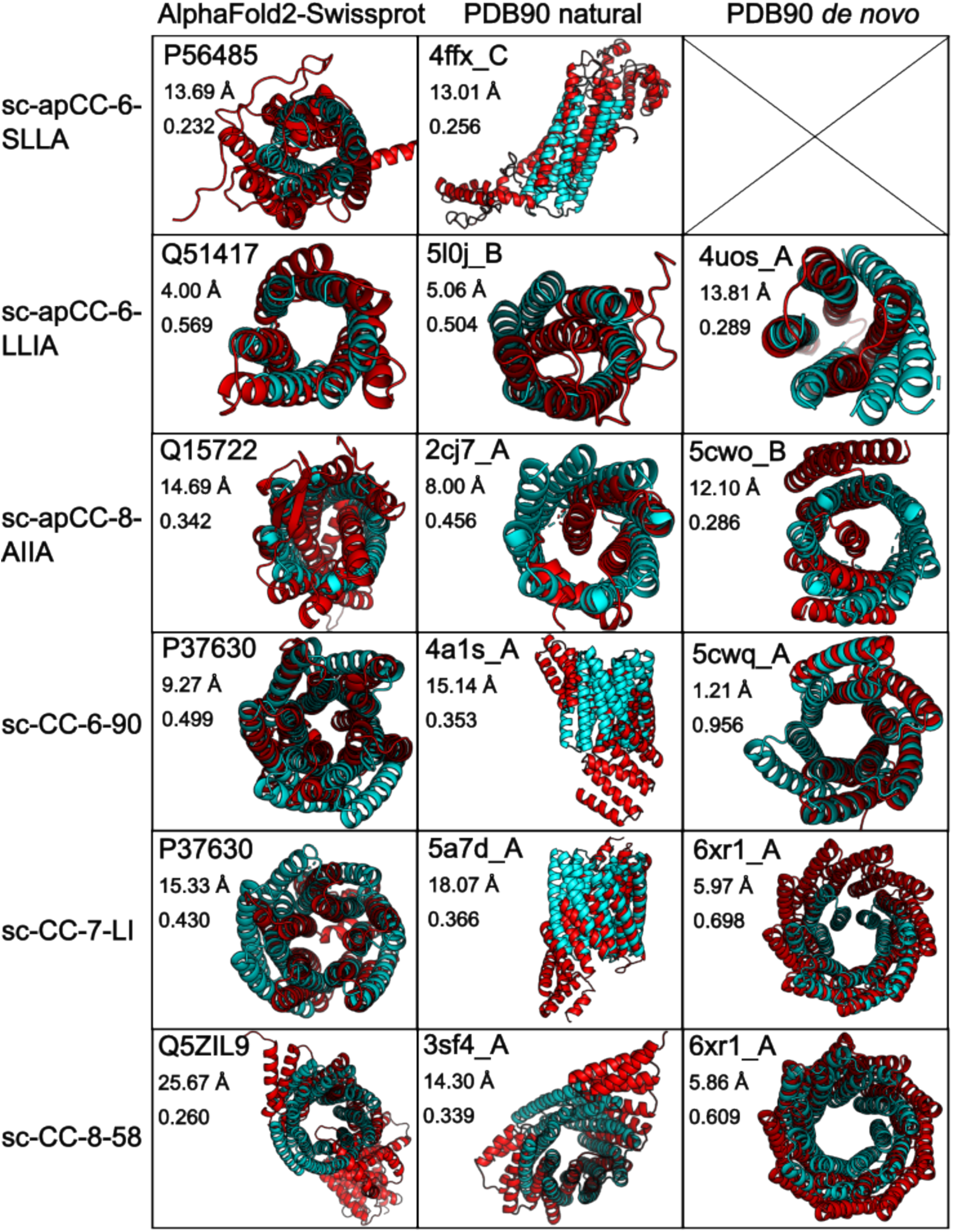
Foldseek^60^ comparison of *de novo* aHB proteins against existing and predicted protein folds. Each *de novo* αHB protein structure determined in this study (cyan) is overlaid with the top match from the AlphaFold2-Swissprot database,^38,63^ and natural and *de novo* sequences from the PDB^61,62^ (red). Within each box, the top value gives the ID of matched structure; the middle value is the backbone RMSD between the query and match; and the bottom value is the TM score^64^ between the two structures.

In more detail, for the antiparallel αHB proteins, sc-apCC-6-SLLA returned partial matches within proteins containing 4-helix bundles (Fig. 5 and Supplementary Tables 32 and 33). We found only hypothetical 6-helix bundles in the wider UniProt database^38,63^ (*e.g.*, UniProt id, A0A2G8LCW8, Supplementary Fig. 66). sc-apCC-6-LLIA recovered a 4-helix bundle from human vinculin (PDB id, 5l0j^65^), and a 6-helix bundle from the putative transporter protein, AmiS, from *P. aeruginosa* (UniProt id, Q51417^38,63^, Fig. 5 and Supplementary Tables 34 and 35). Socket2^46^ located KIH interactions in both of these, but only between pairs of helices (Supplementary Fig. 65). sc-apCC-8 gave mostly poor alignments to helical-repeat proteins (Fig. 5 and Supplementary Tables 36 and 37). Interestingly, we found a match to an uncharacterised sequence from *C. caeruleus* in UniProt (UniProt id, A0A3N1FT86) with a putative 8-helix bundle, which again has KIH packing^46^ between pairs of helices (Supplementary Fig. 67).

The parallel designs all showed some similarity with natural and designed helical solenoid proteins (Fig. 5 and Supplementary Tables 38 – 43). This was anticipated because the h-t-h-t-h tertiary fragments used as connectors came from a set of *de novo* proteins of this type (Supplementary Table 4)^58^. Interestingly, searches with sc-CC-6, sc-CC-7, and sc-CC-8 consistently returned two hits: (1) the *de novo* circular Tandem Repeat Protein, cTRP9 (PDB id, 6xr1^66^), and (2) the putative inner membrane protein from *E. coli*, YhiM (UniProt id, P37630^38,63,67^, Fig. 5 and Supplementary Tables 38 – 43). This model, based on 5 central helices, has the most striking similarity to the parallel αHB proteins (Fig. 5); although we do not have an experimental structure for a *de novo* sc-CC-5 protein yet.

Recently, we expanded the CC+ Database of Coiled-coil Structures to include AlphaFold2 models of 48 proteomes^38,56,63^. Therefore, we searched these for potential single-chain antiparallel and parallel αHB proteins. This confirmed the YhiM and some similar proteins. However, it revealed no further examples of other higher-order antiparallel or parallel-based αHB proteins in PDB or AlphaFold2 databases. Socket2^46^ analysis of the KIH interactions in the top Foldseek^60^ hits revealed only 2- and 3-helix coiled-coil bundles—unlike the C*_n_* symmetric coiled-coil barrels with contiguous KIH interactions that we have targeted and made (Supplementary Fig. 65).

Together, these analyses indicate that the *de novo* αHB proteins that we present are a new class of single-chain coiled-coil protein. The designed proteins have accessible central channels that hit a sweet spot for small-molecule binding and, thus, are ripe for functionalisation^18,33–35^. They have been achieved through an accessible computational-design pipeline that combines rational-design principles and readily available computational design and modelling tools. This allowed us to arrive quickly at designed sequences for new coiled-coil-based proteins that surpass the complexity of natural or *de novo* coiled-coil structures reported to date. Moreover, this was achieved by testing a small number of gene constructs, and with high success rates (≈80%). The solution-phase characterization and high-resolution X-ray structures confirm our targets and, more importantly, our overall strategy of seeding computational design with established and understood rational-design rules. We envisage that the accessibility, versatility, and robustness of this approach will be of value to others in protein design, leading to applications in synthetic and cell biology, materials science, biotechnology, and other areas.

## Methods

### Computational tools

AlphaFold2 using single-sequence mode and 3 recycle steps was used to generate models for *de novo* peptide and protein designs. MASTER^36,37^ was used to build fragments (loops) between adjacent helices in the antiparallel and parallel αHB assemblies to connect the *C* and *N* termini of adjacent helices into single polypeptide chains. The Google colab notebook implementation of loop inpainting using RFDesign^9^ (https://github.com/polizzilab/design_tools) was used to generate short loop sequences (3 – 7 residues) to span between the different helices of the apCC-Hex backbone. ProteinMPNN^8^ was used to optimize the sequences of the MASTER loops for sc-apCC-8 and parallel protein designs. Additional details of scripts used for computational design from starting scaffold seeds are available in the Zenodo repository (https://doi.org/10.5281/zenodo.8277143) and Woolfson Lab github (https://github.com/woolfson-group).

### Peptide synthesis

Standard Fmoc automated microwave SPPS was performed on a 0.1 mmol scale using a Liberty Blue (CEM) synthesizer with inline UV monitoring. Activation was achieved with the coupling reagent *N’N’*-diisopropylcarbodiimid (DIC) in *N,N*-dimethylformamide (DMF) (1.0 mL, 1 M)/Oxyma Pure in DMF (1 mL, 0.5 M). Standard deprotections were performed using 20% (v/v) morpholine in DMF at 90 °C for 1 min (125 W 30 s, 32 W 60 s). All peptides were manually acetyl capped through addition of pyridine (0.5 mL) and acetic anhydride (0.25 mL) in DMF (9.25 mL), shaking at room temperature (rt) for 20 minutes. Peptides were cleaved from the resin with addition of 10 mL of a mixture 95:2.5:2.5 v/v trifluoroacetic acid (TFA)/H_2_O/triisopropylsilane (TIPS), shaking at room temperature for 2 hours. The TFA solution was then filtered to remove the resin beads and was reduced in volume to ≈5 mL or lower using a flow of N_2_. Cleaved peptide was precipitated with cold diethyl ether (≈45 mL), isolated via centrifugation and dissolved in a 1:1 mixture MeCN/H_2_O. Crude peptides were lyophilized to yield a white or off-white powder.

### Peptide purification

All peptides were purified by reverse phase HPLC (JASCO) using a Luna C18 (Phenomenex) column (150 × 10 mm, 5 μm particle size, 100 Å pore size). Crude peptide was injected to the column and eluted with a 3 mL/min linear gradient (40 – 100%) of MeCN in H_2_O with 0.1% TFA each over 30 minutes. Elution of the peptide was detected with inline UV monitoring at 220 and 280 nm wavelengths simultaneously. A column oven (50 °C) was employed to improve separation. Pure fractions were identified by analytical HPLC and matrix-assisted laser desorption/ionization–time of flight (MALDI-TOF) mass spectrometry. Analytical HPLC traces were obtained using a Jasco 2000 series HPLC system and a Phenomenex Kinetex C18 (100 × 4.6 mm, 5 μm particle size, 100 Å pore size) column. Chromatograms were monitored at 220 and 280 nm wavelengths. The linear gradient was 40 – 100% MeCN in water (each containing 0.1% TFA) over 25 min at a flow rate of 1 mL/min. When required, a column oven (50 °C) was employed to assist peptide elution. Matrix-assisted laser desorption/ionization–time of flight (MALDI-TOF) mass spectra were collected on a Bruker UltraFlex MALDI-TOF mass spectrometer operating in positive-ion reflector mode. Peptides were spotted on a ground steel target plate using α-cyano-4-hydroxycinnamic acid dissolved in 1:1 MeCN/H_2_O as the matrix. Masses quoted are for the monoisotopic mass as the singly protonated species.

### Protein expression and purification

All genes were directly cloned into pET28a vector, transformed then expressed in *E. coli* Lemo21-DE3 (NEB). Flasks containing 1 L of LB-kanamycin/chloramphenicol and 0.5 mM L-Rhamnose were inoculated with 5 mL overnight cultures and incubated to an OD_600_ of ≈0.6 at 37 °C with 200 rpm shaking. Expression was induced with 0.5 mM IPTG, and cultures were incubated at 37 °C overnight with 200 rpm shaking. Following expression, cultures were pelleted and resuspended in 20 mL lysis buffer (50 mM Tris, pH 7.4, 500 mM NaCl, 30 mM imidazole, 1 mg/mL lysozyme) for 30 min at 37 °C. Resuspended pellets were sonicated using a Biologics Model 3000 Ultrasonic homogenizer with settings to 50% power, 90% pulser (1 pulse/second) for five minutes, then clarified at 13,000 rpm for 30 minutes. The clarified lysate was heat-shocked at 75 °C for 10 minutes then cooled on ice for 10 minutes before reclarifying at 13,000 rpm for 10 minutes. The expressed proteins were first purified with Ni-affinity chromatography at room temperature. Filtered lysate was loaded onto an ÄKTAprime plus (GE) equipped with a HisTrap-5 mL HP column (Cytiva). His-tagged proteins were eluted using a single step gradient from 0 to 55% Buffer B (Buffer A: 50 mM Tris, 500 mM NaCl, 30 mM imidazole, pH 7.4; Buffer B: 50 mM Tris, 500 mM NaCl, 300 mM imidazole, pH 7.4). Fractions were combined and further purified by size exclusion chromatography using a HiLoad 16/600 Superdex 200 pg size exclusion column (Cytiva) equilibrated in buffer containing 50 mM sodium phosphate, pH 7.4, 150 mM NaCl at room temperature. Eluted fractions were pooled, concentrated, and run on SDS-PAGE to confirm identity.

### Circular dichroism (CD)

Circular dichroism (CD) data were collected on a JASCO J-810 or J-815 spectropolarimeter fitted with a Peltier temperature controller in far-UV region. Peptide samples were made up as 50 μM peptide solution in phosphate buffered saline (PBS; 8.2 mM sodium phosphate dibasic, 1.8 mM potassium phosphate monobasic, 137 mM NaCl, 2.4 mM KCl), pH 7.4 at 5 °C. For the antiparallel protein designs, CD spectra were acquired at 10 μM protein concentration in PBS at 5 °C. For the parallel protein designs, CD spectra were acquired at 5 μM protein concentration at 5 °C. Data were collected in a 1 mm quartz cuvette between 190 and 260 nm with the instrument set as follows: band width 1 nm, data pitch 1 nm, scanning speed 100 nm/min, 1 s response time. Each CD spectrum was obtained by averaging of 8 scans and subtracting the background signal of buffer and cuvette. For thermal-response experiments, the CD signal at 222 nm wavelength was monitored over the temperature range 5 – 95 °C at a ramp rate of 60 °C per hour and with the same settings and peptide or protein concentrations given above. The spectra were converted from ellipticities (mdeg) to mean residue ellipticities (MRE, (deg×cm^2^×dmol^-1^×res^-1^)) by normalizing for concentration of peptide bonds and the cell path length using the equation:

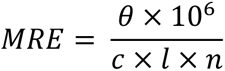

where the variable *θ* is the measured difference in absorbed circularly polarized light in millidegrees, *c* is the µM concentration of the compound, *l* is the path length of the cuvette in mm, and *n* is the number of amide bonds in the polypeptide.

### Analytical ultracentrifugation

Analytical ultracentrifugation (AUC) was performed on a Beckman Optima X-LA or X-LI analytical ultracentrifuge with an An-50-Ti or An-60-Ti rotor (Beckman-Coulter). Buffer densities, viscosities and peptide and protein partial specific volumes (^v̅^) were calculated using SEDNTERP (http://rasmb.org/sednterp/). For sedimentation velocity (SV), peptide sample were prepared in PBS at 150 μM peptide concentration and placed in a sedimentation velocity cell with 2-channel centrepiece and quartz windows. The samples were centrifuged at 50 krpm at 20 °C, with absorbance scans taken over a radial range of 5.8 – 7.3 cm at 5 min intervals to a total of 120 scans. For SV experiments with the antiparallel designs, samples were prepared at 15 μM protein concentration in PBS. The samples were centrifuged at 50 krpm (40 krpm for sc-apCC-8) using the same method as the peptide experiments. For SV experiments with the parallel designs, samples were prepared at 25 μM protein concentration in PBS. The samples were centrifuged at 40 or 50 krpm using the same method as above samples. Data from a single run were fitted to a continuous c(s) distribution model using SEDFIT^68^ at 95% confidence level. Residuals for sedimentation velocity experiments are shown as a bitmap in which the grayscale shade indicates the difference between the fit and raw data (residuals < -0.05 black, > 0.05 white). Good fits are uniformly grey without major dark or light streaks. Sedimentation equilibrium (SE) experiments were performed at 70 μM peptide concentration in 110 μL at 20 °C. The experiment was run in triplicate in a six-channel centrepiece. The samples were centrifuged at speeds in the range of or 20 – 45 krpm and scans at each recorded speed were duplicated after equilibration for 8 hours. Data were fitted using SEDPHAT^69^ to a single species model. Monte Carlo analysis was performed to give 95% confidence limits.

### Ligand binding

Ligand-binding experiments were pipetted in quadruplicate using an epMotion 5070 liquid handler (Eppendorf). The total concentration of ligand was kept constant (1 μM DPH in 5% v/v DMSO) and the concentration of *de novo* peptide assembly and antiparallel protein design varied from 0 – 30 μM. For parallel designs, ligand concentration was kept constant at 0.5 μM and the protein concentration was varied from 0 – 24 μM. Data were collected on a Clariostar plate reader (BMG Labtech) using an excitation wavelength of 350 nm and emission monitored at 450 nm. Binding constants were extracted by fitting the data to the following equation:

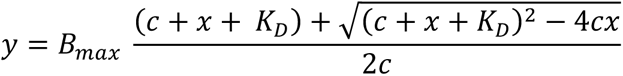

where c is the total concentration of the constant component (*e.g.* DPH), × is the concentration of variable component (*e.g.* peptide/protein), B_max_ is the fluorescence signal when all of the constant component is bound, and y is the fraction of bound component being monitored via fluorescence signal.

### Size exclusion chromatography small angle X-ray scattering (SEC-SAXS)

Data for single-chain protein designs were obtained at the Diamond Light Source on beamline B21. Samples were prepared to 10 mg/mL in 50 mM sodium phosphate, pH 7.4, 150 mM NaCl. A Superdex 200 Increase 3.2/300 was equilibrated in the same buffer at 4 °C. Buffer subtraction and data merging was performed with ScÅtter^70^. q_min_ was taken as the first point of the linear Guinier region, q_max_ was calculated using ShaNum through ATSAS interface^71^. MultiFoxS software (Sali Lab) using a monomer model was used to compare experimental scattering profiles to design models and assess quality of fit by calculating *X*^2 47,48^.

### X-ray crystallography

Diffraction-quality peptide crystals were grown using a sitting-drop vapour-diffusion method. Commercially available sparse matrix screens were used (Morpheus®, JCSG-plus^TM^, Structure Screen 1 and 2, Pact Premier^TM^, ProPlex^TM^; Molecular Dimensions), and the drops were dispensed using a robot (Oryx8; Douglas Instruments). For each well of an MRC 2 drop plate, 0.3 μL of peptide or protein solution and 0.3 μL of reservoir solution in parallel with 0.4 μL of the peptide or protein solution and 0.2 μL of reservoir solution were mixed and the plate was incubated at 20 °C. Crystals of antiparallel and parallel protein designs were obtained by optimization by seeding and cross seeding. Crystals were mounted and transferred into a cryogenic solution made of the corresponding reservoir solution supplemented with 25% glycerol, and flash-cooled in liquid nitrogen. Diffraction data for the crystals were obtained at the Diamond Light Source (Didcot, UK) on beamlines I04 or I24. Data were processed using the automated pipelines: Xia2 pipelines^72^, which ports data through DIALS^73^ or MOSFLM^74^ to POINTLESS and AIMLESS^75^ as implemented in the CCP4 suite^76^, or XDS to XSCALE^77^; or the AUTOPROC pipelines, which use the same integrating and data reduction software in addition to STARANISO^78^. The data were phased using either *ab initio* phasing using ARCIMBOLDO_LITE^79,80^ or molecular replacement using an AlphaFold2 model for PHASER^81^. Final structures were obtained after iterative rounds of model building with COOT^82^ and refinement with REFMAC5^83^ and Phenix Refine^84^.

## Data availability

The MASTER^36,37^, CC+^38^ and Foldseek^60^ databases are open and publicly accessible. The coordinate and structure factor files for g-a-d-e = ALIA, g-a-d-e = GLIA, apCC-Hex, sc-apCC-6-LLIA, sc-apCC-6-SLLA, sc-apCC-8, sc-CC-6-95, sc-CC-7-LI, and sc-CC-8-58 have been deposited in the Protein Data Bank with accession codes 8qaa, 8qac, 8qab, 8qad, 8qae, 8qaf, 8qag, 8qai, and 8qah respectively. The raw data and code used in this publication has been deposited in the Zenodo repository (https://doi.org/10.5281/zenodo.8277143) and Woolfson Lab github (https://github.com/woolfson-group).

## Acknowledgements

K.I.A., O.D.W. and D.N.W. are supported by a BBSRC-NSF grant (BB/V004220/1 and 2019598). We are also grateful to the Max Planck-Bristol Centre for Minimal Biology, which supports K.I.A. and D.N.W.. R.P. is supported by a BBSRC-funded PhD studentship and by Rosa Biotech (SWBio DTP). F.P. was supported by an EPSRC programme grant to G.J.G and D.N.W. (EP/T012455/1). E.A.N. was supported by a BBSRC grant to Nigel Savery and D.N.W. (BB/S002820/1). O.D.W. is grateful for a National Institutes of Health grant (GM-118167). We thank the University of Bristol, School of Chemistry, Mass Spectrometry Facility for access to the EPSRC-funded Bruker Ultraflex MALDI-TOF instrument (EP/K03927X/1). We would like to thank Diamond Light Source for access to beamlines I04, 24, and B21 (Proposals mx23269 and mx31440). Finally, we thank Nigel Savery for advice and support in the early stages of our peptides-to-proteins programme, and Joel Chubb, Kelsey Kean, Bram Mylemans, and members of the Woolfson laboratory for many helpful discussions.

## Author contributions

K.I.A. and R.P contributed equally. K.I.A., R.P., F.P., W.M.D., and D.N.W. conceived the study and contributed to experimental design. K.I.A. designed and characterised the individual peptides. E.A.N. characterised the ap-CC-Hex peptide and determined the crystal structure. K.I.A. designed and characterised the antiparallel αHB proteins. R.P. designed and characterised the parallel αHB proteins. K.I.A. and R.P. collected the X-ray data and solved the crystal structures. R.P. collected the SEC-SAXS data and K.I.A. and R.P. analysed the data. With D.N.W., D.A.S, G.J.L, O.D.W., and T.A.A.O provided supervision/mentorship for these researchers. K.I.A., R.P., and D.N.W. wrote the paper. All authors have read and contributed to the preparation of the manuscript.

