## Supplementary for "Rationally seeded computational protein design"

### 1 Methods

#### 1.1 General

All solvents, chemicals, and reagents were purchased from commercial sources and used without further purification. Fluorenylmethoxycarbonyl(Fmoc)-α-L-amino acids, Rink amide MBHA resin for solid-phase peptide synthesis (SPPS) and N,N-dimethylformamide (DMF) were purchased from Sigma-Aldrich and Cambridge Reagents. Coupling reagents Oxyma Pure and diisopropylcarbodiimide (DIC) were purchased from Fluorochem. Morpholine, trifluoroacetic acid (TFA) were purchased from Sigma-Aldrich; pyridine was from Thermo Fisher; triisopropylsilane (TIPS) was from Acros Organics. Luria-Broth (LB), antibiotics, IPTG, and L-rhamnose were purchased from Thermo Fisher. All other chemicals were reagent grade and purchased from Sigma-Aldrich. Peptide biophysical characterization were recorded in phosphate buffered saline (PBS; 8.2 mM sodium phosphate dibasic, 1.8 mM potassium phosphate monobasic, 137 mM NaCl, 2.4 mM KCl), pH 7.4. Protein biophysical characterization were recorded in an adjusted PBS buffer (50 mM sodium phosphate, 150 mM NaCl, pH 7.4). For conciseness, both buffers are referred to as PBS in the manuscript. Peptide characterisation data for CC-Type2-$(A_gL_aI_d)_4$, CC-Type2-$(A_gI_aI_d)_4$, CC-Type2-$(A_gI_aV_d)_4$, CC-Type2-$(T_gL_aI_d)_4$, and CC-Type2-EEKK-$(L_gL_aI_d)_4$ (apCCHex-LLIA) have been published previously[1-4].

#### 1.2 Computational tools

##### 1.2.1 AlphaFold2 prediction

AlphaFold2[5,6] using single-sequence mode and 3 recycle steps was used to assess the antiparallel sc-apCC-6, sc-apCC-8 and parallel sc-CC-5, sc-CC-6, sc-CC-7, sc-CC-8 top 5 models. Predicted local-distance difference test (pLDDT) per position, predicted template modelling score (pTM), and the predicted alignment error (pAE) were collected to evaluate the models. In particular, pTM and average pLDDT were evaluated for the single-chain proteins. AlphaFold-Multimer[7] predictions were performed using the ColabFold (version 1.3.0) using single-sequence mode and 3 recycle steps to return the top 5 ranked models for sequences used to seed the sc-apCC-8 designs.

##### 1.2.2 MASTER

MASTER[8,9] was used to build fragments (loops) between adjacent helices in the antiparallel and parallel αHB assemblies to connect the *C* and *N* termini of adjacent helices into single polypeptide chains. The built-in MASTER database was used and the following parameters were set to find matches to the antiparallel queries (the first and last heptad of each chain): rmsdcut = 0.5 and wgap = '3 – 10'. This was the final step for sc-apCC-6 MASTER loop designs. A dataset of 3-helix coiled-coil bundles was generated using CC+[10,11] (50% sequence identity, 7 Å cutoff, 1059 bundles in total) and used for loop queries for the parallel helical assemblies (using the last and the first heptad of each peptide chain). Parameters: rmsdcut = 0.7 and wgap = '30 – 100'.

##### 1.2.3 ColabPaint: Inpainting using RFDesign

A third approach to designing loops was used for the sc-apCC-6 target. The Google colab notebook implementation of loop inpainting using RFDesign[12] written by Nick Polizzi (https://github.com/polizzilab/design_tools) was used to generate short loop sequences (3 – 7 residues) to span between the different helices of the apCC-Hex backbone. Two different sequences were returned and evaluated using AlphaFold2 as described above in 1.2.1.

##### 1.2.4 sc-apCC-8 loop design

First, MASTER[8,9] (same protocol as 1.2.2 used for sc-apCC-6) was used to build fragments to connect the *C* and *N* termini of adjacent helices into single polypeptide chains for AlphaFold2 models of apCC-Oct (g-a-d-e = Ala-Ile-Ile-Ala, Gly-Ile-Ile-Ala, Gly-Leu-Leu-Ala, and g-$(a\text{-}d)_2(a\text{-}d)_2$-e (the two a-d combinations are repeated through the first two and last two heptads respectively) = Gly-$(\text{Ile-Leu})_2(\text{Leu-Ile})_2$-Ala and Gly-$(\text{Leu-Ile})_2(\text{Ile-Leu})_2$-Ala. Next, all residues of the barrel scaffold (gabcdef) were fixed and ProteinMPNN[13] (T = 0.1, 10 sequences) was used to optimize the sequences of the MASTER loops for sc-apCC-8 sequences. The 10 sequences were then run through AlphaFold2 and pTM and average pLDDT for all models were evaluated. g-a-d-e = Gly-Ile-Ile-Ala and Gly-Leu-Leu-Ala had lower pTM and pLDDT scores and not all models were folded as open, meandering up-down-up αHB proteins, thus were not considered.

We then selected a single sequence with the highest pLDDT and pTM for both g-a-d-e = Ala-Ile-Ile-Ala and Gly-(Ile-Leu)$_2$(Leu-Ile)$_2$-Ala to carry forward for characterization.

##### 1.2.5 Iterative parallel variant design

After MASTER, ProteinMPNN (T = 0.2, Cys and Met omitted, 100 sequences) was used to design the sequences for sc-CC-6, sc-CC-7 and sc-CC-8. The core g-a-d-e positions were fixed to Ala-Leu-Ile-Ala for sc-CC-6 and sc-CC-7, and Ala-Ile-Ile-Ala for sc-CC-8 throughout the design process. For the first round of sequence designs, the b-c-f positions on external helices were fixed to Lys-Glu-Gln to achieve a native pI distribution. The designed sequences were repacked onto the starting backbone, relaxed with Rosetta[14,15] and filtered with the following filters: *pI < 9; Rosetta packing statistic > mean of all sequences; Rosetta energy score < mean of all sequences*. Filtered sequences were run through AlphaFold2 (single-sequence mode), and the model with the best max pLDDT score was used for the next round of sequence design with ProteinMPNN.

For further design rounds, Lys-Glu-Gln surface constraint was changed into Lys-Glu amino acid bias (K = –0.35, E = 0.35 for sc-CC-7, K = –0.4, E = 0.4 for sc-CC-6 and sc-CC-8) as this gave the desired pI distribution but reduced the sequence repetitiveness. An additional exposed hydrophobics score filter (< mean of all sequences) was also added to minimize the exposure of hydrophobic amino acids on the surface of the expressed proteins. Even with the hydrophobics filter, the b-c-f positions of the *N*- and *C*-terminal helices of sc-CC-8 had to be fixed to Lys-Glu-Gln to reduce the exposure of hydrophobic amino acids.

The design process was repeated until the Rosetta energy score and the RMSD between the ProteinMPNN input and the corresponding AlphaFold2 output converged. This took 2 design rounds for sc-CC-6 and 3 rounds for sc-CC-7 and sc-CC-8. The resulting designs were filtered with 85% sequence identity, visually inspected and 4 final designs were picked based on Rosetta energy score and max AlphaFold2 pLDDT.

#### 1.3 Peptide synthesis and purification

##### 1.3.1 Automated microwave Fmoc/tBu solid-phase peptide synthesis (SPPS)

Automated microwave SPPS was performed on a Liberty Blue (CEM) synthesizer with inline UV monitoring. Syntheses were performed on 0.1 mmol scales with side-chain protection of the amino acids as follows: Gln(Trt), Glu(O*t*Bu), Lys(Boc), Tyr(*t*Bu), Trp(Boc). The resin (Rink amide MBHA, 0.65 mmol/g loading, 100 – 200 mesh) was weighted to enable a synthesis on a 0.1 mmol scale. The coupling reactions were performed by adding protected amino acids dissolved in DMF (2.5 mL, 0.2 M), the coupling reagent DIC in DMF (1.0 mL, 1 M) and Oxyma Pure in DMF (1 mL, 0.5 M) to the respective resin. Standard couplings were performed at 90 °C for 4 min (100 W for 20 s, 60 W for 10 s, 35 W for 240 s). Standard deprotections were performed using 20% (v/v) morpholine in DMF at 90 °C for 1 min (125 W 30 s, 32 W 60 s). All peptides were manually acetyl capped through addition of pyridine (0.5 mL) and acetic anhydride (0.25 mL) in DMF (9.25 mL), shaking at room temperature (rt) for 20 minutes. The resin was washed three time with DMF followed by six times with DCM before cleavage. Peptides were cleaved from the resin with addition of 10 mL of a mixture 95:2.5:2.5 v/v trifluoroacetic acid (TFA)/$H_2O$/triisopropylsilane (TIPS), shaking at room temperature for 2 hours. The TFA solution was then filtered to remove the resin beads and was reduced in volume to ≈5 mL or lower using a flow of $N_2$. Cleaved peptide was precipitated with cold diethyl ether (≈45 mL), isolated via centrifugation and dissolved in a 1:1 mixture MeCN/$H_2O$. Crude peptides were lyophilized to yield a white or off-white powder.

##### 1.3.2 Semi-preparative High Performance Liquid Chromatography (HPLC)

All peptides were purified by reverse phase HPLC (JASCO) using a Luna C18 (Phenomenex) column (150 x 10 mm, 5 µm particle size, 100 Å pore size). Crude peptide was dissolved at 7 mg/mL in 40% v/v MeCN in $H_2O$ with 0.1% TFA, injected to the column and eluted with a 3 mL/min linear gradient (40 – 100%) of MeCN in $H_2O$ with 0.1% TFA each over 30 minutes. Elution of the peptide was detected with inline UV monitoring at 220 and 280 nm wavelengths simultaneously. A column oven (50 °C) was employed to improve separation. Pure fractions were identified by analytical HPLC and matrix-assisted laser desorption/ionization–time of flight (MALDI-TOF) mass spectrometry, then pooled, and freeze-dried.

##### 1.3.3 Analytical HPLC

Analytical HPLC traces were obtained using a Jasco 2000 series HPLC system and a Phenomenex Kinetex C18 (100 x 4.6 mm, 5 µm particle size, 100 Å pore size) column. Chromatograms were monitored at 220 and 280 nm wavelengths. The linear gradient was 40 – 100% MeCN in water (each containing 0.1% TFA) over 25 min at a flow rate of 1 mL/min. When required, a column oven (50 °C) was employed to assist peptide elution.

##### 1.3.4 Mass spectrometry

Matrix-assisted laser desorption/ionization–time of flight (MALDI-TOF) mass spectra were collected on a Bruker UltraFlex MALDI-TOF mass spectrometer operating in positive-ion reflector mode. Peptides were spotted on a ground steel target plate using α-cyano-4-hydroxycinnamic acid dissolved in 1:1 MeCN/$H_2O$ as the matrix. Masses quoted are for the monoisotopic mass as the singly protonated species.

#### 1.4 Protein expression and purification

##### 1.4.1 Cloning and protein sequence

All genes were directly cloned into pET28a vector using NdeI and XhoI restriction sites and purchased from Twist Biosciences or GeneArt from Fisher Scientific.

##### 1.4.2 Protein expression and purification

All genes were transformed then expressed in *E. coli* Lemo21-DE3 (NEB). Flasks containing 1 L of LB-kanamycin/chloramphenicol and 0.5 mM L-Rhamnose were inoculated with 5 mL overnight cultures and incubated to an $OD_{600}$ of ≈0.6 at 37 ˚C with 200 rpm shaking. Expression was induced with 0.5 mM IPTG, and cultures were incubated at 37 ˚C overnight with 200 rpm shaking.

Following expression, cultures were pelleted at 4000 rpm for 10 minutes. Cell pellets were resuspended in 20 mL lysis buffer (50 mM Tris, pH 7.4, 500 mM NaCl, 30 mM imidazole, 1 mg/mL lysozyme) for 30 min at 37 ˚C. Resuspended pellets were sonicated using a Biologics Model 3000 Ultrasonic homogenizer with settings to 50% power, 90%

pulser (1 pulse/second) for five minutes, then clarified at 13,000 rpm for 30 minutes. The clarified lysate was heat-shocked at 75 ˚C for 10 minutes then cooled on ice for 10 minutes before reclarifying at 13,000 rpm for 10 minutes. The expressed proteins were first purified with Ni-affinity chromatography at room temperature. Filtered lysate was loaded onto an ÄKTAprime plus (GE) equipped with a HisTrap-5 mL HP column (Cytiva). His-tagged proteins were eluted using a single step gradient from 0 to 55% Buffer B (Buffer A: 50 mM Tris, 500 mM NaCl, 30 mM imidazole, pH 7.4; Buffer B: 50 mM Tris, 500 mM NaCl, 300 mM imidazole, pH 7.4). Fractions were combined and further purified by size exclusion chromatography using a HiLoad 16/600 Superdex 200 pg size exclusion column (Cytiva) equilibrated in buffer containing 50 mM sodium phosphate, pH 7.4, 150 mM NaCl at room temperature. Eluted fractions were pooled, concentrated, and run on SDS-PAGE to confirm identity. The parallel designs were purified following the same protocol and lower concentration (150 mM NaCl) salt buffers from above for the Ni-affinity chromatography.

#### 1.5 Solution-phase biophysical characterizations

##### 1.5.1 Peptide and protein concentration determination

Peptide and protein concentration was determined at 280 nm using a Nanodrop 2000 (ThermoScientific) spectrometer ($\varepsilon_{280}$(Trp) = 5690 $M^{-1}cm^{-1}$; $\varepsilon_{280}$ (proteins), see Supplementary Tables 5, 10, 15, 20, 22, 24).

##### 1.5.2 Circular dichroism (CD) spectroscopy

Circular dichroism (CD) data were collected on a JASCO J-810 or J-815 spectropolarimeter fitted with a Peltier temperature controller in far-UV region. Peptide samples were made up as 50 µM peptide solution in phosphate buffered saline (PBS; 8.2 mM sodium phosphate dibasic, 1.8 mM potassium phosphate monobasic, 137 mM NaCl, 2.4 mM KCl), pH 7.4 at 5 °C. For the antiparallel protein designs, CD spectra were acquired at 10 µM protein concentration in 50 mM sodium phosphate, pH 7.4, 150 mM NaCl at 5 °C. For the parallel protein designs, CD spectra were acquired at 5 µM protein concentration at 5 °C. Data were collected in a 1 mm quartz cuvette between 190 and 260 nm with the instrument set as follows: band width 1 nm, data pitch 1 nm, scanning speed 100 nm/min, 1 s response time. Each CD spectrum was obtained by averaging of 8 scans and subtracting the background signal of buffer and cuvette.

For thermal-response experiments, the CD signal at 222 nm wavelength was monitored over the temperature range 5 – 95 °C at a ramp rate of 60 °C per hour and with the same settings and peptide or protein concentrations given above.

The spectra were converted from ellipticities (mdeg) to mean residue ellipticities (MRE, ($deg \cdot cm^2 \cdot dmol^{-1} \cdot res^{-1}$)) by normalizing for concentration of peptide bonds and the cell path length using the equation:

$$MRE = \frac{\theta \times 10^6}{c \times l \times n}$$

where the variable $\theta$ is the measured difference in absorbed circularly polarized light in millidegrees, $c$ is the µM concentration of the compound, $l$ is the path length of the cuvette in mm, and $n$ is the number of amide bonds in the polypeptide.

##### 1.5.3 Analytical ultracentrifugation

Analytical ultracentrifugation (AUC) was performed on a Beckman Optima X-LA or X-LI analytical ultracentrifuge with an An-50-Ti or An-60-Ti rotor (Beckman-Coulter). Buffer densities, viscosities and peptide and protein partial specific volumes ($\overline{\nu}$) were calculated using SEDNTERP (http://rasmb.org/sednterp/).

For sedimentation velocity (SV), peptide sample solutions of 310 or 410 µL were prepared in PBS at 150 µM peptide concentration and placed in a sedimentation velocity cell with an epon or aluminium, respectively, 2-channel centrepiece and quartz windows. The reference channel was loaded with 320 µL (or 420 µL, respectively) of PBS buffer. The samples were centrifuged at 50 krpm at 20 °C, with absorbance scans taken over a radial range of 5.8 – 7.3 cm at 5 min intervals to a total of 120 scans. For SV experiments with the antiparallel designs, samples were prepared at 15 µM protein concentration in 50 mM sodium phosphate, pH 7.4, 150 mM NaCl. The samples were centrifuged at 50 krpm (40 krpm for sc-apCC-8) using the same method as the peptide experiments. For SV experiments with the parallel designs, samples were prepared at 25 µM protein concentration in 50 mM sodium phosphate, pH 7.4, 150 mM NaCl. The samples were centrifuged at 40 or 50 krpm using the same method as above samples. Data from a single run were fitted to a continuous c(s) distribution model using SEDFIT[16] at 95% confidence level. Residuals for sedimentation velocity experiments are shown as a bitmap in which the grayscale shade indicates the difference between the fit and raw data (residuals < -0.05 black, > 0.05 white). Good fits are uniformly grey without major dark or light streaks.

Sedimentation equilibrium (SE) experiments were performed at 70 μM peptide concentration in 110 μL at 20 °C. The experiment was run in triplicate in a six- channel centrepiece. The samples were centrifuged at speeds in the range of or 20 – 45 krpm and scans at each recorded speed were duplicated after equilibration for 8 hours. Data were fitted using SEDPHAT[17] to a single species model. Monte Carlo analysis was performed to give 95% confidence limits.

##### 1.5.4 Ligand binding

Ligand-binding experiments were pipetted in quadruplicate using an epMotion 5070 liquid handler (Eppendorf). The total concentration of ligand was kept constant (1 μM DPH in 5% v/v DMSO) and the concentration of *de novo* peptide assembly and antiparallel protein design varied from 0 – 30 μM. For parallel designs, ligand concentration was kept constant at 0.5 μM and the protein concentration was varied from 0 – 24 μM. Data were collected on a Clariostar plate reader (BMG Labtech) using an excitation wavelength of 350 nm and emission monitored at 450 nm. Binding constants were extracted by fitting the data to the following equation:

$$y = B_{max} \frac{(c + x + K_D) + \sqrt{(c + x + K_D)^2 - 4cx}}{2c}$$

where c is the total concentration of the constant component (*e.g.* DPH), x is the concentration of variable component (*e.g.* peptide/protein), $B_{max}$ is the fluorescence signal when all of the constant component is bound, and y is the fraction of bound component being monitored via fluorescence signal.

#### 1.6 Structural characterization

##### 1.6.1 Size exclusion chromatography small angle X-ray scattering (SEC-SAXS)

Data for single-chain protein designs were obtained at the Diamond Light Source on beamline B21. Samples were prepared to 10 mg/mL in 50 mM sodium phosphate, pH 7.4, 150 mM NaCl. A Superdex 200 Increase 3.2/300 was equilibrated in the same buffer at 4 °C. Buffer subtraction and data merging was performed with ScÅtter[18]. $q_{min}$ was taken as the first point of the linear Guinier region, $q_{max}$ was calculated using ShaNum through ATSAS interface[19]. MultiFoxS software (Sali Lab) was used to compare experimental scattering profiles to design models and assess quality of fit by calculating $\chi^2$ [20,21]. The data were fit to His-tagged models of the single-chain proteins.

##### 1.6.2 Crystals growth

Diffraction-quality peptide crystals were grown using a sitting-drop vapour-diffusion method. Freeze-dried peptides were dissolved in ultrapure water and diluted to 10 mg/mL. Purified antiparallel protein designs in 20 mM Tris pH 8.0, 50 mM NaCl were concentrated to 10 mg/mL. Commercially available sparse matrix screens were used (Morpheus®, JCSG-plus™, Structure Screen 1 and 2, Pact Premier™, ProPlex™; Molecular Dimensions), and the drops were dispensed using a robot (Oryx8; Douglas Instruments). For each well of an MRC 2 drop plate, 0.3 µL of peptide or protein solution and 0.3 µL of reservoir solution in parallel with 0.4 µL of the peptide or protein solution and 0.2 µL of reservoir solution were mixed and the plate was incubated at 20 °C. Crystals generally formed within a month, and after looping were soaked in reservoir solution containing 25% glycerol as a cryoprotectant. Crystals of antiparallel and parallel protein designs were obtained by optimization by seeding and cross seeding. For seeding experiments, 6 – 8 mg/mL of protein solution were used and for each well of an MRC 2 drop plate, 0.3 µL of peptide or protein solution, 0.1µL of seed, and 0.2 µL of reservoir solution in parallel with 0.4 µL of the peptide or protein solution, 0.1 µL of seed, and 0.1 µL of reservoir solution were mixed and the plate was incubated at 20 °C.

Final crystallization conditions for all peptides and proteins are provided in Supplementary Table 29.

##### 1.6.3 X-ray crystal structure determination

Diffraction data for the crystals were obtained at the Diamond Light Source (Didcot, UK) on beamlines I04 or I24. Data were processed using the automated pipelines: Xia2 pipelines[22], which ports data through DIALS[23] or MOSFLM[24] to POINTLESS and AIMLESS[25] as implemented in the CCP4 suite[26], or XDS to XSCALE[27]; or the AUTOPROC pipelines, which use the same integrating and data reduction software in addition to STARANISO[28]. apCC-Hex, apCC-Hex-ALIA collapsed bundle, apCC-Oct-GLIA collapsed bundle, sc-apCC-6-LLIA, and sc-apCC-8-AIIA were phased using *ab initio* phasing using ARCIMBOLDO_LITE[29,30]. The initial phases were modelled into and refined using BUCCANEER[31]. sc-apCC-6-SLLA was solved by molecular replacement using the AlphaFold2 model for PHASER[32]. Final structures were obtained after iterative rounds of model building with COOT[33] and refinement with REFMAC5[34] and Phenix Refine[35]. Solvent-exposed atoms lacking map density were either deleted or left at full occupancy. When applicable, symmetry mates were generated using PISA[26,36]. Data collection and refinement statistics are provided in Supplementary Table 30.

### 2 Supplementary data

#### 2.1 Supplementary tables

##### 2.1.1 *De novo* peptide sequences and characterization

**Supplementary Table 1. Sequences of *de novo* peptides characterized in this study.**

| Systematic name | *gade* | Sequences<br>*gabcdefgabcdefgabcdefgabcdef* | Mass (g/mol) |
|---|---|---|---|
| CC-Type2-EEKK-($A_gI_aI_dA_e$)$_4$ | AIIA | Ac–G AIEEIAQAIEEIAKAIKKIAWAIKKIAQ G–$NH_2$ | 3246.84 |
| CC-Type2-EEKK-($A_gI_aL_dA_e$)$_4$ | AILA | Ac–G AIEELAQAIEELAKAIKKLAWAIKKLAQ G–$NH_2$ | 3246.84 |
| CC-Type2-EEKK-($A_gL_aI_dA_e$)$_4$ | ALIA | Ac–G ALEEIAQALEEIAKALKKIAWALKKIAQ G–$NH_2$ | 3246.84 |
| CC-Type2-EEKK-($A_gL_aL_dA_e$)$_4$ | ALLA | Ac–G ALEELAQALEELAKALKKLAWALKKLAQ G–$NH_2$ | 3246.84 |
| CC-Type2-EEKK-($G_gI_aI_dA_e$)$_4$ | GIIA | Ac–G GIEEIAQGIEEIAKGIKKIAWGIKKIAQ G–NH2 | 3190.70 |
| CC-Type2-EEKK-($G_gI_aL_dA_e$)$_4$ | GILA | Ac–G GIEELAQGIEELAKGIKKLAWGIKKLAQ G–NH2 | 3190.70 |
| CC-Type2-EEKK ($G_gI_aL_dA_e$)$_2$/($G_gL_aI_dA_e$)$_2$ | G(IL)$_2$(LI)$_2$A | Ac–G GIEELAQGIEELAKGLKKIAWGLKKIAQ G–NH2 | 3190.70 |
| CC-Type2-EEKK-($G_gL_aI_dA_e$)$_4$ | GLIA | Ac–G GLEEIAQGLEEIAKGLKKIAWGLKKIAQ G–NH2 | 3190.70 |
| CC-Type2-EEKK-($G_gL_aI_dA_e$)$_2$/($G_gI_aL_dA_e$)$_2$ | G(LI)$_2$(IL)$_2$A | Ac–G GLEEIAQGLEEIAKGIKKLAWGIKKLAQ G–NH2 | 3190.70 |
| CC-Type2-EEKK-($G_gL_aL_dA_e$)$_4$ | GLLA | Ac–G GLEELAQGLEELAKGLKKLAWGLKKLAQ G–NH2 | 3190.70 |
| CC-Type2-EEKK-($L_gI_aI_dA_e$)$_4$ | LIIA | Ac–G LIEEIAQLIEEIAKLIKKIAWLIKKIAQ G–$NH_2$ | 3415.16 |
| CC-Type2-EEKK-($L_gI_aL_dA_e$)$_4$ | LILA | Ac–G LIEELAQLIEELAKLIKKLAWLIKKLAQ G–$NH_2$ | 3415.16 |
| CC-Type2-EEKK-($L_gL_aI_dA_e$)$_4$ apCC-Hex[4] | LLIA | Ac–G LLEEIAQLLEEIAKLLKKIAWLLKKIAQ G–$NH_2$ | 3415.16 |
| CC-Type2-EEKK-($L_gL_aL_dA_e$)$_4$ | LLLA | Ac–G LLEELAQLLEELAKLLKKLAWLLKKLAQ G–$NH_2$ | 3415.16 |
| CC-Type2-EEKK-($M_gI_aI_dA_e$)$_4$ | MIIA | Ac–G MIEEIAQMIEEIAKMIKKIAWMIKKIAQ G–$NH_2$ | 3487.32 |
| CC-Type2-EEKK-($M_gI_aL_dA_e$)$_4$ | MILA | Ac–G MIEELAQMIEELAKMIKKLAWMIKKLAQ G–$NH_2$ | 3487.32 |
| CC-Type2-EEKK-($M_gL_aI_dA_e$)$_4$ | MLIA | Ac–G MLEEIAQMLEEIAKMLKKIAWMLKKIAQ G–$NH_2$ | 3487.32 |
| CC-Type2-EEKK-($M_gL_aL_dA_e$)$_4$ | MLLA | Ac–G MLEELAQMLEELAKMLKKLAWMLKKLAQ G–$NH_2$ | 3487.32 |
| CC-Type2-EEKK-($S_gI_aI_dA_e$)$_4$ | SIIA | Ac–G SIEEIAQSIEEIAKSIKKIAWSIKKIAQ G–$NH_2$ | 3310.84 |
| CC-Type2-EEKK-($S_gI_aL_dA_e$)$_4$ | SILA | Ac–G SIEELAQSIEELAKSIKKLAWSIKKLAQ G–$NH_2$ | 3310.84 |
| CC-Type2-EEKK-($S_gL_aI_dA_e$)$_4$ | SLIA | Ac–G SLEEIAQSLEEIAKSLKKIAWSLKKIAQ G–$NH_2$ | 3310.84 |
| CC-Type2-EEKK-($S_gL_aL_dA_e$)$_4$ | SLLA | Ac–G SLEELAQSLEELAKSLKKLAWSLKKLAQ G–$NH_2$ | 3310.84 |

**Supplementary Table 2. Sedimentation velocity AUC fitting statistics for the *de novo* peptides characterized in this study.**

| ***gade*** | **Mass** (g/mol) | **$\bar{v}^{1}$** (cm$^3$ g$^{-1}$) | **Fitted Mass SV**[2] (95% confidence, 3 SF) | **f/f0**[3] | **s**[4] (S) | **s20,w**[5] (S) | **SV weight/ monomer mass** | **Fitted Mass SE**[2] **(lower, upper)** (95% confidence, 3 SF) | **SE weight/ monomer mass** |
|---|---|---|---|---|---|---|---|---|---|
| AIIA | 3246.84 | 0.779 | 22400 | 1.189 | 1.886 | 1.935 | 6.9 | 19100 (18800, 19400) | 5.9 |
| ALIA | 3246.84 | 0.779 | 19300 | 1.171 | 1.733 | 1.778 | 5.9 | 16300 (15800, 16700) | 5.0 |
| ALLA | 3246.84 | 0.779 | 20200 | 1.2 | 1.745 | 1.79 | 6.2 | 18400 (18300, 18500) | 5.7 |
| GIIA | 3190.7 | 0.77 | 19000 | 1.09 | 1.929 | 1.976 | 6.0 | 20900 (20700, 21200) | 6.6 |
| G(IL)$_2$(LI)$_2$A | 3190.7 | 0.77 | 20200 | 1.126 | 1.945 | 1.993 | 6.3 | 20900 (20800, 21200) | 6.6 |
| GLIA | 3190.7 | 0.77 | 19200 | 1.133 | 1.866 | 1.911 | 6.0 | 20900 (20800, 21100) | 6.6 |
| G(LI)$_2$(IL)$_2$A | 3190.7 | 0.77 | 20900 | 1.133 | 1.979 | 2.027 | 6.6 | 21000 (20800, 21200) | 6.6 |
| GLLA | 3190.7 | 0.77 | 16900 | 1.133 | 1.716 | 1.757 | 5.3 | 21100 (20300, 21800) | 6.6 |
| LIIA* | 3415.16 | 0.799 | 22700 | 1.198 | 1.698 | 1.747 | 6.6 | 21900 (21600, 22400) | 6.4 |
| LLIA | 3415.16 | 0.799 | 18700 | 1.168 | 1.592 | 1.648 | 5.5 | 20100 (19900, 20300) | 5.9 |
| LLLA | 3415.16 | 0.799 | 18400 | 1.2 | 1.46 | 1.514 | 5.4 | 20100 (19900, 20300) | 5.9 |
| MIIA | 3487.32 | 0.778 | 15700 | 1.047 | 1.71 | 1.753 | 4.5 | 22100 (21300, 22900) | 6.3 |
| MLIA | 3487.32 | 0.778 | 23100 | 1.2 | 1.884 | 1.967 | 6.6 | 21400 (20900, 21800) | 6.1 |
| MLLA | 3487.32 | 0.778 | 20500 | 1.228 | 1.728 | 1.772 | 5.9 | 20400 (20200, 20600) | 5.8 |
| SIIA | 3310.84 | 0.767 | 15900 | 0.936 | 2.018 | 2.067 | 4.8 | 22000 (21000, 23000) | 6.6 |
| SLIA | 3310.84 | 0.767 | 14200 | 1.177 | 1.491 | 1.526 | 4.3 | 13800 (13500, 14000) | 4.2 |
| SLLA | 3310.84 | 0.767 | 16200 | 1.189 | 1.614 | 1.652 | 4.9 | 16200 (16000, 16300) | 4.9 |

1 Partial specific volume calculated using Sednterp (http://rasmb.org/sednterp/)
2 Mass quoted to 3 significant figures
3 Best-fit frictional ratio
4 Sedimentation coefficient
5 Normalized sedimentation coefficient in water at 20 °C
* Multiple species, reported values refer to major species

**Supplementary Table 3. Summary of biophysical characterization of *de novo* peptides characterized in this study.**

| ***gade*** | **CD Helicity** (%) | **Tm** (°C) | **DPH K$_D$** (µM) | **Fitted Mass/ Monomer mass SV** | **Fitted Mass/ Monomer mass SE** | **Crystal structure** |
|---|---|---|---|---|---|---|
| AIIA†⊥ | 77.3 | >95 | 14.8 ± 3.9 | 6.9 | 5.9 | |
| AILA* | 57.5 | 42 | nd | nd | nd | |
| ALIA** | 60.6 | >95 | 9.4 ± 3.9 | 5.9 | 5.0 | 8qaa, 1.6 Å |
| ALLA† | 87.0 | >95 | 1.2 ± 0.3 | 6.2 | 5.7 | |
| GIIA†⊥ | 71.4 | >95 | 3.9 ± 0.4 | 6.0 | 6.6 | |
| GILA* | 47.3 | 39 | nd | nd | nd | |
| G(IL)$_2$(LI)$_2$A⊥ | 77.7 | >95 | 0.1± 0.05 | 6.3 | 6.6 | |
| GLIA** | 57.4 | >95 | 9.0 ± 1.8 | 6.0 | 6.6 | 8qac, 2.3 Å |
| G(LI)$_2$(IL)$_2$A⊥ | 72.1 | >95 | 0.1± 0.05 | 6.6 | 6.6 | |
| GLLA†⊥ | 52.7 | >95 | 0.2 ± 0.1 | 5.3 | 6.6 | |
| LIIA† | 72.7 | 70 | 1.0 ± 0.3 | 6.6 | 6.4 | |
| LILA* | 67.4 | 31 | nd | nd | nd | |
| LLIA† | 84.5 | 46 | 0.8 ± 0.3 | 5.5 | 5.9 | 8qab, 1.4 Å |
| LLLA† | 92.0 | >95 | 1.8 ± 0.9 | 5.4 | 5.9 | |
| MIIA† | 89.8 | >95 | 0.5 ± 0.2 | 4.5 | 6.3 | |
| MILA* | 48.6 | 35 | nd | nd | nd | |
| MLIA† | 82.3 | >95 | 2.4 ± 1.1 | 6.6 | 6.1 | |
| MLLA† | 84.9 | >95 | 0.6 ± 0.3 | 5.9 | 5.9 | |
| SIIA† | 87.3 | >95 | 2.7 ± 0.7 | 4.8 | 6.6 | |
| SILA* | 58.6 | 53 | nd | nd | nd | |
| SLIA** | 77.5 | >95 | 8.6 ± 2.4 | 4.3 | 4.2 | |
| SLLA† | 77.4 | >95 | 0.1± 0.01 | 4.9 | 4.9 | |

***a*** = Ile with ***d*** = Leu sequences were the least helical and least stable in the peptide screen and were not characterized further beyond CD experiments.

** ***g-a-d-e*** sequence space not modelled in antiparallel αHB protein design as in crystal state were collapsed bundles or smaller oligomer states.

† ***g-a-d-e*** sequence space modelled in sc-apCC-6 design

⊥ ***g-a-d-e*** sequence space modelled in sc-apCC-8 design

##### 2.1.2 *De novo* αHB proteins computational design

**Supplementary Table 4. MASTER hits for each designed protein in study.**

Available upon request to

##### 2.1.3 sc-apCC-6 protein sequences and characterization

**Supplementary Table 5. Sequences of sc-apCC-6 designs**

Available upon request to

**Supplementary Table 6. Evaluated AlphaFold2 metrics for sc-apCC-6 designs in this study.** Grey designs were not characterized experimentally due to poor expression.

| Protein | AlphaFold2 pLDDT | AlphaFold2-to-apCC-Hex-LLIA xtal backbone RMSD (Å) |
|---|---|---|
| sc-apCC-6-ALLA inpainting | 92.42 | 1.943 |
| sc-apCC-6-ALLA lit loops | 86.81 | 4.963 |
| sc-apCC-6-LLIA lit loops | 93.80 | 0.235 |
| sc-apCC-6-LLIA master | 95.01 | 0.241 |
| sc-apCC-6-MIIA inpainting | 94.70 | 1.068 |
| sc-apCC-6-SIIA inpainting | 87.69 | 0.900 |
| sc-apCC-6-SLLA master | 88.58 | 2.288 |

**Supplementary Table 7. Sedimentation velocity AUC fitting statistics for sc-apCC-6 designs in this study.**

| Protein | **Mass** (g/mol) | **v̄**[1] ($cm^3$ $g^{-1}$) | **Fitted Mass**[2] (95% confidence, 3 SF) | **f/f0**[3] | **s**[4] (S) | **s20,w**[5] (S) | **SV weight/ monomer mass** |
|---|---|---|---|---|---|---|---|
| sc-apCC-6-ALLA lit loops | 21121 | 0.76 | 24300 | 1.216 | 2.136 | 2.185 | 1.2 |
| sc-apCC-6-LLIA master | 24206 | 0.78 | 23400 | 1.286 | 1.784 | 1.830 | 1.0 |
| sc-apCC-6-MIIA inpainting | 24163 | 0.765 | 24800 | 1.233 | 2.083 | 2.132 | 1.0 |
| sc-apCC-6-SIIA inpainting | 23104 | 0.756 | 27400 | 1.286 | 1.986 | 2.037 | 1.2 |
| sc-apCC-6-SLLA master | 23566 | 0.756 | 22400 | 1.184 | 2.103 | 2.168 | 1.0 |

1 Partial specific volume calculated using Sednterp (http://rasmb.org/sednterp/)
2 Mass quoted to 3 significant figures
3 Best-fit frictional ratio
4 Sedimentation coefficient
5 Normalized sedimentation coefficient in water at 20 °C
* Multiple species, reported values refer to major species

**Supplementary Table 8. Summary of biophysical characterization for sc-apCC-6 designs in this study.**

| **systematic name** | ***gade*** | **CD Helicity** (%) | **Tm (°C)** | **DPH $K_D$** (μM) | **SV weight/ monomer mass** | **Crystal structure** |
|---|---|---|---|---|---|---|
| sc-apCC-6-ALLA lit loops | ALLA | 71.9 | >95 | 11.7 ± 0.7 | 1.2 | |
| sc-apCC-6-LLIA master | LLIA | 56.3 | >95 | 4.0 ± 0.4 | 1 | 8qad, 2.25 Å |
| sc-apCC-6-MIIA inpainting | MIIA | 66.1 | >95 | 7.6 ± 2 | 1 | |
| sc-apCC-6-SIIA inpainting | SIIA | 58.5 | >95 | 12.3 ± 3 | 1 | |
| sc-apCC-6-SLLA master | SLLA | 74.1 | >95 | 1.5 ± 0.2 | 1 | 8qae, 1.9 Å |

**Supplementary Table 9. SAXS statistics for sc-apCC-6**

| | **sc-apCC-6-ALLA** | **sc-apCC6-LLIA** | **sc-apCC6-MIIA** | **sc-apCC-6-SIIA** | **sc-apCC-6-SLLA** |
|---|---|---|---|---|---|
| Guinier analysis | | | | | |
| $I(0)$ ($cm^{-1}$) | 0.009517 | 0.0237 | 0.03995 | 0.04618 | 0.01595 |
| $R_g$ (Å) | 23.44 | 31.21 | 25.42 | 23.65 | 21.17 |
| P(r) analysis | | | | | |
| $I(0)$ ($cm^{-1}$) | 0.009517 | 0.0237 | 0.03995 | 0.04618 | 0.01595 |
| $R_g$ (Å) | 23.45 | 31.26 | 25.43 | 23.66 | 21.17 |
| $d_{max}$ (Å) | 75.01 | 91.3 | 73.07 | 67.77 | 60.86 |
| $q$ range ($Å^{-1}$) | 0.0159 - 0.2393 | 0.0114 - 0.2346 | 0.0123 - 0.2393 | 0.0125 - 0.2393 | 0.0139 - 0.2393 |
| FoXS | | | | | |
| $\chi^2$, $P$-value | 1.34 | 1.5 | 1.22 | 1.28 | 1.18 |
| Predicted $R_g$ (Å) | 22.73 | 22.26 | 21.6 | 22.01 | 21.87 |
| $c_1$, $c_2$ | 1.02, 2.00 | 1.05, 2.00 | 1.04, 2.00 | 1.03, 2.00 | 0.99, 1.70 |

##### 2.1.4 sc-CC-7 sequences and characterization

**Supplementary Table 10. Sequences of sc-CC-7 designs.**

Available upon request to

**Supplementary Table 11. Evaluated design metrics for expressed sc-CC-7 sequences.** Grey designs were not characterized experimentally due to poor expression.

| | **Exposed hydrophobics** | **Packstat** | **Charge** | **Rosetta energy** | **pI** | **Max pLDDT** |
|---|---|---|---|---|---|---|
| 26 | 4151.91 | 0.641 | +3 | -1346.42 | 8.51 | 97.41 |
| LI | 3672.98 | 0.635 | -9 | -1341.22 | 5.22 | 97.19 |
| IV | 3717.67 | 0.591 | -9 | -1202.69 | 5.22 | 96.7 |
| 80 | 4069.88 | 0.655 | -2 | -1339.59 | 5.90 | 97.31 |
| 99 | 4134.27 | 0.648 | -20 | -1354.32 | 4.80 | 97.07 |

**Supplementary Table 12. Sedimentation velocity AUC fitting statistics for sc-CC-7 proteins in this study.**

| **Protein** | **Mass** (g/mol) | $\bar{v}$[1] (cm$^3$ g$^{-1}$) | **Fitted Mass**[2] (95% confidence, 3 SF) | **f/f0**[3] | **s**[4] (S) | **s20,w**[5] (S) | **SV weight/monomer mass** |
|---|---|---|---|---|---|---|---|
| sc-CC-7-LI | 41252 | 0.768 | 37418 | 1.203 | 2.706 | 2.835 | 0.9 |
| sc-CC-7-IV | 40859 | 0.764 | 36993 | 1.193 | 2.761 | 2.892 | 0.9 |
| sc-CC-7-80 | 41538 | 0.771 | 77200 | 1.194 | 4.357 | 4.567 | 1.9 |

1 Partial specific volume calculated using Sednterp (http://rasmb.org/sednterp/)
2 Mass quoted to 3 significant figures
3 Best-fit frictional ratio
4 Sedimentation coefficient
5 Normalized sedimentation coefficient in water at 20 °C

**Supplementary Table 13. Summary of biophysical characterization for parallel proteins in this study.**

| *gade* | systematic name | CD Helicity (%) | Tm (°C) | DPH $K_D$ (μM) | SV weight/monomer mass | Crystal structure |
|---|---|---|---|---|---|---|
| ALIA | sc-CC-7-LI | 56.2 | >95 | 3.8 ± 0.8 | 1 | 8qai, 2.5 Å |
| AIVA | sc-CC-7-IV | 56.5 | >95 | 2.1 ± 0.3 | 1 | |
| ALIA | sc-CC-7-80 | 74.3 | >95 | 4.0 ± 1.9 | 2 | |

**Supplementary Table 14. SAXS statistics for sc-CC-7**

| | sc-CC-7-LI | sc-CC-7-IV | sc-CC-7-80* |
|---|---|---|---|
| Guinier analysis | | | |
| $I(0)$ ($cm^{-1}$) | 0.073 | 0.0484 | 0.0246 |
| $R_g$ (Å) | 22.7 | 25.3 | 28.9 |
| P(r) analysis | | | |
| $I(0)$ ($cm^{-1}$) | 0.073 | 0.0484 | 0.0246 |
| $R_g$ (Å) | 22.8 | 25.3 | 29 |
| $d_{max}$ (Å) | 73 | 74 | 88 |
| $q$ range (Å$^{-1}$) | 0.01-0.248 | 0.03-0.235 | 0.02-0.250 |
| FoXS | | | |
| $\chi^2$, $P$-value | 1.43 | 1.96 | 6.3 |
| Predicted $R_g$ (Å) | 20.32 | 20.02 | 20.17 |
| $c_1$, $c_2$ | 1.02, 1.63 | 1.03, 2.00 | 1.04, 2.00 |
| * sc-CC-7-80 aggregated at 10 mg/mL, which is observed in the SEC-SAXS collection and AUC. | | | |

##### 2.1.5 sc-ap-CC-8 and sc-CC-5, 6, 8 sequences and characterization

**Supplementary Table 15. Sequences of sc-apCC-8 designs.**

Available upon request to

**Supplementary Table 16. Evaluated AlphaFold2 metrics for sc-apCC-8 designs.**

| Protein | AlphaFold2 pLDDT |
|---|---|
| sc-apCC-8-AIIA master mpnn_1 | 92.02 |
| sc-apCC-8-G(IL)$_2$(LI)$_2$A master mpnn_2 | 92.58 |

**Supplementary Table 17. Sedimentation velocity AUC fitting statistics for sc-apCC-8 designs in this study.**

| Protein | Mass (g/mol) | $\bar{v}$[1] (cm$^3$ g$^{-1}$) | Fitted Mass[2] (95% confidence, 3 SF) | f/f0[3] | s[4] (S) | s20,w[5] (S) | SV weight/ monomer mass |
|---|---|---|---|---|---|---|---|
| sc-apCC-8-AIIA master mpnn_1 | 32092.53 | 0.780 | 34500 | 1.309 | 2.253 | 2.33 | 1.1 |
| sc-apCC-8-G(IL)$_2$(LI)$_2$A master mpnn_2 | 31977.54 | 0.770 | 32100 | 1.261 | 2.343 | 2.419 | 1.0 |

1 Partial specific volume calculated using Sednterp (http://rasmb.org/sednterp/)
2 Mass quoted to 3 significant figures
3 Best-fit frictional ratio
4 Sedimentation coefficient
5 Normalized sedimentation coefficient in water at 20 °C

**Supplementary Table 18. Summary of biophysical characterization for sc-apCC-8 designs in this study.**

| systematic name | *gade* | CD Helicity (%) | Tm (°C) | DPH $K_D$ (μM) | SV weight/monomer mass | Crystal structure |
|---|---|---|---|---|---|---|
| sc-apCC-8-AIIA master mpnn_1 | AIIA | 62.3 | >95 | 0.5 ± 0.1 | 1.1 | 8qaf, 2.0 Å* |
| sc-apCC-8-G(IL)$_2$(LI)$_2$A master mpnn_2 | G(IL)$_2$(LI)$_2$A | 67.4 | >95 | 0.03 ± 0.0.02 | 1 | |

*sc-apCC-8-AIIA crystallized with loops averaged thus only two chains are in the asymmetric unit. $C_4$ symmetry of the structure completes the biological assembly of a single-chain antiparallel αHB protein with 8 helices

**Supplementary Table 19. SAXS statistics for sc-apCC-8**

| | **sc-apCC-8-AIIA** | **sc-apCC-8-G(IL)$_2$(LI)$_2$A*** |
|---|---|---|
| Guinier analysis | | |
| $I(0)$ ($cm^{-1}$) | 0.04097 | 0.008312 |
| $R_g$ (Å) | 28.66 | 32.86 |
| P(r) analysis | | |
| $I(0)$ ($cm^{-1}$) | 0.04098 | 0.008312 |
| $R_g$ (Å) | 28.75 | 32.88 |
| $d_{max}$ (Å) | 88.1 | 93.59 |
| $q$ range (Å$^{-1}$) | 0.0179 - 0.2653 | 0.0097 - 0.200 |
| FoXS | | |
| $\chi^2$, $P$-value | 1.58 | 2.23 |
| Predicted $R_g$ (Å) | 26.78 | 26.92 |
| $c_1$, $c_2$ | 1.05, 2.00 | 1.05, 2.00 |

* sc-apCC-8-G(IL)$_2$(LI)$_2$ aggregated at 10 mg/mL, which is observed in the SEC-SAXS collection and in attempting to set up crystallization trials with this construct.

**Supplementary Table 20. Sequences of sc-CC-5 designs.**

Available upon request to

**Supplementary Table 21. Evaluated design metrics for expressed sc-CC-5 sequences.**

| | Exposed hydrophobics | Packstat | Charge | Rosetta energy | pI | Max pLDDT |
|---|---|---|---|---|---|---|
| 17 | 2727.41 | 0.541 | +3 | -832.63 | 8.61 | 95.69 |
| 24 | 2506.40 | 0.548 | -6 | -808.49 | 5.46 | 95.97 |
| 77 | 2372.89 | 0.530 | -4 | -812.06 | 5.47 | 95.94 |

**Supplementary Table 22. Sequences of sc-CC-6 designs.**

Available upon request to

**Supplementary Table 23. Evaluated design metrics for expressed sc-CC-6 sequences.** Grey designs were not characterized experimentally due to poor expression.

| | Exposed hydrophobics | Packstat | Charge | Rosetta energy | pI | Max pLDDT |
|---|---|---|---|---|---|---|
| 67 | 3160.15 | 0.605 | -8 | -1059.53 | 5.24 | 97.61 |
| 77 | 3155.46 | 0.626 | +6 | -1091.85 | 8.96 | 97.56 |
| 90 | 2932.13 | 0.655 | +7 | -1059.02 | 8.97 | 97.52 |
| 95 | 3086.07 | 0.616 | -6 | -1069.84 | 5.37 | 97.46 |

**Supplementary Table 24. Sequences of sc-CC-8 designs.**

Available upon request to

**Supplementary Table 25. Design metrics for expressed sc-CC-8 sequences.** Grey designs were not characterized experimentally due to poor expression.

| | Exposed hydrophobics | Packstat | Charge | Rosetta energy | pI | Max pLDDT |
|---|---|---|---|---|---|---|
| 13 | 4901.98 | 0.629 | -2 | -1385.76 | 5.89 | 96.77 |
| 22 | 5053.44 | 0.635 | +5 | -1394.39 | 8.73 | 96.77 |
| 58 | 4498.68 | 0.609 | +5 | -1383.1 | 8.75 | 96.67 |
| 84 | 4698.42 | 0.652 | +5 | -1381.11 | 8.73 | 96.32 |

**Supplementary Table 26. Sedimentation velocity AUC fitting statistics for sc-CC-5, 6, and 8 proteins in this study.**

| **Protein** | **Mass** (g/mol) | $\bar{v}$[1] ($cm^3$ $g^{-1}$) | **Fitted Mass**[2] (95% confidence, 3 SF) | **f/f0**[3] | **s**[4] (S) | **s20,w**[5] (S) | **SV weight/monomer mass** |
|---|---|---|---|---|---|---|---|
| sc-CC-5-17 | 32369 | 0.758 | 27508 | 1.196 | 2.325 | 2.434 | 0.85 |
| sc-CC-5-24 | 32298 | 0.753 | 27750 | 1.203 | 2.380 | 2.490 | 0.9 |
| sc-CC-5-77 | 32079 | 0.751 | 28125 | 1.205 | 2.408 | 2.519 | 0.9 |
| sc-CC-6-67 | 35478 | 0.767 | 31241 | 1.193 | 2.432 | 2.548 | 0.9 |
| sc-CC-6-90 | 35716 | 0.767 | 32714 | 1.208 | 2.477 | 2.595 | 0.9 |
| sc-CC-6-95 | 35389 | 0.764 | 35600 | 1.225 | 2.601 | 2.725 | 1.0 |
| sc-CC-8-22 | 46252 | 0.767 | 41336 | 1.199 | 2.916 | 3.056 | 0.9 |
| sc-CC-8-58 | 46006 | 0.766 | 45100 | 1.191 | 3.085 | 3.243 | 1.0 |
| sc-CC-8-84 | 45940 | 0.768 | 42757 | 1.209 | 2.943 | 3.084 | 0.9 |

1 Partial specific volume calculated using Sednterp (http://rasmb.org/sednterp/)
2 Mass quoted to 3 significant figures
3 Best-fit frictional ratio
4 Sedimentation coefficient
5 Normalized sedimentation coefficient in water at 20 °C

**Supplementary Table 27. Summary of biophysical characterization for sc-CC-5, 6, and 8 proteins in this study.**

| *gade* | systematic name | CD Helicity (%) | Tm (°C) | DPH $K_D$ (μM) | SV weight/monomer mass | Crystal structure |
|---|---|---|---|---|---|---|
| TLIA | sc-CC-5-17 | 56.2 | >95 | - | 1 | |
| TLIA | sc-CC-5-24 | 67.5 | >95 | - | 1 | |
| TLIA | sc-CC-5-77 | 62.0 | >95 | - | 1 | |
| ALIA | sc-CC-6-67 | 58.5 | >95 | 0.27 ± 0.11 | 1 | |
| ALIA | sc-CC-6-90 | 63.9 | >95 | 1.0 ± 0.1 | 1 | |
| ALIA | sc-CC-6-95 | 57.6 | >95 | 0.23 ± 0.05 | 1 | 8qag, 2.8 Å |
| AIIA | sc-CC-8-22 | 63.0 | >95 | 1.3 ± 0.2 | 1 | |
| AIIA | sc-CC-8-58 | 71.2 | >95 | 2.3 ± 0.6 | 1 | 8qah, 2.35 Å |
| AIIA | sc-CC-8-84 | 65.3 | >95 | 0.99 ± 0.42 | 1 | |

**Supplementary Table 28. SAXS statistics for sc-CC-5, 6, and 8 proteins in this study.**

| | **sc-CC-5-17** | **sc-CC-5-24** | **sc-CC-5-77*** |
|---|---|---|---|
| Guinier analysis | | | |
| $I(0)$ ($cm^{-1}$) | 0.0126 | 0.0226 | 0.0239 |
| $R_g$ (Å) | 28.3 | 24.9 | 33.1 |
| P(r) analysis | | | |
| $I(0)$ ($cm^{-1}$) | 0.0125 | 0.0226 | 0.0239 |
| $R_g$ (Å) | 28.3 | 24.9 | 33.2 |
| $d_{max}$ (Å) | 85 | 74 | 106 |
| $q$ range ($Å^{-1}$) | 0.02-0.272 | 0.01-0.254 | 0.01-0.226 |
| FoXS | | | |
| $\chi^2$, *P*-value | 3.02 | 1.52 | 14.69 |
| Predicted $R_g$ (Å) | 20.81 | 20.41 | 20.37 |
| $c_1$, $c_2$ | 1.05, 2.00 | 1.04, 2.00 | 1.05, 2.0 |

* sc-CC-5-77 aggregated at 10 mg/mL, which is observed in the SEC-SAXS collection.

**Supplementary Table 28 cont.**

| | sc-CC-6-67 | sc-CC-6-90 | sc-CC-6-95 |
|---|---|---|---|
| Guinier analysis | | | |
| $I(0)$ ($cm^{-1}$) | 0.0255 | 0.0232 | 0.0221 |
| $R_g$ (Å) | 25.2 | 21.1 | 26.3 |
| P(r) analysis | | | |
| $I(0)$ ($cm^{-1}$) | 0.0255 | 0.0234 | 0.0221 |
| $R_g$ (Å) | 25.2 | 21 | 26.3 |
| $d_{max}$ (Å) | 77 | 51 | 80 |
| $q$ range (Å$^{-1}$) | 0.0167-0.2159 | 0.012-0.213 | 0.02-0.233 |
| FoXS | | | |
| $\chi^2$, $P$-value | 3.44 | 1.14 | 2.15 |
| Predicted $R_g$ (Å) | 19.25 | 19.57 | 19.94 |
| $c_1$, $c_2$ | 1.04, 2.00 | 1.03, 2.00 | 1.04, 2.00 |

**Supplementary Table 28 cont.**

| | **sc-CC-8-22** | **sc-CC-8-58** | **sc-CC-8-84** |
|---|---|---|---|
| Guinier analysis | | | |
| $I(0)$ ($cm^{-1}$) | 0.0592 | 0.0401 | 0.0454 |
| $R_g$ (Å) | 25.2 | 22.6 | 30.7 |
| P(r) analysis | | | |
| $I(0)$ ($cm^{-1}$) | 0.0592 | 0.0401 | 0.0454 |
| $R_g$ (Å) | 25.2 | 22.6 | 30.8 |
| $d_{max}$ (Å) | 77 | 65 | 60.5 |
| $q$ range ($Å^{-1}$) | 0.030-0.253 | 0.013-0.239 | 0.030-0.239 |
| FoXS | | | |
| $\chi^2$, *P*-value | 1.41 | 1.67 | 2.38 |
| Predicted $R_g$ (Å) | 21.44 | 22.4 | 21.32 |
| $c_1$, $c_2$ | 1.03, 1.80 | 1.03, -0.30 | 1.04, 2.00 |

#### 2.1.6 Structural characterization statistics and measurements

**Supplementary Table 29. Crystallization conditions used to obtain the structures discussed in this study.**

| Systematic name | Crystal structure | Crystallization condition | Seed condition |
|---|---|---|---|
| CC-Type2-EEKK-($A_gL_aI_d$)$_4$ | apCC-Hex-ALIA collapsed bundle | Morpheus C12: 12.5% w/v PEG 1000, 12.5% w/v PEG 3350, 12.5% v/v MPD, 0.03 M of each NPS, 0.1 M bicine/Trizma base pH 8.5 | None |
| CC-Type2-EEKK-($G_gL_aI_d$)$_4$ | apCC-Oct-GLIA collapsed bundle | Morpheus D9: 10% w/v PEG 20000, 20% v/v PEG MME 550, 0.02 M of each alcohol, 0.1 M bicine/Trizma base pH 8.5 | None |
| CC-Type2-EEKK-($L_gL_aI_d$)$_4$ | apCC-Hex | JCSG-plus G9: 0.1 M Potassium thiocyanate, 30 % w/v PEG 2000 MME | None |
| sc-apCC-6-LLIA master | sc-apCC-6-LLIA | JCSG-plus A10: 0.2 M potassium formate, 20% w/v PEG 3350 | PACT Premier B11: 0.2 M CaCl2, 0.1 M MES pH 6.0, 20% w/v PEG 6000 |
| sc-apCC-6-SLLA master | sc-apCC-6-SLLA | PACT Premier B11: 0.2 M CaCl2, 0.1 M MES pH 6.0, 20% w/v PEG 6000 | PACT Premier B11: 0.2 M CaCl2, 0.1 M MES pH 6.0, 20% w/v PEG 6000 |
| sc-apCC-8-AIIA master mpnn 1 | sc-apCC-8-AIIA | PACT Premier H6: 0.2 M sodium formate, 0.1 M Bis Tris propane, pH 8.5, 20 % w/v PEG 3350 | None |
| sc-CC-6-95 | sc-CC-6-95 | 75% JCSG A6: 0.2 M lithium sulfate, 0.1 M phosphate-citrate pH 4.2, 20% PEG 1000.<br>25% Structure Screen 1 & 2 A11: 0.2 M<br>calcium chloride dihydrate, 0.1 M sodium acetate, pH 4.6, 20% w/v 2-propanol | None |
| sc-CC-7-LI | sc-CC-7-LI | JCSG-plus C2: 1.0 M Lithium chloride, 0.1 M citrate, pH 4.0, 20 % w/v PEG 6000 | None |
| sc-CC-8-58 | sc-CC-8-58 | Pact Premier D11: 0.2 M calcium chloride, 0.1 M TRICS, pH 8, 20 % w/v PEG 6000 | Pact Premier C2: 0.1 M PCTP, pH 5.0, 25 % w/v PEG 1500 |

**Supplementary Table 30. Merging and refinement statistics for all X-ray crystal structure**s.

| ***de novo* peptides** | **g-a-d-e = ALIA** | **g-a-d-e = GLIA** | **apCCHex-LLIA** |
|---|---|---|---|
| **PDB ID** | 8QAA | 8QAC | 8QAB |
| **Data Collection** | | | |
| Source | Diamond I04 | Diamond I04 | Diamond I24 |
| Detector | EIGER2 XE 16M | EIGER2 XE 16M | PILATUS3 6M |
| Wavelength (Å) | 0.98 | 0.98 | 0.85 |
| Resolution range | 44.75 - 1.6 (1.657 - 1.6) | 35.68 - 2.3 (2.382 - 2.3) | 34.88 - 1.4 (1.45 - 1.4) |
| Space group | P 1 21 1 | P 1 21 1 | P 1 21 1 |
| Unit cell: *a, b, c (Å)* | 29.2744 89.4965 32.6667 | 71.2714 45.8732 72.8802 | 46.0784 30.9144 51.9531 |
| *α, β, γ (°)* | 90 115.212 90 | 90 103.943 90 | 90 88.6417 90 |
| Total reflections | 138060 (12752) | 144912 (14403) | 175748 (14321) |
| Unique reflections | 19749 (1934) | 20647 (2038) | 29174 (2850) |
| Multiplicity | 7.0 (6.6) | 7.0 (7.1) | 6.0 (5.0) |
| Completeness (%) | 98.26 (96.93) | 99.51 (99.46) | 99.83 (99.20) |
| Mean I/sigma(I) | 8.63 (1.11) | 14.52 (2.45) | 22.17 (0.67) |
| Wilson B-factor | 17.31 | 44.63 | 20.99 |
| R-merge | 0.1184 (0.8176) | 0.08693 (0.6312) | 0.03595 (0.8515) |
| R-meas | 0.1285 (0.8881) | 0.09436 (0.6822) | 0.03944 (0.9542) |
| R-pim | 0.04923 (0.3437) | 0.03623 (0.2564) | 0.01597 (0.421) |
| CC1/2 | 0.997 (0.783) | 0.99 (0.903) | 1 (0.625) |
| CC* | 0.999 (0.937) | 0.997 (0.974) | 1 (0.877) |
| **Refinement** | | | |
| Reflections used in refinement | 19709 (1929) | 20552 (2027) | 29127 (2850) |
| Reflections used for R- free | 1005 (87) | 1033 (104) | 1415 (158) |
| R-work | 0.1808 (0.2532) | 0.2285 (0.2433) | 0.1740 (0.2925) |
| R-free | 0.2247 (0.3219) | 0.2782 (0.3019) | 0.1893 (0.3332) |
| CC(work) | 0.973 (0.883) | 0.934 (0.917) | 0.718 (0.086) |
| CC(free) | 0.962 (0.828) | 0.851 (0.782) | 0.929 (0.053) |
| Number of non-hydrogen atoms | 1437 | 2969 | 1468 |
| macromolecules | 1248 | 2949 | 1284 |
| ligands | 30 | 0 | 93 |
| solvent | 171 | 20 | 143 |
| Protein residues | 180 | 456 | 179 |
| RMS(bonds) | 0.009 | 0.001 | 0.011 |
| RMS(angles) | 1.35 | 0.28 | 1.13 |
| Ramachandran favored (%) | 100 | 99.53 | 100 |
| Ramachandran allowed (%) | 0 | 0.47 | 0 |
| Ramachandran outliers (%) | 0 | 0 | 0 |
| Rotamer outliers (%) | 0 | 0 | 0 |
| Clashscore | 2.36 | 0.52 | 1.11 |
| Average B-factor | 21.82 | 54.94 | 28.45 |
| macromolecules | 20.14 | 54.96 | 26.58 |
| ligands | 30.85 | 52.57 | 44.91 |
| solvent | 33.09 | | 40.51 |
| Number of TLS groups | 6 | 16 | 6 |

**Supplementary Table 30 cont.**

| **antiparallel α-HB proteins** | **sc-apCC-6-LLIA** | **sc-apCC-6-SLLA** | **sc-apCC-8-AIIA** |
|---|---|---|---|
| **PDB ID** | 8QAD | 8QAE | 8QAF |
| **Data Collection** | | | |
| Source | Diamond I24 | Diamond I04 | Diamond I24 |
| Detector | PILATUS3 6M | EIGER2 XE 16M | PILATUS3 6M |
| Wavelength (Å) | 0.999 | 0.92 | 0.999 |
| Resolution range | 47.98 - 2.25 (2.33 - 2.25) | 21.11 - 1.9 (1.968 - 1.9) | 37.66 - 2.0 (2.072 - 2.0) |
| Space group | P 21 21 21 | C 1 2 1 | P 4 21 2 |
| Unit cell: *a, b, c (Å)* | 34.549 60.875 77.949 | 126.823 42.1757 71.5217 | 50.002 50.002 57.249 |
| *α, β, γ (°)* | 90 90 90 | 90 123.509 90 | 90 90 90 |
| Total reflections | 16509 (1600) | 169771 (14896) | 10580 (1008) |
| Unique reflections | 8255 (800) | 25168 (2501) | 5290 (504) |
| Multiplicity | 2.0 (2.0) | 6.7 (6.0) | 2.0 (2.0) |
| Completeness (%) | 99.64 (100.00) | 99.84 (99.92) | 99.62 (100.00) |
| Mean I/sigma(I) | 9.72 (3.00) | 14.57 (1.66) | 15.06 (5.24) |
| Wilson B-factor | 46.33 | 35.53 | 35.06 |
| R-merge | 0.02253 (0.2053) | 0.08096 (0.4644) | 0.01179 (0.09224) |
| R-meas | 0.03186 (0.2903) | 0.08813 (0.5099) | 0.01668 (0.1305) |
| R-pim | 0.02253 (0.2053) | 0.03443 (0.2069) | 0.01179 (0.09224) |
| CC1/2 | 0.999 (0.927) | 0.985 (0.903) | 1 (0.976) |
| CC* | 1 (0.981) | 0.996 (0.974) | 1 (0.994) |
| **Refinement** | | | |
| Reflections used in refinement | 8225 (800) | 25150 (2499) | 5272 (504) |
| Reflections used for R-free | 373 (40) | 1347 (123) | 248 (26) |
| R-work | 0.2488 (0.2529) | 0.2077 (0.2651) | 0.2496 (0.2656) |
| R-free | 0.2781 (0.2464) | 0.2308 (0.2609) | 0.2520 (0.3244) |
| CC(work) | 0.953 (0.924) | 0.955 (0.890) | 0.934 (0.918) |
| CC(free) | 0.934 (0.863) | 0.934 (0.874) | 0.997 (0.959) |
| Number of non-hydrogen atoms | 972 | 2314 | 386 |
| macromolecules | 952 | 2118 | 372 |
| ligands | 38 | 110 | 0 |
| solvent | 3 | 149 | 14 |
| Protein residues | 145 | 310 | 59 |
| RMS(bonds) | 0.001 | 0.003 | 0.006 |
| RMS(angles) | 0.26 | 0.5 | 0.57 |
| Ramachandran favored (%) | 100 | 98.66 | 100 |
| Ramachandran allowed (%) | 0 | 1.34 | 0 |
| Ramachandran outliers (%) | 0 | 0 | 0 |
| Rotamer outliers (%) | 0 | 0 | 0 |
| Clashscore | 0 | 1.66 | 1.36 |
| Average B-factor | 58.91 | 42.93 | 41.25 |
| macromolecules | 58.75 | 42.59 | 41.02 |
| ligands | 65.28 | 47.66 | |
| solvent | 75.6 | 46.22 | 47.25 |
| Number of TLS groups | 1 | 2 | 1 |

**Supplementary Table 30 cont.**

| **parallel α-HB proteins** | **sc-CC-6-95** | **sc-CC-7-LI** | **sc-CC-8-58** |
|---|---|---|---|
| **PDB ID** | 8QAG | 8QAI | 8QAH |
| **Data Collection** | | | |
| Source | Diamond I24 | Diamond I04 | Diamond I24 |
| Detector | PILATUS3 6M | EIGER2 XE 16M | PILATUS3 6M |
| Wavelength (Å) | 0.999 | 0.92 | 0.999 |
| Resolution range | 57.98 - 2.8 (2.9 - 2.8) | 30.63 - 2.5 (2.589 - 2.5) | 49.09 - 2.35 (2.434 - 2.35) |
| Space group | P 43 21 2 | P1 | P 1 21 1 |
| Unit cell: *a, b, c (Å)* | 88.9498 88.9498 149.518 | 126.823 42.1757 71.5217 | 46.054 78.587 98.174 |
| *α, β, γ (°)* | 90 90 90 | 99.4037 100.941 105.332 | 90 90.145 90 |
| Total reflections | 768941 (75362) | 58865 (5729) | 57509 (5634) |
| Unique reflections | 15396 (1488) | 23776 (2330) | 29247 (2871) |
| Multiplicity | 49.9 (50.6) | 2.5 (2.5) | 2.0 (2.0) |
| Completeness (%) | 99.83 (99.07) | 98.44 (97.24) | 99.56 (97.85) |
| Mean I/sigma(I) | 15.29 (1.87) | 9.11 (1.79) | 5.98 (1.81) |
| Wilson B-factor | 69.96 | 39.62 | 14.18 |
| R-merge | 0.1387 (0.6924) | 0.08575 (0.2149) | 0.09588 (0.4639) |
| R-meas | 0.1402 (0.6993) | 0.1068 (0.2704) | 0.1356 (0.6561) |
| R-pim | 0.02015 (0.0976) | 0.06252 (0.1616) | 0.09588 (0.4639) |
| CC1/2 | 0.999 (0.972) | 0.991 (0.923) | 0.988 (0.681) |
| CC* | 1 (0.993) | 0.998 (0.98) | 0.997 (0.9) |
| **Refinement** | | | |
| Reflections used in refinement | 15385 (1487) | 23743 (2326) | 29191 (2869) |
| Reflections used for R-free | 768 (78) | 1572 (154) | 1393 (121) |
| R-work | 0.2284 (0.3309) | 0.1986 (0.2174) | 0.1934 (0.2503) |
| R-free | 0.2725 (0.3615) | 0.2362 (0.2985) | 0.2341 (0.3127) |
| CC(work) | 0.949 (0.870) | 0.961 (0.908) | 0.966 (0.838) |
| CC(free) | 0.927 (0.792) | 0.963 (0.842) | 0.955 (0.725) |
| Number of non-hydrogen atoms | 3864 | 4866 | 5534 |
| macromolecules | 3833 | 4832 | 5392 |
| ligands | 27 | 7 | 0 |
| solvent | 4 | 27 | 142 |
| Protein residues | 594 | 732 | 851 |
| RMS(bonds) | 0.025 | 0.002 | 0.003 |
| RMS(angles) | 1.08 | 0.24 | 0.44 |
| Ramachandran favored (%) | 98.96 | 99.58 | 99.76 |
| Ramachandran allowed (%) | 1.04 | 0.42 | 0.24 |
| Ramachandran outliers (%) | 0 | 0 | 0 |
| Rotamer outliers (%) | 1.63 | 0.31 | 0.96 |
| Clashscore | 8.48 | 1.36 | 2.81 |
| Average B-factor | 74.95 | 44.93 | 24.25 |
| macromolecules | 75.1 | 44.86 | 24 |
| ligands | 56.59 | 60.15 | - |
| solvent | 54.27 | 54.39 | 33.88 |
| Number of TLS groups | 2 | 2 | 16 |

**Supplementary Table 31. Backbone Cα alignments for all protein structures, AlphaFold2 models and starting backbone templates in this work.** Backbone Cα alignments are calculated for all protein structures, AlphaFold2[5,6] models and starting backbone templates in this work. For the antiparallel αHB proteins, the full $RMSD_{bb}$ is equivalent to the inner αHB $RMSD_{bb}$.

| | **AlphaFold2** $RMSD_{bb}$ (Å) | **Original backbone** | | |
|---|---|---|---|---|
| | | Inner αHB $RMSD_{bb}$ (Å) | Outer 3HB $RMSD_{bb}$ (Å) | Full $RMSD_{bb}$ (Å) |
| sc-apCC-6-LLIA | 0.269 | - | - | 1.177 |
| sc-apCC-6-SLLA | 14.144 | - | - | 7.913 |
| sc-apCC-8 | 0.501 | - | - | * |
| sc-CC-6-95 | 0.300 | 0.584 | 0.538 | 0.781 |
| sc-CC-7-LI | 0.433 | 0.930 | 0.574 | 1.032 |
| sc-CC-8-58 | 0.530 | 1.057 | 0.656 | 0.979 |
| *CC-Type2-$A_gI_aI_dA_e$ is a parallel octamer in a crystal structure[3]. As discussed above, the starting seed for the sc-apCC-8 designs was the AlphaFold2[5] model for each respective peptide assembly, thus the AlphaFold2[5,6] $RMSD_{bb}$ represents the alignment to both the peptide and protein models. | | | | |

**Supplementary Table 32. Top 10 Foldseek[37] results for sc-apCC-6-SLLA against AlphaFold2-Swissprot database[5,38].**

| **Accession code** | **Sequence identity, %** | **Probability** | **E-score** | **Length** | **RMSD, Å** | **TM-score** |
|---|---|---|---|---|---|---|
| P56485 | 8.3 | 0.941 | 0.07883 | 202 | 13.69 | 0.232 |
| Q8K209 | 13.8 | 0.933 | 0.07883 | 207 | 19.52 | 0.228 |
| Q9SI74 | 13.3 | 0.887 | 0.7489 | 175 | 15.57 | 0.262 |
| Q5W6M3 | 14.8 | 0.872 | 0.2123 | 174 | 20.42 | 0.257 |
| O23447 | 10.6 | 0.795 | 0.7489 | 190 | 18.22 | 0.226 |
| Q57674 | 9.1 | 0.772 | 0.7834 | 205 | 13.28 | 0.267 |
| Q99N03 | 8.9 | 0.747 | 2.891 | 133 | 6.71 | 0.407 |
| Q646D1 | 8.1 | 0.663 | 0.3483 | 202 | 14.79 | 0.207 |
| P0CB17 | 11.7 | 0.632 | 1.123 | 140 | 11.27 | 0.267 |
| Q7TN49 | 6.4 | 0.632 | 0.3987 | 202 | 19.49 | 0.250 |

**Supplementary Table 33. Top10 Foldseek[37] results for sc-apCC-6-SLLA against PDB database[39,40] filtered with 90% sequence identity.**

| Accession code | Sequence identity, % | Probability | E-score | Length | RMSD, Å | TM-score |
|---|---|---|---|---|---|---|
| 4ffx_C | 13.1 | 0.99 | 0.02861 | 199 | 13.01 | 0.256 |
| 4xt3_A | 7 | 0.872 | 0.1733 | 203 | 14.36 | 0.278 |
| 3d19_B | 11.9 | 0.747 | 0.3726 | 204 | 14.55 | 0.210 |
| 7sk8_A | 7.6 | 0.72 | 0.1983 | 206 | 16.67 | 0.230 |
| 7yu4_A | 7.6 | 0.632 | 0.5845 | 200 | 18.04 | 0.268 |
| 5wb2_A | 7.4 | 0.632 | 0.3562 | 201 | 14.3 | 0.213 |
| 1t8b_B | 12 | 0.601 | 0.227 | 209 | 21.01 | 0.178 |
| 4q25_B | 9.8 | 0.601 | 0.2718 | 205 | 19.95 | 0.184 |
| 1sum_B | 12.1 | 0.569 | 0.2975 | 198 | 19.28 | 0.180 |
| 7d77_R | 11.6 | 0.505 | 0.4461 | 202 | 12.98 | 0.236 |

**Supplementary Table 34. Top 10 Foldseek[37] results for sc-apCC-6-LLIA against AlphaFold2-Swissprot database[5,38].**

| Accession code | Sequence identity, % | Probability | E-score | Length | RMSD, Å | TM-score |
|---|---|---|---|---|---|---|
| Q51417 | 13.1 | 0.998 | 0.1764 | 143 | 4 | 0.569 |
| P56583 | 11.5 | 0.663 | 1.915 | 140 | 10.89 | 0.447 |
| O28133 | 15.6 | 0.382 | 0.5262 | 142 | 13.98 | 0.277 |
| P9WI95 | 12.7 | 0.382 | 1.494 | 142 | 9.53 | 0.338 |
| P9WI94 | 12.7 | 0.214 | 3.308 | 142 | 9.53 | 0.337 |
| P65721 | 13.8 | 0.196 | 2.995 | 142 | 9.79 | 0.327 |
| Q83VR1 | 12.1 | 0.15 | 7.697 | 65 | 5.07 | 0.498 |
| M2WJF5 | 8.1 | 0.138 | 2.115 | 143 | 13.65 | 0.314 |
| P09384 | 16.7 | 0.116 | 5.713 | 141 | 13.74 | 0.375 |
| A1VD51 | 7.7 | 0.098 | 6.004 | 91 | 6.75 | 0.374 |

**Supplementary Table 35. Top10 Foldseek[37] results for sc-apCC-6-LLIA against PDB database[39,40] filtered with 90% sequence identity.** * denotes *de novo* proteins.

| Accession code | Sequence identity, % | Probability | E-score | Length | RMSD, Å | TM-score |
|---|---|---|---|---|---|---|
| 5l0j_B | 14.6 | 0.817 | 0.6451 | 92 | 5.06 | 0.504 |
| 1sum_B | 17.7 | 0.473 | 0.1773 | 140 | 12.97 | 0.264 |
| 3vf0_A | 14.9 | 0.301 | 4.262 | 92 | 4.9 | 0.529 |
| 2ccy_B | 18.5 | 0.277 | 1.743 | 115 | 11.85 | 0.443 |
| 2b0h_A | 10.7 | 0.233 | 1.925 | 141 | 10.25 | 0.444 |
| * 4uos_A | 22 | 0.106 | 2.593 | 143 | 13.81 | 0.289 |
| 3zc1_F | 18.4 | 0.106 | 3.672 | 141 | 13.87 | 0.327 |
| 1jsw_B | 10.4 | 0.106 | 8.131 | 143 | 8.54 | 0.314 |
| 5nqn_A | 8.4 | 0.082 | 4.262 | 141 | 14.54 | 0.356 |
| * 6xr2_D | 25.4 | 0.082 | 8.131 | 143 | 9.03 | 0.351 |

**Supplementary Table 36. Top 10 Foldseek[37] results for sc-apCC-8 against AlphaFold2-Swissprot database[5,38].**

| Accession code | Sequence identity, % | Probability | E-score | Length | RMSD, Å | TM-score |
|---|---|---|---|---|---|---|
| Q15722 | 8.9 | 0.933 | 0.1293 | 239 | 14.69 | 0.342 |
| Q9FK05 | 7.6 | 0.855 | 2.137 | 126 | 5.71 | 0.527 |
| Q9R0Q2 | 8.8 | 0.692 | 0.5619 | 239 | 14.29 | 0.345 |
| P30875 | 8.6 | 0.692 | 0.2881 | 244 | 17.91 | 0.278 |
| P41146 | 9 | 0.692 | 0.3935 | 242 | 18.23 | 0.283 |
| P30680 | 8.6 | 0.663 | 0.3443 | 244 | 17.91 | 0.278 |
| P30874 | 8.4 | 0.663 | 0.4702 | 244 | 17.41 | 0.300 |
| Q9UKP6 | 6.9 | 0.663 | 0.6421 | 244 | 12.77 | 0.331 |
| P35377 | 9.7 | 0.601 | 0.4916 | 244 | 18.11 | 0.292 |
| P34993 | 8.6 | 0.601 | 0.36 | 241 | 7.05 | 0.312 |

**Supplementary Table 37. Top10 Foldseek[37] results for sc-apCC-8 against PDB database[39,40] filtered with 90% sequence identity.** * denotes *de novo* proteins.

| Accession code | Sequence identity, % | Probability | E-score | Length | RMSD, Å | TM-score |
|---|---|---|---|---|---|---|
| 2cj7_A | 13.8 | 0.632 | 1.822 | 127 | 8.00 | 0.456 |
| 3d19_B | 11.8 | 0.569 | 1.22 | 234 | 8.00 | 0.382 |
| 4bv0_A | 8.2 | 0.505 | 0.4582 | 239 | 15.35 | 0.311 |
| 7xmr_R | 6.7 | 0.382 | 1.276 | 241 | 15.34 | 0.326 |
| 5dhh_A | 8.7 | 0.353 | 1.991 | 242 | 13.24 | 0.350 |
| 4z35_A | 11.3 | 0.326 | 0.7477 | 237 | 22.24 | 0.272 |
| 7vgv_B | 9.7 | 0.277 | 2.488 | 243 | 14.00 | 0.365 |
| * 5cwo_B | 16.4 | 0.254 | 1.276 | 235 | 12.10 | 0.286 |
| 6ibb_C | 6.2 | 0.254 | 2.276 | 241 | 14.00 | 0.341 |
| 7piu_R | 9.1 | 0.233 | 2.177 | 238 | 15.05 | 0.334 |

**Supplementary Table 38. Top 10 Foldseek[37] results for sc-CC-6-95 against AlphaFold2-Swissprot database[5,38].**

| Accession code | Sequence identity, % | Probability | E-score | Length | RMSD, Å | TM-score |
|---|---|---|---|---|---|---|
| P37630 | 12 | 1 | 0.01674 | 305 | 9.27 | 0.499 |
| O59815 | 9.9 | 0.999 | 0.1156 | 310 | 8.57 | 0.531 |
| P50537 | 10.2 | 0.956 | 0.4927 | 305 | 10.14 | 0.501 |
| A9JR78 | 13.8 | 0.933 | 0.1262 | 265 | 13.23 | 0.418 |
| D4A6D7 | 17.7 | 0.923 | 0.07787 | 267 | 14.94 | 0.355 |
| Q8CC21 | 17.4 | 0.912 | 0.09282 | 265 | 14.51 | 0.357 |
| P0CT95 | 8 | 0.887 | 0.7989 | 296 | 9.15 | 0.487 |
| P0CT97 | 8.2 | 0.872 | 0.7989 | 291 | 9.32 | 0.467 |
| Q9VSA4 | 14.1 | 0.872 | 0.1505 | 252 | 12.7 | 0.399 |
| P0CT94 | 7.7 | 0.855 | 0.8347 | 293 | 9.25 | 0.466 |

**Supplementary Table 39. Top 10 Foldseek[37] results for sc-CC-6-95 against PDB database[39,40] filtered with 90% sequence identity.** * denotes *de novo* proteins.

| Accession code | Sequence identity, % | Probability | E-score | Length | RMSD, Å | TM-score |
|---|---|---|---|---|---|---|
| * 5cwq_A | 43.3 | 1 | 3.07E-08 | 218 | 1.21 | 0.956 |
| * 6xr1_A | 32.8 | 1 | 1.97E-10 | 285 | 6.32 | 0.66 |
| * 6xr2_C | 33.3 | 1 | 7.65E-05 | 164 | 3.2 | 0.779 |
| * 6e9t_B | 26.7 | 1 | 0.0001686 | 230 | 6.67 | 0.612 |
| * 7rdr_A | 27.6 | 1 | 1.06E-05 | 303 | 13.05 | 0.441 |
| * 6e9y_A | 27.1 | 1 | 0.0006026 | 253 | 10.06 | 0.57 |
| * 5cwn_A | 27.6 | 1 | 0.02753 | 223 | 8.03 | 0.493 |
| * 6xns_B | 18.5 | 0.997 | 0.008046 | 307 | 20.33 | 0.422 |
| 4a1s_B | 15.7 | 0.988 | 0.007701 | 285 | 15.14 | 0.353 |
| 5a6c_A | 15.8 | 0.961 | 0.02753 | 288 | 16.12 | 0.362 |

**Supplementary Table 40. Top 10 Foldseek[37] results for sc-CC-7-LI against AlphaFold2-Swissprot database[5,38].**

| Accession code | Sequence identity, % | Probability | E-score | Length | RMSD, Å | TM-score |
|---|---|---|---|---|---|---|
| P37630 | 12.1 | 0.993 | 0.03357 | 368 | 15.33 | 0.430 |
| D4A6D7 | 14.4 | 0.988 | 0.02954 | 314 | 12.6 | 0.401 |
| Q8CC21 | 14.4 | 0.984 | 0.0185 | 312 | 12.26 | 0.370 |
| P44741 | 12.6 | 0.837 | 0.4132 | 245 | 7.11 | 0.433 |
| A4QP73 | 14.4 | 0.837 | 0.1203 | 312 | 16.18 | 0.367 |
| Q32NU8 | 15.7 | 0.663 | 0.248 | 316 | 21.28 | 0.351 |
| Q54SU1 | 11.5 | 0.663 | 0.3486 | 322 | 20.55 | 0.364 |
| Q9LD83 | 11 | 0.569 | 1.419 | 305 | 11.12 | 0.485 |
| Q2U0E0 | 15.9 | 0.505 | 0.7823 | 193 | 14.05 | 0.411 |
| Q9FLV9 | 13.3 | 0.442 | 1.249 | 251 | 7.91 | 0.414 |

**Supplementary Table 41. Top 10 Foldseek[37] results for sc-CC-7-LI against PDB database[39,40] filtered with 90% sequence identity.** * denotes *de novo* proteins.

| Accession code | Sequence identity, % | Probability | E-score | Length | RMSD, Å | TM-score |
|---|---|---|---|---|---|---|
| * 6xr1_A | 34.6 | 1 | 3.00E-13 | 360 | 5.97 | 0.698 |
| * 5cwq_A | 39 | 1 | 7.43E-07 | 219 | 1.41 | 0.944 |
| * 7rdr_A | 33.7 | 1 | 8.90E-09 | 345 | 13.62 | 0.470 |
| * 6xr2_C | 35.5 | 1 | 0.0003869 | 165 | 2.63 | 0.840 |
| * 6e9t_B | 28.1 | 1 | 0.01859 | 220 | 5.28 | 0.644 |
| 5a7d_C | 16.5 | 1 | 0.001074 | 329 | 18.07 | 0.366 |
| * 6e9y_A | 28.1 | 1 | 0.1699 | 220 | 5.45 | 0.631 |
| * 5cwn_A | 28.1 | 0.999 | 0.06663 | 219 | 7.0 | 0.510 |
| 7ep7_A | 13.8 | 0.993 | 0.01323 | 310 | 12.33 | 0.400 |
| 4jhr_B | 16.3 | 0.981 | 0.03232 | 292 | 9.75 | 0.390 |

**Supplementary Table 42. Top 10 Foldseek[37] results for sc-CC-8-58 against AlphaFold2-Swissprot database[5,38].**

| Accession code | Sequence identity, % | Probability | E-score | Length | RMSD, Å | TM-score |
|---|---|---|---|---|---|---|
| Q5ZIL9 | 11.2 | 0.997 | 0.004664 | 416 | 25.67 | 0.260 |
| P37630 | 8.8 | 0.956 | 0.08607 | 421 | 10.09 | 0.418 |
| Q8SYD0 | 13 | 0.872 | 0.1246 | 344 | 11.32 | 0.407 |
| Q9P7N6 | 13.8 | 0.747 | 0.2403 | 305 | 13.35 | 0.381 |
| Q7XZF5 | 11.5 | 0.663 | 1.242 | 411 | 8.84 | 0.411 |
| Q8BYG0 | 14 | 0.569 | 0.4829 | 211 | 13.11 | 0.349 |
| Q9UUG6 | 14.4 | 0.569 | 0.4448 | 381 | 13.07 | 0.359 |
| Q8NA31 | 8 | 0.442 | 0.7903 | 358 | 15.39 | 0.364 |
| Q9LD83 | 9.8 | 0.382 | 1.294 | 274 | 9.92 | 0.362 |
| A4IIL4 | 12.9 | 0.301 | 1.098 | 214 | 13.05 | 0.352 |

**Supplementary Table 43. Top 10 Foldseek[37] results for sc-CC-8-58 against PDB database[39,40] filtered with 90% sequence identity.** * denotes *de novo* proteins.

| Accession code | Sequence identity, % | Probability | E-score | Length | RMSD, Å | TM-score |
|---|---|---|---|---|---|---|
| * 6xr1_A | 33.1 | 1 | 4.10E-16 | 417 | 5.86 | 0.609 |
| * 5cwq_A | 42.7 | 1 | 4.52E-08 | 219 | 2.15 | 0.890 |
| * 7rdr_A | 28.3 | 1 | 1.39E-11 | 396 | 14.79 | 0.470 |
| * 6xr2_C | 40.8 | 1 | 0.0002047 | 168 | 1.93 | 0.878 |
| * 6e9t_B | 23.3 | 1 | 0.006442 | 226 | 4.24 | 0.729 |
| * 5cwn_A | 31.3 | 1 | 0.01012 | 219 | 6.04 | 0.564 |
| * 6e9y_A | 25.8 | 1 | 0.06163 | 246 | 7.41 | 0.667 |
| 3sf4_B | 15.2 | 1 | 0.000181 | 382 | 14.3 | 0.339 |
| 4a1s_B | 17.7 | 1 | 0.0002133 | 367 | 15.73 | 0.337 |
| 4jhr_B | 17.9 | 0.986 | 0.02602 | 294 | 8.72 | 0.393 |

#### 2.2 Supplementary figures

##### 2.2.1 Biophysical and structural characterization of *de novo* peptides

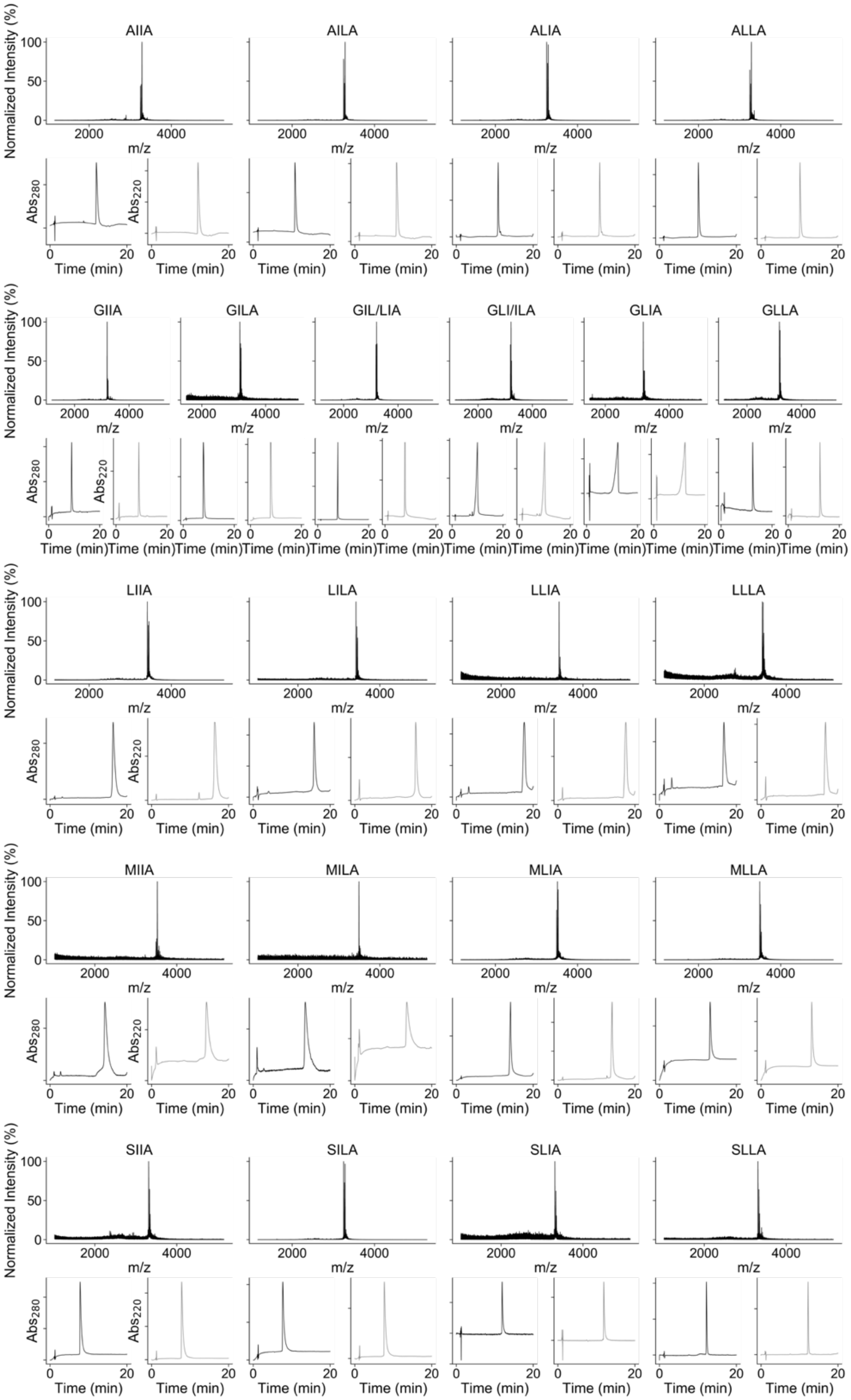

**Supplementary Figure 1. MALDI-TOF spectra (top) and analytical HPLC traces (bottom) of the *de novo* peptides designed for this study.** Sequences, calculated mass and observed mass for individual peptides can be found in Supplementary Table 1.

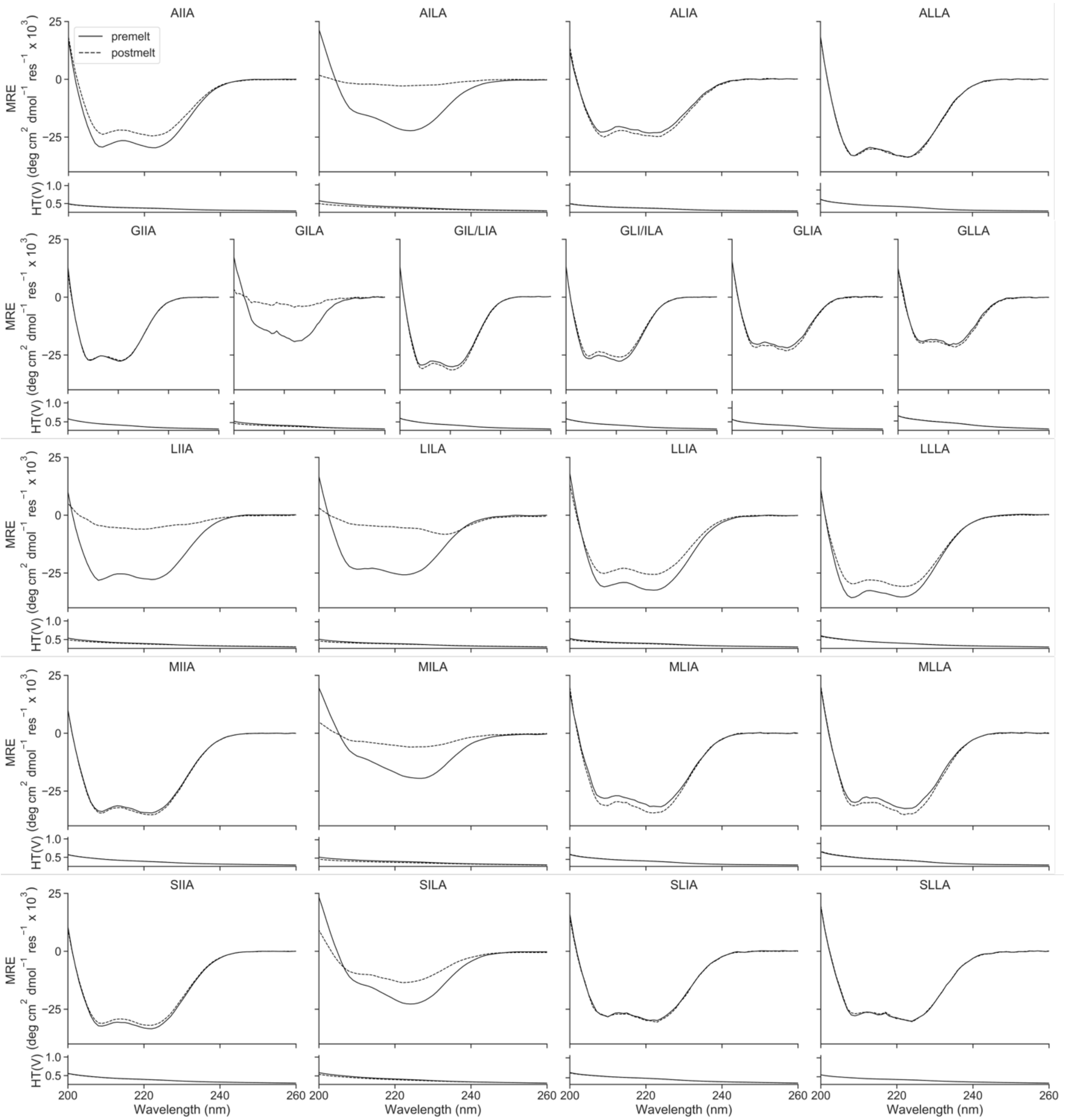

**Supplementary Figure 2. CD spectra of the *de novo* peptides designed for this study.** Sequences for individual peptides can be found in Supplementary Table 1. Summary of biophysical characterization can be found in Supplementary Table 3. Conditions: 50 μM peptide, PBS, pH 7.4, 5 °C. CD scans before thermal ramps up (solid line) and after thermal ramps down (dashed line).

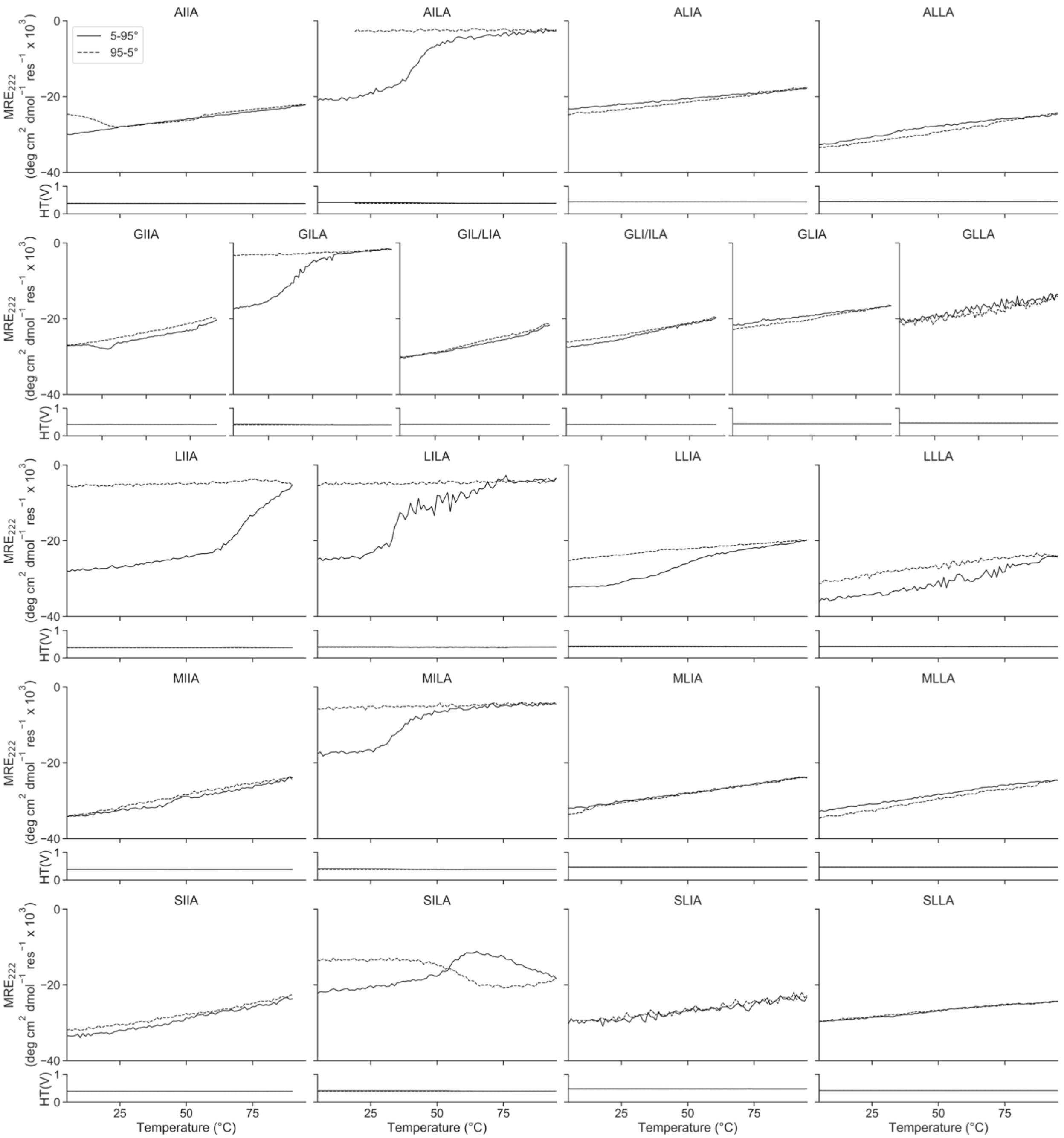

**Supplementary Figure 3. CD thermal-response profiles of the *de novo* peptides designed for this study.** Sequences for individual peptides can be found in Supplementary Table 1. Summary of biophysical characterization can be found in Supplementary Table 3. Conditions: 50 µM peptide, PBS, pH 7.4, 5 – 95 °C. CD scans before ramping up (solid line) and ramping down (dashed line).

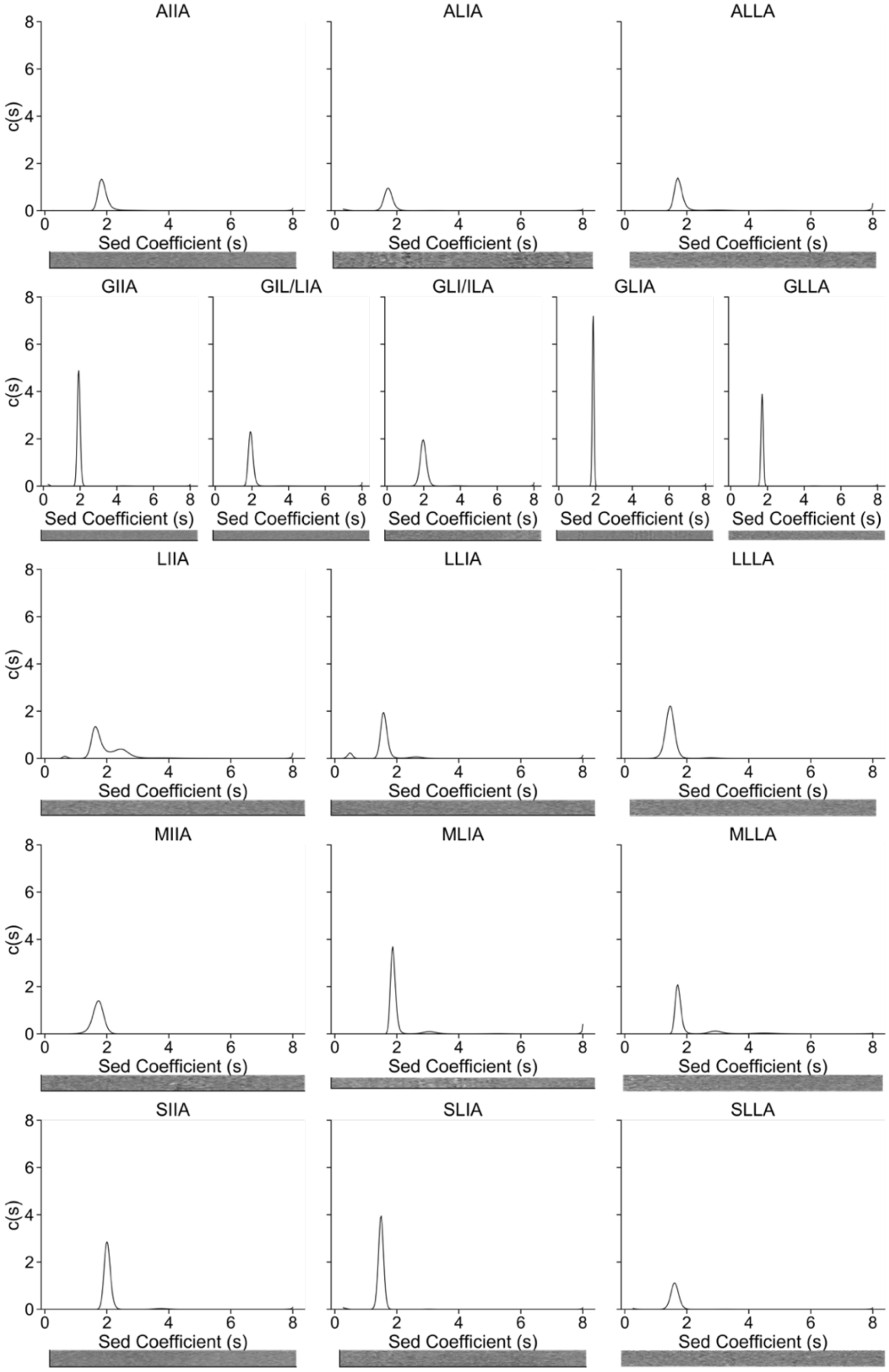

**Supplementary Figure 4. Sedimentation velocity (SV) AUC traces of the *de novo* peptides designed for this study.** Sequences for individual peptides can be found in Supplementary Table 1. Fit data for individual peptides can be found in Supplementary Table 2. Summary of biophysical characterization can be found in Supplementary Table 3. Residuals are shown as a bitmap below the fitted data. Conditions: 150 μM peptide, PBS, pH 7.4, 20 °C.

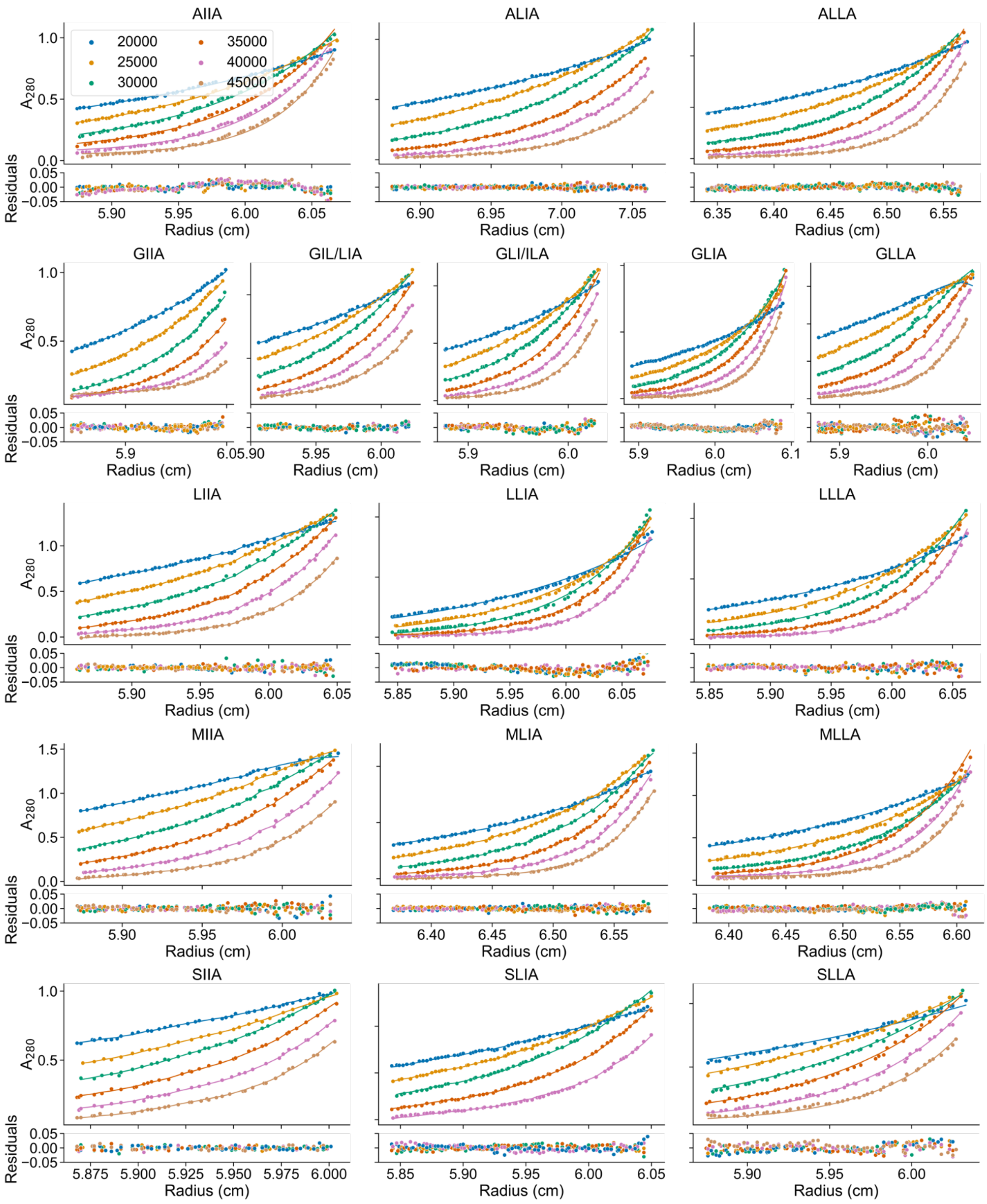

**Supplementary Figure 5. Sedimentation equilibrium (SE) AUC traces of the *de novo* peptides designed for this study.** Sequences for individual peptides can be found in Supplementary Table 1. Fit data for individual peptides can be found in Supplementary Table 2. Summary of biophysical characterization can be found in Supplementary Table 3. Data were collected at either 20 to 45 krpm at 5 krpm intervals or 24 to 48 krpm at 6 krpm intervals. Fitted single-ideal species model curves are overlaid in solid line, 95% confidence limits. Conditions: 70 mM peptide, PBS, pH 7.4, 20 °C.

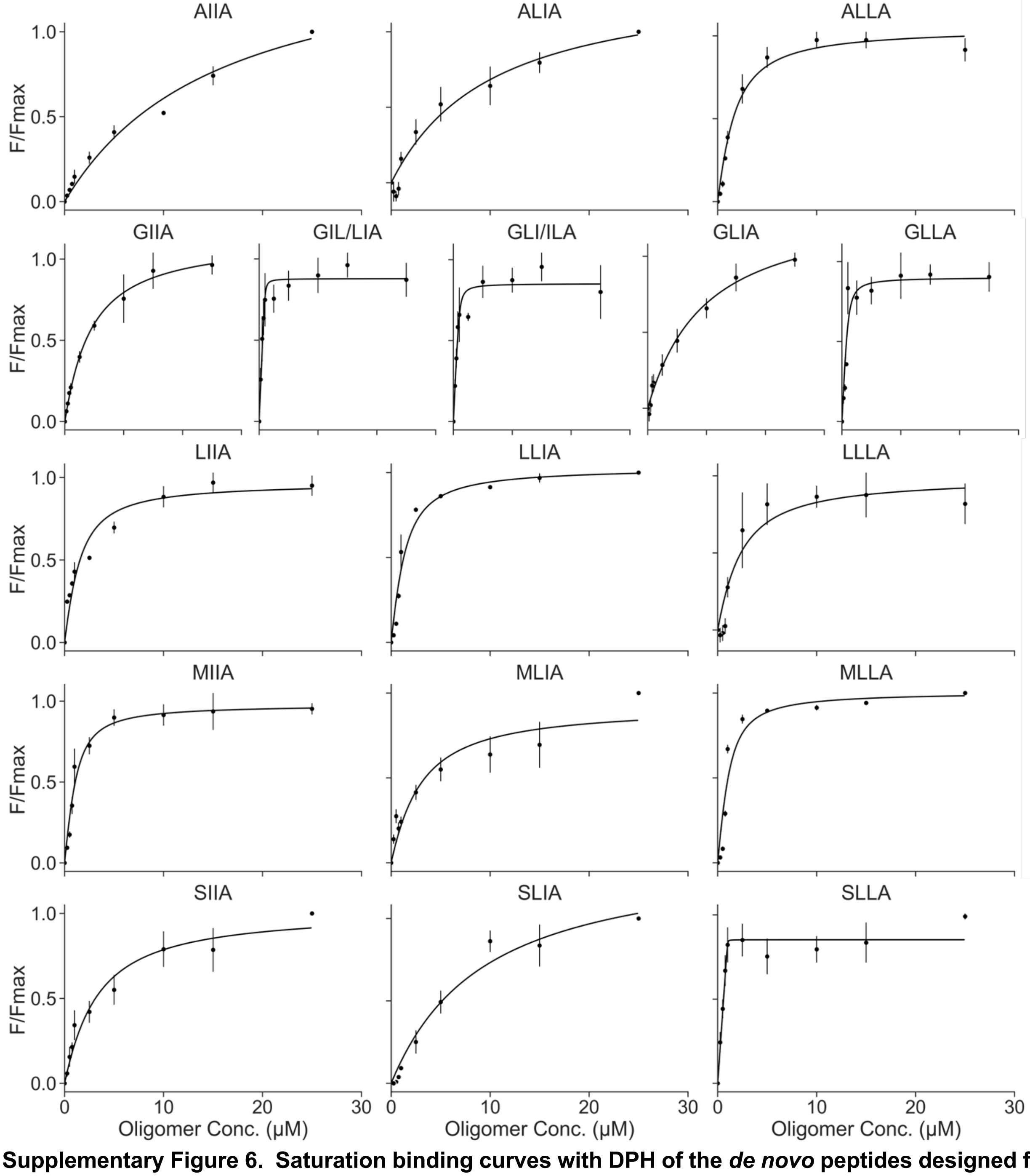

**Supplementary Figure 6. Saturation binding curves with DPH of the *de novo* peptides designed for this study.** Sequences for individual peptides can be found in Supplementary Table 1. Summary of biophysical characterization can be found in Supplementary Table 3. Conditions: Oligomer concentration 0 – 30 μM peptide assembly, PBS, pH 7.4, 20 °C. Final concentration 1 μM DPH (5% v/v DMSO).

**Supplementary Figure 7. X-ray crystal structure of apCC-Hex-ALIA collapsed bundle. a,** 1.6 Å X-ray crystal structure of antiparallel 6-helix ALIA collapsed bundle (PDB id, 8qaa). Coiled-coil regions identified by Socket2[41] (packing-cutoff 7.0 Å ) are coloured as chainbows from *N* (blue) to *C* (red) termini. **b,** Slice through the structure for a heptad repeat showing the knobs-into-holes packing: *a* knobs, red; *d*, green; *g*, magenta; *e*, cyan. Socket2[41] identifies KIH between two pairs of helices in the assembly. **c,** Heptad-repeat slice of apCC-Hex-ALIA AlphaFold-Multimer model[7] (magenta) aligned with its crystal structure (chainbows) (backbone RMSD = 2.531 Å). **Supplementary Figure 8. X-ray crystal structure of apCC-Oct-GLIA collapsed bundle**. **a,** 2.3 Å X-ray crystal structure of apCC-Oct-GLIA collapsed bundle (PDB id, 8qac). Coiled-coil regions identified by Socket2[41] (packing-cutoff 7.0 Å ) are coloured as chainbows from *N* (blue) to *C* (red) termini. **b,** Slice through the structure for a heptad repeat showing the knobs-into-holes packing: *a* knobs, red; *d*, green; *g*, magenta; *e*, cyan. Socket2[41] identifies KIH between two pairs of helices in the assembly. **c,** Heptad-repeat slice of apCC-Oct-GLIA AlphaFold-Multimer model[7] (magenta) aligned with its crystal structure (chainbows) (backbone RMSD = 3.482 Å).

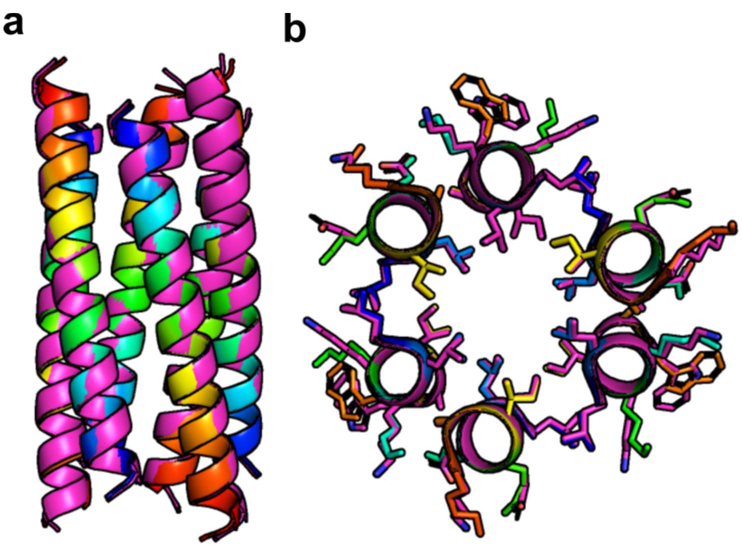

**Supplementary Figure 9. X-ray crystal structure of apCC-Hex-LLIA αHB.** **a,** 1.4 Å X-ray crystal structure of apCC-Hex-LLIA αHB (PDB id, 8qab). Coiled-coil regions identified by Socket2[41] (packing-cutoff 7.0 Å ) are coloured as chainbows from *N* (blue) to *C* (red) termini. Crystal structure is aligned to apCC-Hex-LLIA AlphaFold-Multimer model[7] (magenta) (backbone RMSD = 0.361 Å). **b,** Heptad-repeat slice of apCC-Hex-LLIA crystal structure and AlphaFold-Multimer model[7].

#### Computational analysis of sc-apCC-6

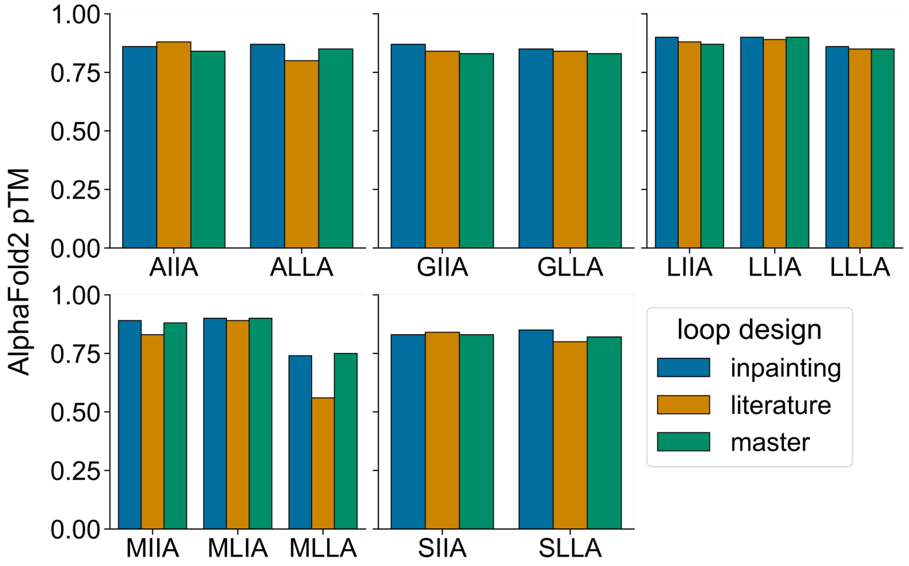

**Supplementary Figure 10. AlphaFold2 pTM for sc-ap-CC-6 designs.** AlphaFold2 predictions were performed using the ColabFold (version 1.3.0) using single-sequence mode and 3 recycle steps to generate 5 models. pTM of the top ranked model of each loop design method (bar colors) is compared for each possible g-a-d-e combination (columns).

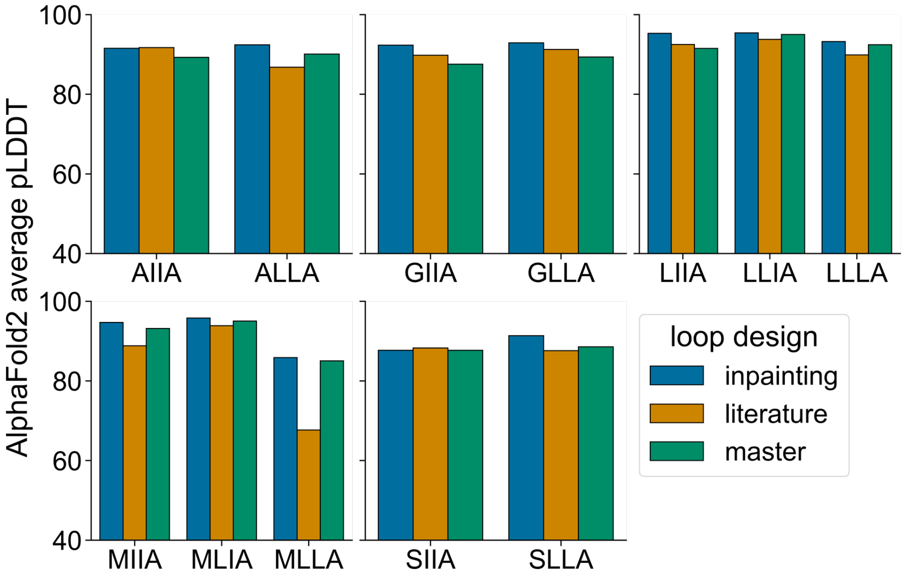

**Supplementary Figure 11. Average AlphaFold2 pLDDT for sc-apCC-6 designs.** AlphaFold2 predictions were performed using the ColabFold (version 1.3.0) using single-sequence mode and 3 recycle steps to generate 5 models. Average pLDDT of the top ranked model of each loop design method (bar colors) is compared for each possible g-a-d-e combination (columns).

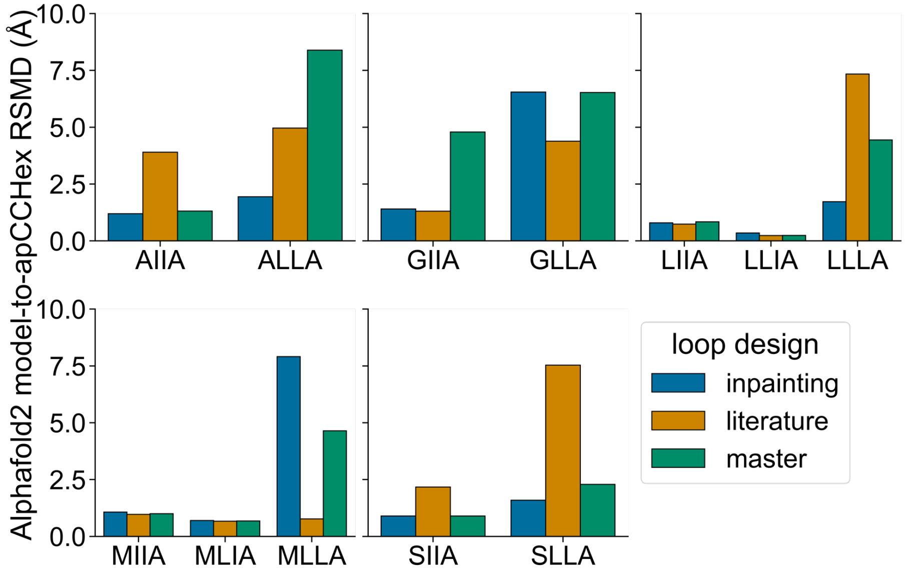

**Supplementary Figure 12. Calculated backbone RMSD (Å) between apCC-Hex-LLIA structure and AlphaFold2 models for sc-apCC-6 designs.** The top ranked AlphaFold2 models aligned to apCC-Hex crystal structure and RMSD calculated for backbone atoms in PyMOL. RMSD of each loop design method (bar colors) is compared for each possible g-a-d-e combination (columns).

#### Biophysical and structural characterization of sc-apCC-6

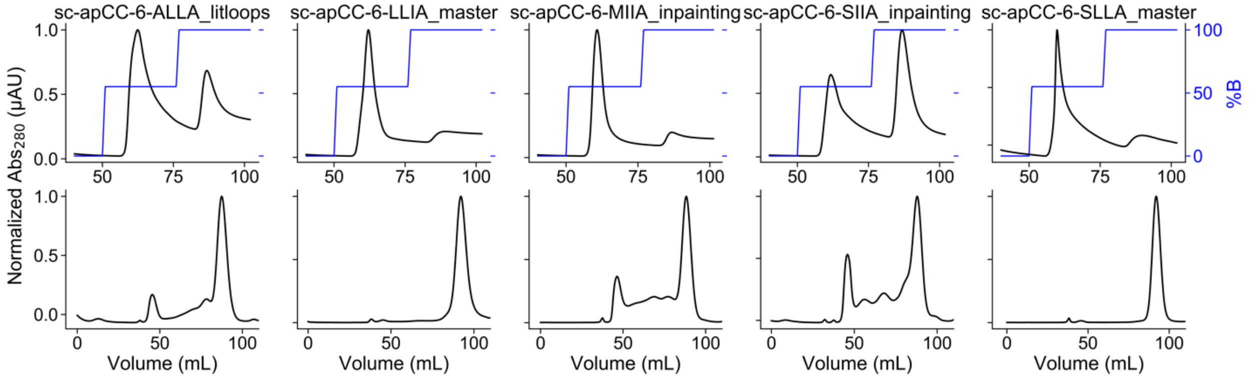

**Supplementary Figure 13. FPLC of sc-apCC-6 designs purification.** Sequences for individual proteins can be found in Supplementary Table 5. Top: sc-apCC-6 Ni-affinity chromatography. Fractions eluting at 55% B were pulled and concentrated then further purified by size exclusion chromatography. Conditions: Buffer A: 50 mM Tris, pH 7.4, 500 mM NaCl, 30 mM imidazole, B: 300 mM imidazole. Bottom: sc-apCC-6 variants size exclusion chromatography. Fractions eluting at 90 mL were pooled and concentrated to be analysed in further experiments. Conditions: 50 mM sodium phosphate, pH 7.4, 150 mM NaCl.

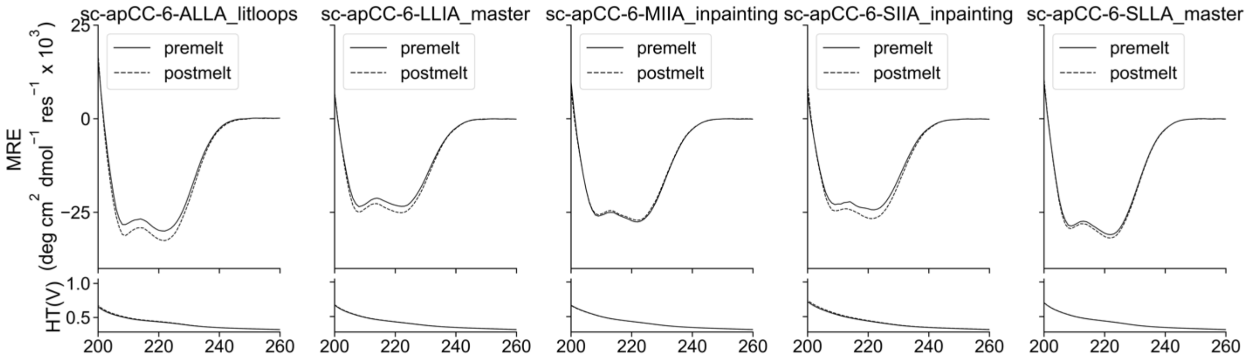

**Supplementary Figure 14. CD spectra of sc-apCC-6 proteins designed for this study.** Sequences for individual proteins can be found in Supplementary Table 5. Summary of biophysical data can be found in Supplementary Table 8. Conditions: 10 μM protein, 50 mM sodium phosphate, pH 7.4, 150 mM NaCl, 5 °C. CD scans before thermal ramps up (solid line) and after thermal ramps down (dashed line).

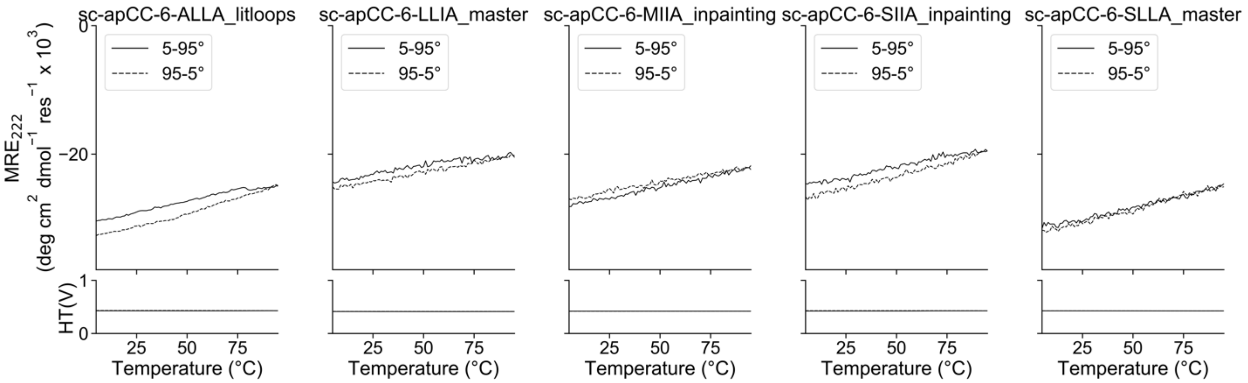

**Supplementary Figure 15. CD thermal-response profiles of sc-apCC-6 proteins designed for this study.** Sequences for individual proteins can be found in Supplementary Table 5. Summary of biophysical data can be found in Supplementary Table 8. Conditions: 10 μM protein, 50 mM sodium phosphate, pH 7.4, 150 mM NaCl, 5 – 95 °C. CD scans before ramping up (solid line) and ramping down (dashed line).

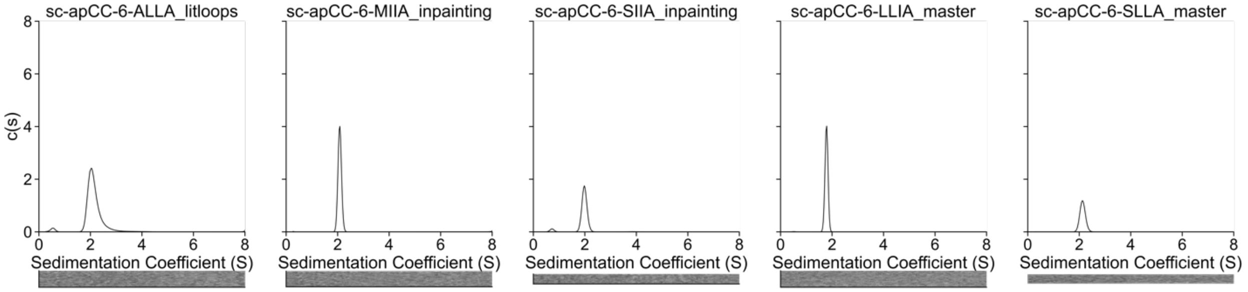

**Supplementary Figure 16. Sedimentation velocity (SV) AUC traces of sc-apCC-6 proteins designed for this study**. Sequences for individual proteins can be found in Supplementary Table 5. Fit data for individual proteins can be found in Supplementary Table 7. Summary of biophysical data can be found in Supplementary Table 8. Residuals are shown as a bitmap below the fitted data. Conditions: 15 μM protein, 50 mM sodium phosphate, pH 7.4, 150 mM NaCl, 20 °C.

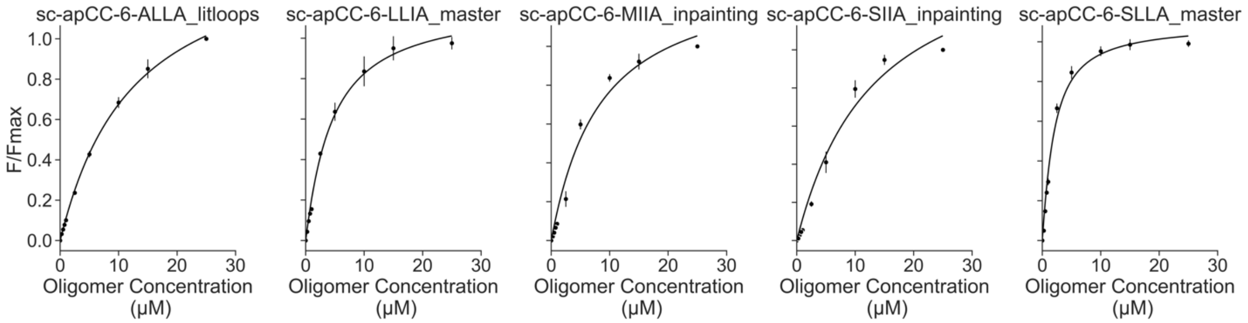

**Supplementary Figure 17. Saturation binding curves with DPH of sc-apCC-6 proteins designed for this study.** Sequences for individual proteins can be found in Supplementary Table 5. Summary of biophysical data can be found in Supplementary Table 8. Conditions: Oligomer concentration 0 – 30 μM protein, 50 mM sodium phosphate, pH 7.4, 150 mM NaCl, 20 °C. Final concentration 1 μM DPH (5% v/v DMSO).

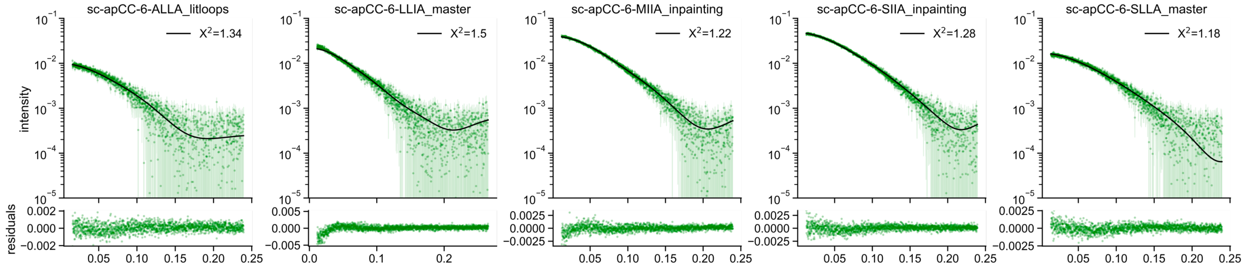

**Supplementary Figure 18. FoXs SAXS profile with fit (top) and residuals (bottom) to AlphaFold2 model of sc-apCC-6 proteins designed for this study.** Sequences for individual proteins can be found in Supplementary Table 5. Summary of biophysical data can be found in Supplementary Table 8. SAXS fit metrics can be found in Supplementary Table 9. Conditions: 10 mg/mL, 50 mM sodium phosphate, pH 7.4, 150 mM NaCl.

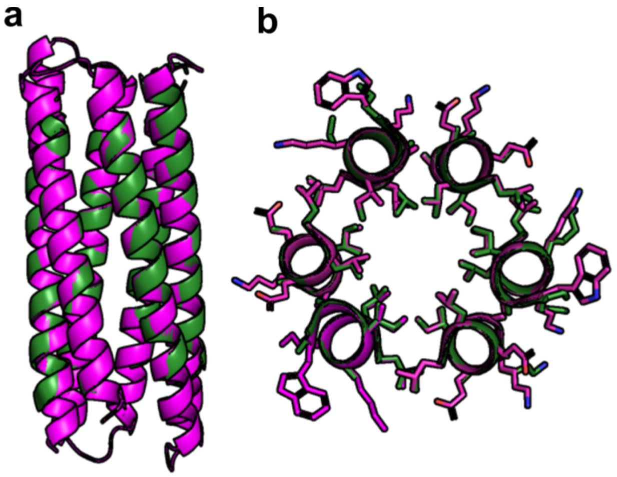

**Supplementary Figure 19. X-ray crystal structure of sc-apCC-6-LLIA**. **a,** 2.25 Å X-ray crystal structure of sc-apCC-6-LLIA (PDB id, 8qad). Crystal structure (green) is aligned to sc-apCC-6-LLIA_master AlphaFold2 model (magenta, backbone RMSD = 0.269 Å). **b,** Heptad-repeat slice of sc-apCC-6-LLIA crystal structure and AlphaFold2 model[5,6].

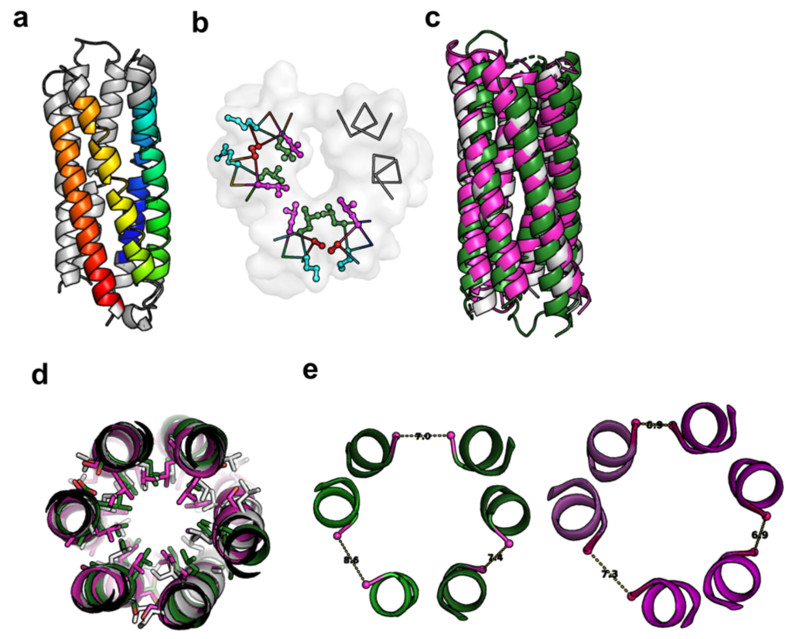

**Supplementary Figure 20. X-ray crystal structure of sc-apCC-6-SLLA. a,** 1.90 Å X-ray crystal structure of sc-apCC-6-SLLA (PDB id, 8qae). Coiled-coil regions identified by Socket2[41] (packing-cutoff 7.5 Å) are coloured as chainbows from *N* (blue) to *C* (red) termini. **b,** Slice through the structure for a heptad repeat showing the knobs-into-holes packing: *a* knobs, red; *d*, green; *g*, magenta; *e*, cyan. Socket2[41] identifies KIH between pairs of helices in the assembly. **c-d,** Crystal structure of sc-apCC-6-SLLA (green) aligned to AlphaFold2 model[5,6] of sc-apCC-6-SLLA_master (magenta) and apCC-Hex-LLIA (grey, backbone RMSD = 14.144, 7.913 Å). **(d),** Heptad-repeat slice of sc-apCC-6-SLLA crystal structure AlphaFold2 model[5,6], and apCC-Hex-LLIA. **e,** Ser-Ser (g-g) helical interface in sc-apCC-6-SLLA structure (green) differs at one interface by 1 Å relative to the average of the other interfaces. In the AlphaFold2 model (magenta), the interhelical distances are the same throughout.

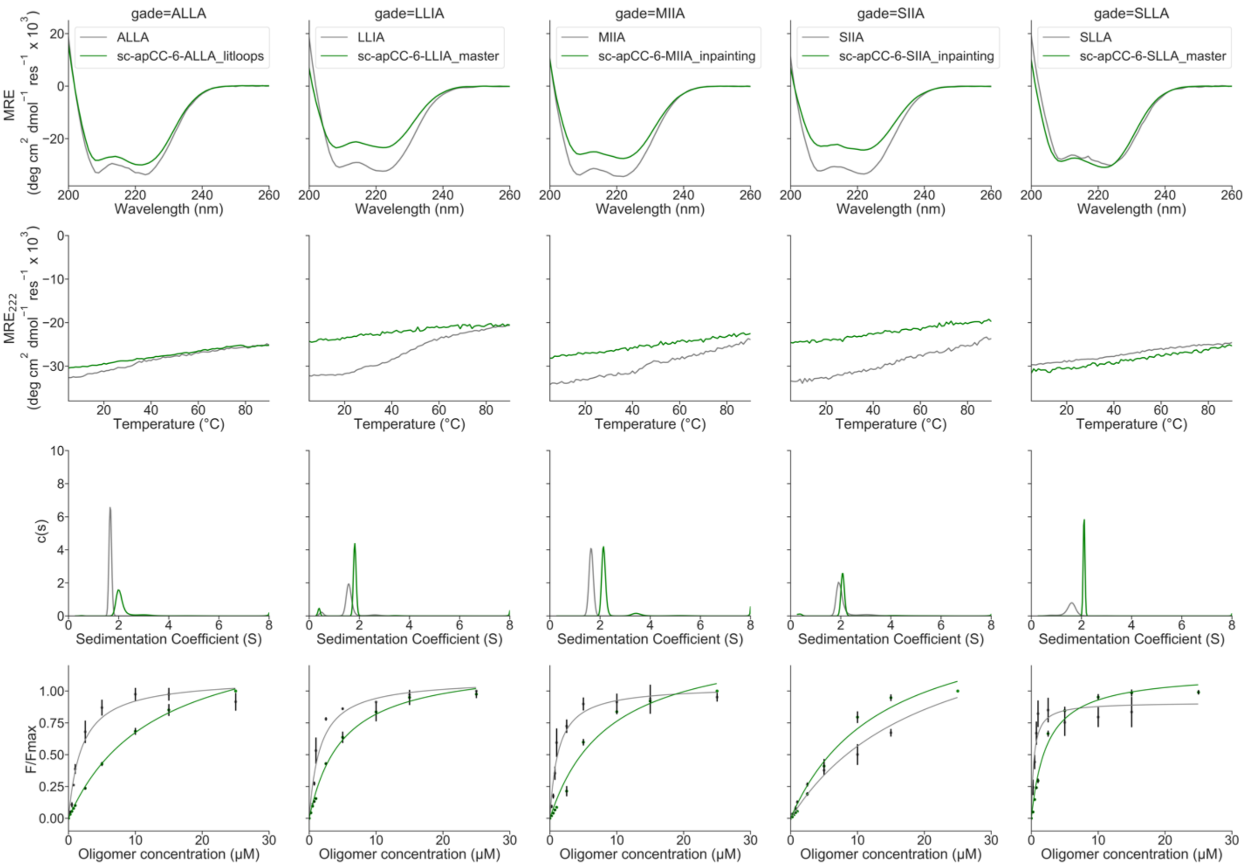

**Supplementary Figure 21. Direct peptide-to-protein biophysical characterization comparisons for sc-apCC-6 designs**. Peptide data are shown in grey, protein in green. Summary of biophysical data can be found in Supplementary Tables 3 and 8. Conditions: same as mentioned above for respective experiment.

##### 2.2.4 Computational analysis of sc-CC-7

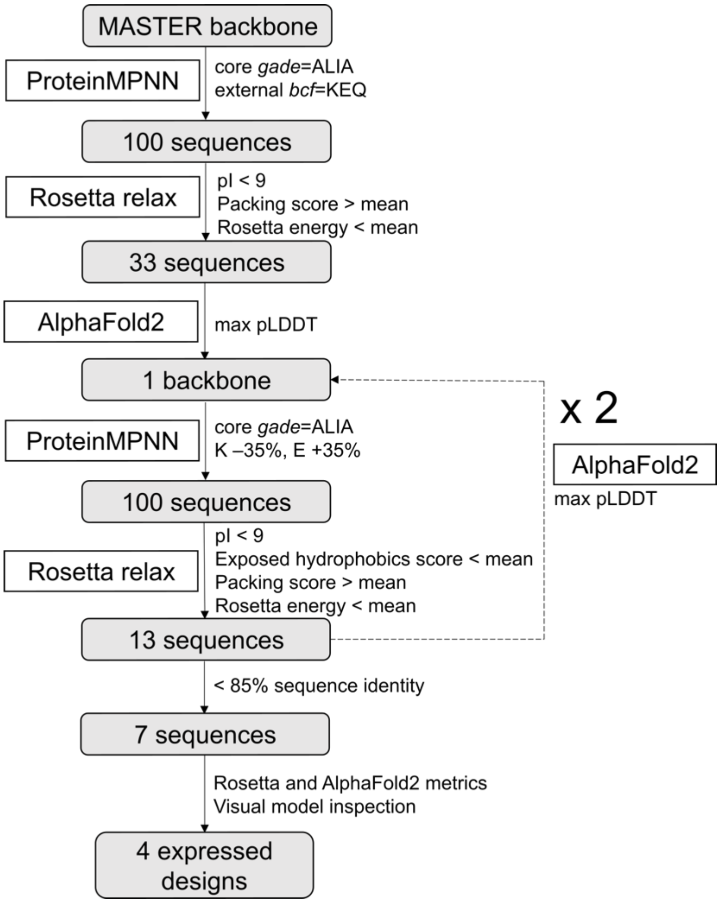

**Supplementary Figure 22. ProteinMPNN/AlphaFold2 pipeline used for sc-CC-7 design.** The same pipeline was used for sc-CC-5, sc-CC-6 and sc-CC-8. Only the KE bias was varied to maintain the desired pI distribution. K -40%, E +40% for sc-CC-6 and sc-CC-8, K -20%, E +20% for sc-CC-5. Sc-CC-5 converged after 5 iterations, sc-CC-6 after 2 and sc-CC-8 after 3 iterations.

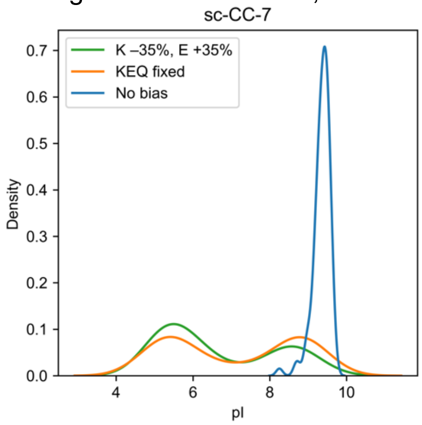

**Supplementary Figure 23. pI distribution after ProteinMPNN design with different sequence constraints for sc-CC-7.**

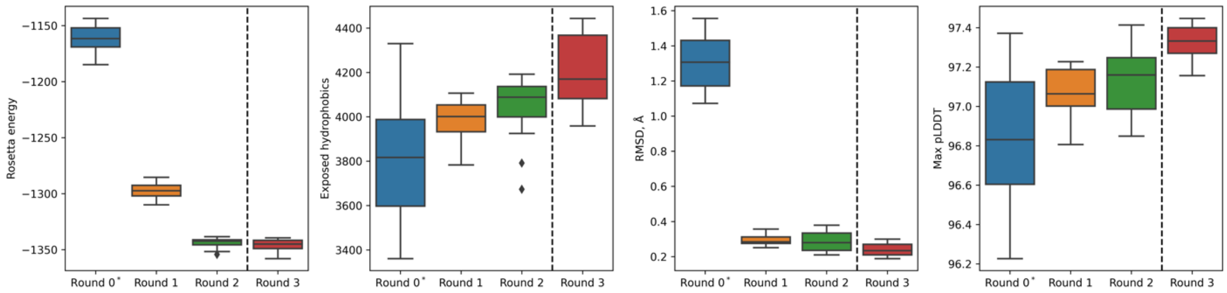

**Supplementary Figure 24. Design metrics after filtering at each iteration for sc-CC-7.** RMSD is calculated using the input backbone at each iteration. *External b-c-f fixed to Lys-Glu-Gln. Dotted lines show where the iterative design was stopped due to Rosetta energy convergence.

#### 2.2.5 Biophysical and structural characterization of sc-CC-7

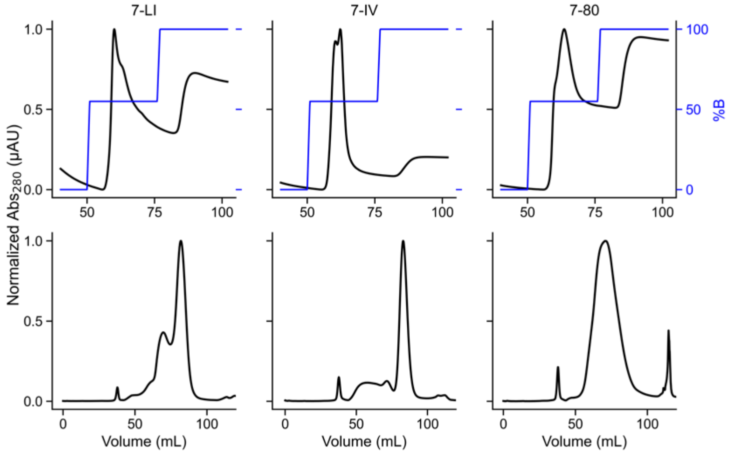

**Supplementary Figure 25. FPLC of sc-CC-7 variants purification.** Sequences for individual proteins can be found in Supplementary Table 10. Top: Ni-affinity chromatography. Fractions eluting at 55% B were pulled and concentrated then further purified by size exclusion chromatography. Conditions: Buffer A: 50 mM Tris, pH 7.4, 150 mM NaCl, 30 mM imidazole, B: 300 mM imidazole. Bottom: Size exclusion chromatography. Fractions eluting at 85 mL were pooled and concentrated to be analysed in further experiments. Conditions: 50 mM sodium phosphate, pH 7.4, 150 mM NaCl.

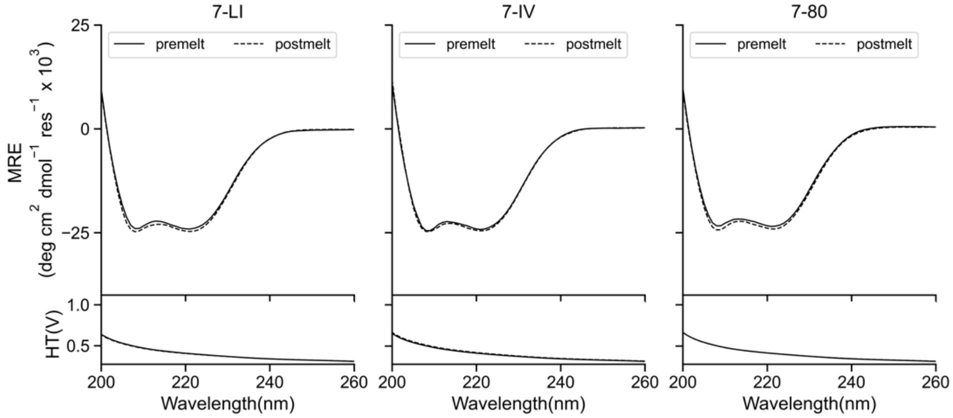

**Supplementary Figure 26. CD spectra of sc-CC-7 proteins designed for this study.** Sequences for individual proteins can be found in Supplementary Table 10. Summary of biophysical characterization can be found in Supplementary Table 13. Conditions: 5 µM protein, 50 mM sodium phosphate, pH 7.4, 150 mM NaCl, 5 °C. CD scans before thermal ramps up (solid line) and after thermal ramps down (dashed line).

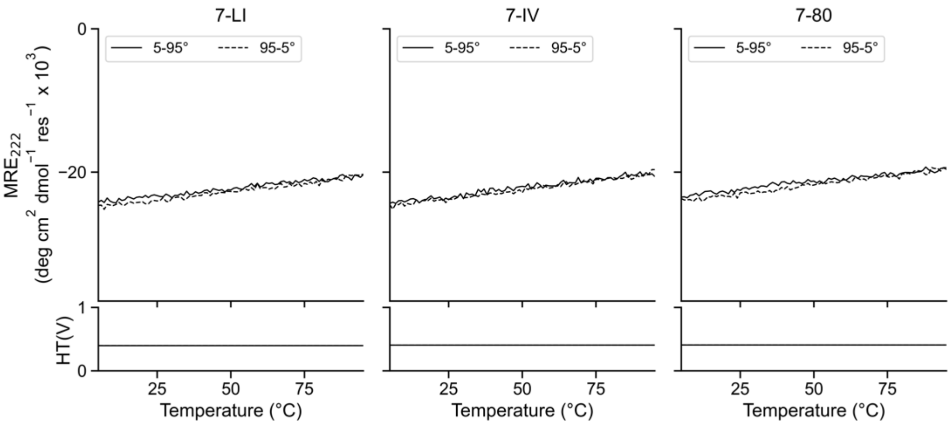

**Supplementary Figure 27. CD thermal-response profiles of sc-CC-7 proteins designed for this study.** Sequences for individual proteins can be found in Supplementary Table 10. Summary of biophysical characterization can be found in Supplementary Table 13. Conditions: 5 µM protein, 50 mM sodium phosphate, pH 7.4, 150 mM NaCl, 5 – 95 °C. CD scans before ramping up (solid line) and ramping down (dashed line).

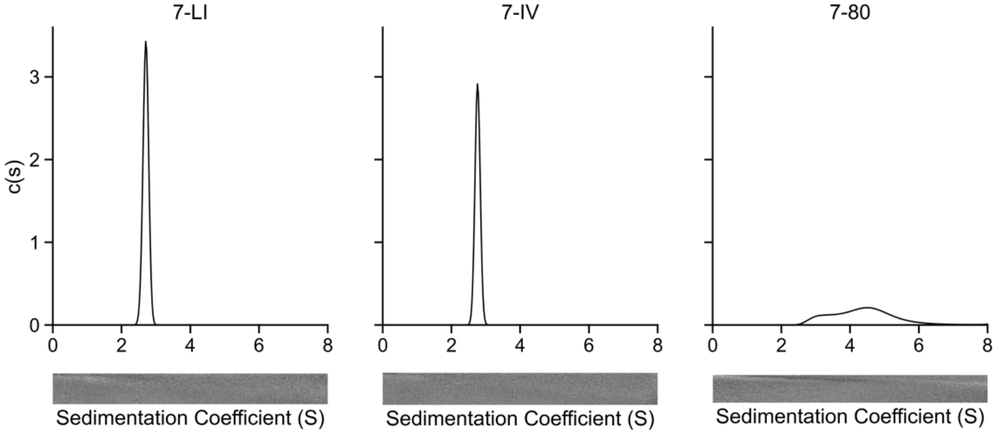

**Supplementary Figure 28. Sedimentation velocity (SV) AUC traces of sc-CC-7 proteins designed for this study**. Sequences for individual proteins can be found in Supplementary Table 10. Summary of fit data can be found in Supplementary Table 12. Summary of biophysical characterization can be found in Supplementary Table 13. Residuals are shown as a bitmap below the fitted data. Conditions: 25 µM protein, 50 mM sodium phosphate, pH 7.4, 150 mM NaCl, 20 °C.

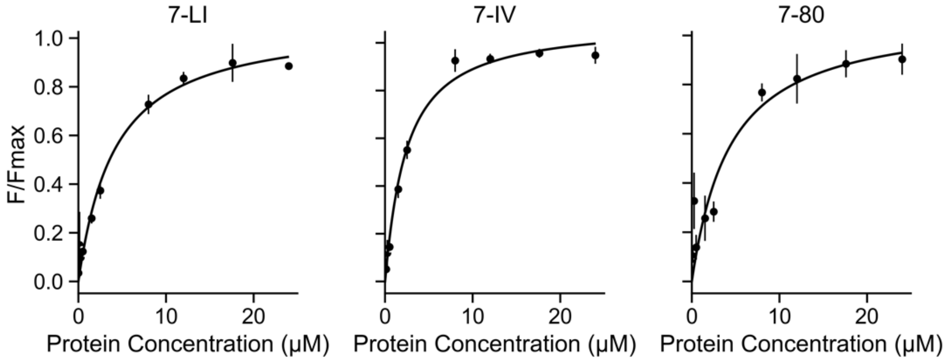

**Supplementary Figure 29. Saturation binding curves with DPH of sc-CC-7 proteins designed for this study**. Sequences for individual proteins can be found in Supplementary Table 10. Summary of biophysical characterization can be found in Supplementary Table 13. Conditions: Oligomer concentration 0 – 24 μM protein, 50 mM sodium phosphate, pH 7.4, 150 mM NaCl, 20 °C. Final concentration 0.5 μM DPH (5% v/v DMSO).

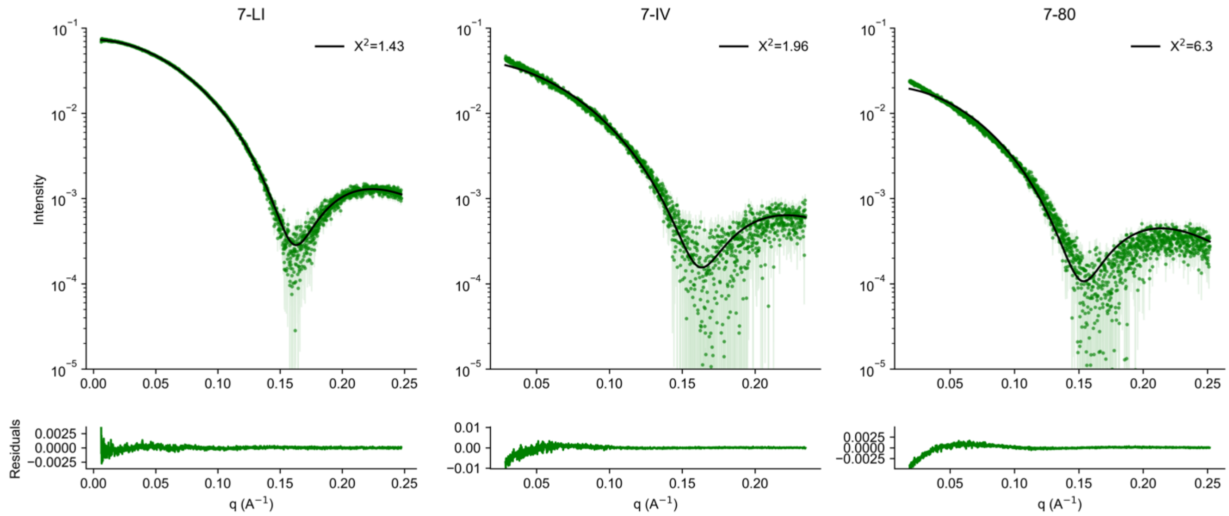

**Supplementary Figure 30. FoXs SAXS profile with fit (top) and residuals (bottom) to AlphaFold2 model of sc-CC-7 proteins designed for this study.** Sequences for individual proteins can be found in Supplementary Table 10. Summary of biophysical characterization can be found in Supplementary Table 13. Summary of SAXS metrics can be found in Supplementary Table 14. Conditions: 10 mg/mL, 50 mM sodium phosphate, pH 7.4, 150 mM NaCl. * sc-CC-7-80 aggregated at 10 mg/mL, which is observed in the SEC-SAXS collection and AUC.

##### 2.2.6 Computational analysis of sc-apCC-8

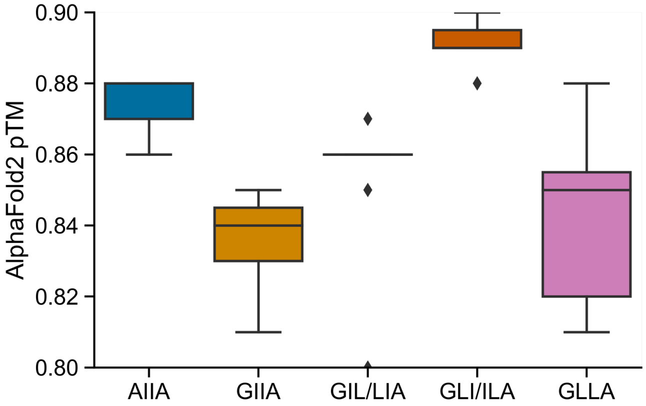

**Supplementary Figure 31. AlphaFold2 pTM for sc-apCC-8 designs.** The 10 ProteinMPNN sequences for each sc-ap-CC-8 starting scaffold were modelled using AlphaFold2. AlphaFold2 predictions were performed using the ColabFold (version 1.3.0) using single-sequence mode and 3 recycle steps to generate 5 models. AlphaFold2 pTM distribution for the top ranked model for each ProteinMPNN sequence was compared.

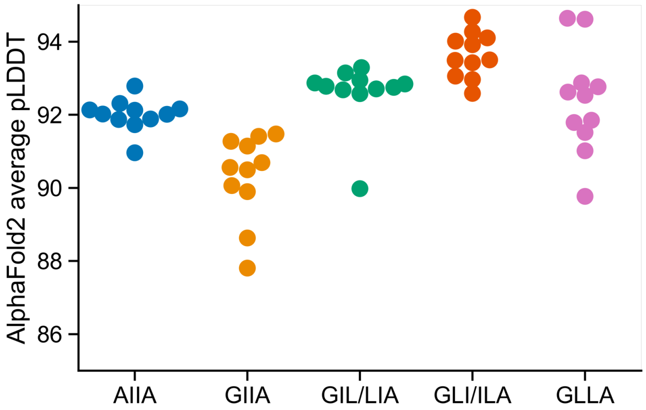

**Supplementary Figure 32. Average AlphaFold2 pLDDT for sc-apCC-8 designs.** The 10 ProteinMPNN sequences for each sc-ap-CC-8 starting scaffold were modelled using AlphaFold2. AlphaFold2 predictions were performed using the ColabFold (version 1.3.0) using single-sequence mode and 3 recycle steps to generate 5 models. AlphaFold2 average pLDDT distribution for the top ranked model for each ProteinMPNN sequence was compared.

#### 2.2.7 Biophysical and structural characterization of sc-apCC-8

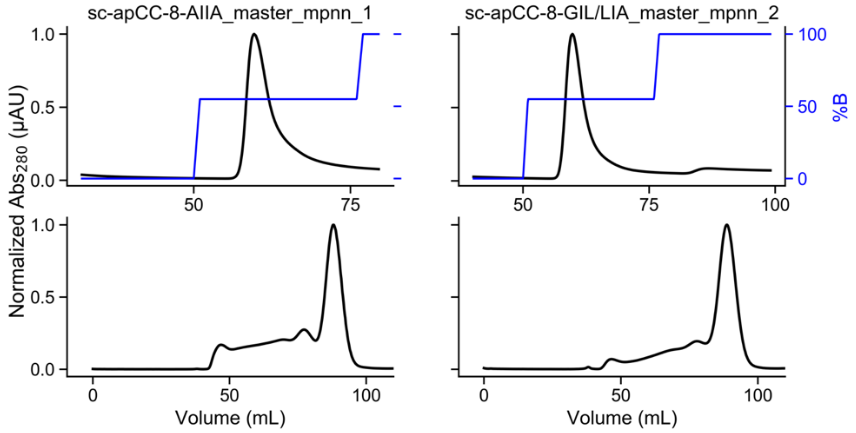

**Supplementary Figure 33. FPLC of sc-apCC-8 designs purification.** Sequences for individual proteins can be found in Supplementary Table 15. Top: sc-apCC-8 Ni-affinity chromatography. Fractions eluting at 55% B were pulled and concentrated then further purified by size exclusion chromatography. Conditions: Buffer A: 50 mM Tris, pH 7.4, 500 mM NaCl, 30 mM imidazole, B: 300 mM imidazole. Bottom: sc-apCC-8 variants size exclusion chromatography. Fractions eluting at 90 mL were pooled and concentrated to be analysed in further experiments. Conditions: 50 mM sodium phosphate, pH 7.4, 150 mM NaCl.

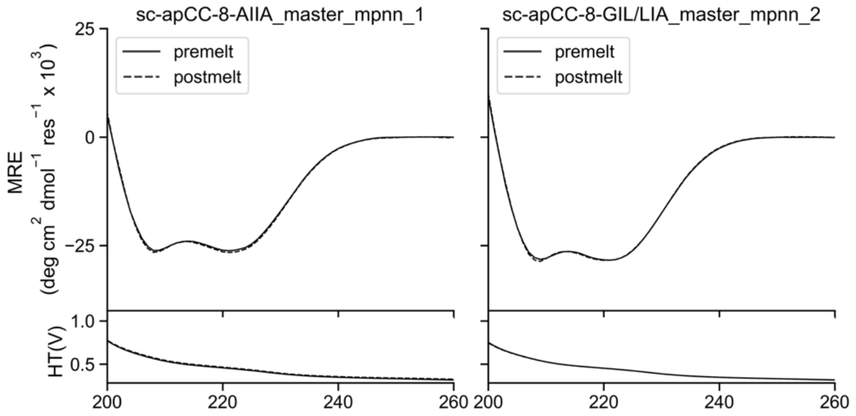

**Supplementary Figure 34. CD spectra of sc-apCC-8 proteins designed for this study.** Sequences for individual proteins can be found in Supplementary Table 15. Summary of biophysical data can be found in Supplementary Table 18. Conditions: 10 µM protein, 50 mM sodium phosphate, pH 7.4, 150 mM NaCl, 5

°C. CD scans before thermal ramps up (solid line) and after thermal ramps down (dashed line).

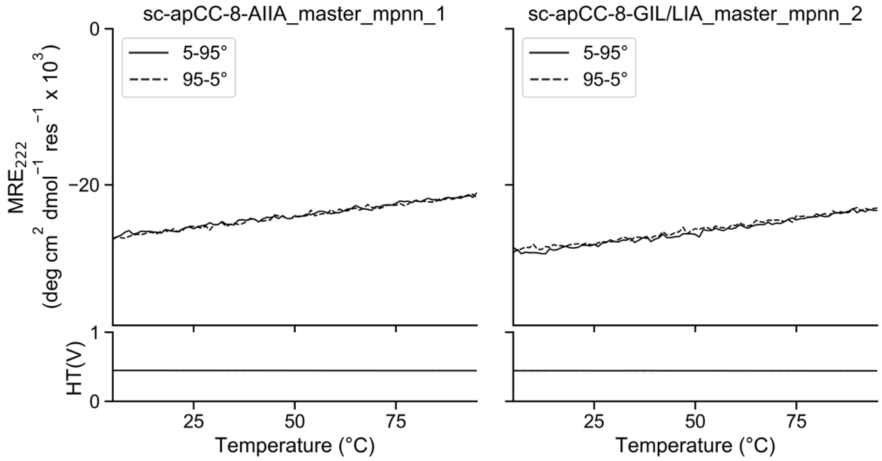

**Supplementary Figure 35. CD thermal-response profiles of sc-apCC-8 proteins designed for this study.** Sequences for individual proteins can be found in Supplementary Table 15. Summary of biophysical data can be found in Supplementary Table 18. Conditions: 10 μM protein, 50 mM sodium phosphate, pH 7.4, 150 mM NaCl, , 5 – 95 °C. CD scans before ramping up (solid line) and ramping down (dashed line).

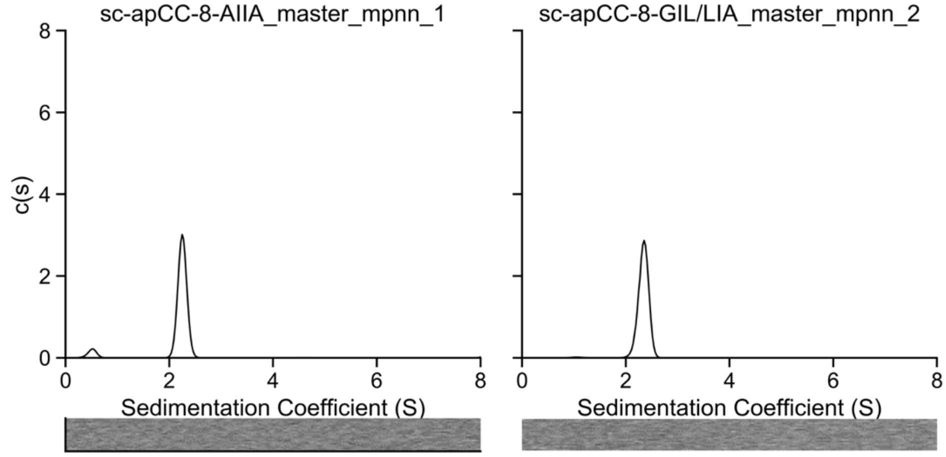

**Supplementary Figure 36. Sedimentation velocity (SV) AUC traces of sc-apCC-8 proteins designed for this study.** Sequences for individual proteins can be found in Supplementary Table 15. Fit data for individual proteins can be found in Supplementary Table 17. Summary of biophysical data can be found in Supplementary Table 18. Residuals are shown as a bitmap below the fitted data. Conditions: 15 μM protein, 50 mM sodium phosphate, pH 7.4, 150 mM NaCl, 20 °C.

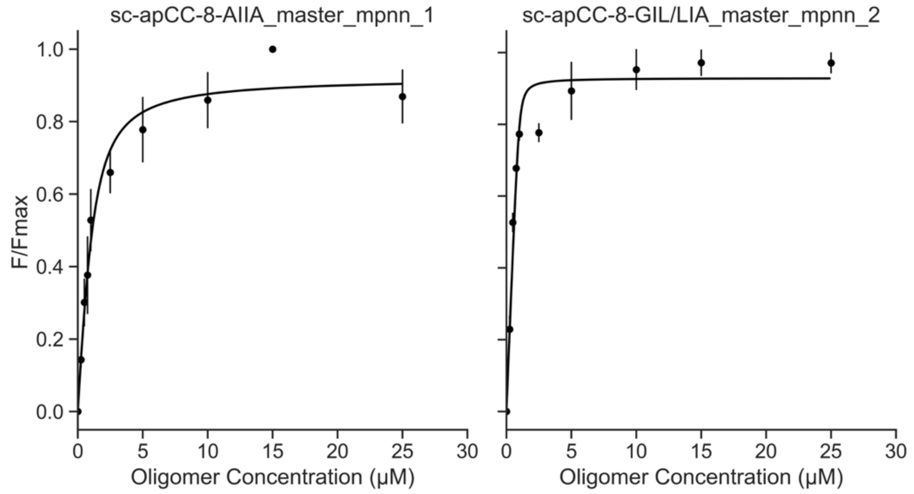

**Supplementary Figure 37. Saturation binding curves with DPH of sc-apCC-8 proteins designed for this study.** Sequences for individual proteins can be found in Supplementary Table 15. Summary of biophysical data can be found in Supplementary Table 18. Conditions: Oligomer concentration 0 – 30 µM protein, 50 mM sodium phosphate, pH 7.4, 150 mM NaCl, 20 °C. Final concentration 1 µM DPH (5% v/v

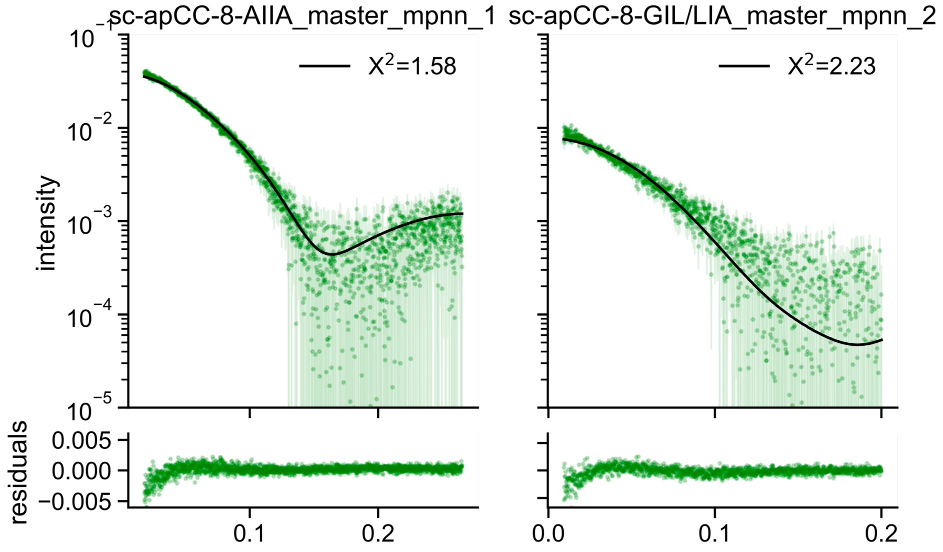

DMSO).

**Supplementary Figure 38. FoXs SAXS profile with fit (top) and residuals (bottom) to AlphaFold2 model of sc-apCC-8 proteins designed for this study.** Sequences for individual proteins can be found in Supplementary Table 15. Summary of biophysical data can be found in Supplementary Table 18. Summary of SAXS metrics can be found in Supplementary Table 19. Conditions: 10 mg/mL, 50 mM sodium phosphate, pH 7.4, 150 mM NaCl.

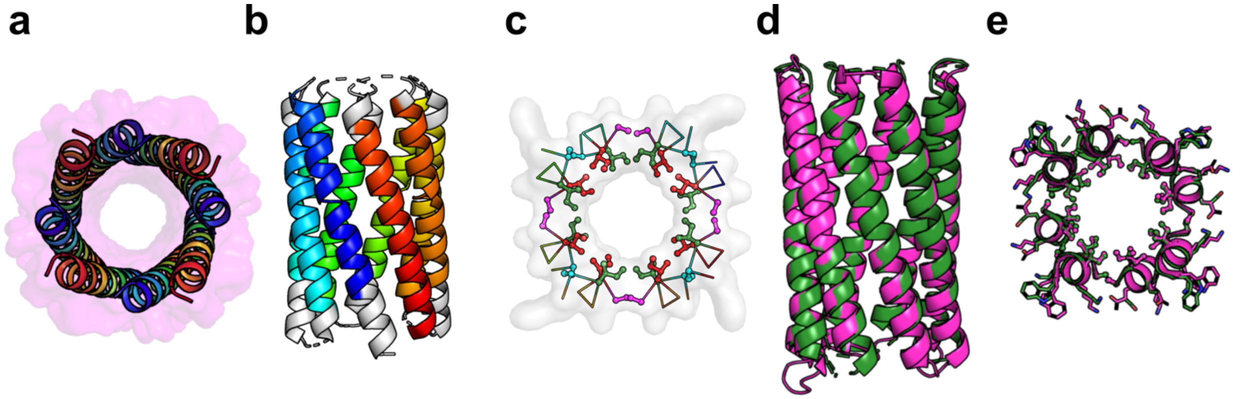

**Supplementary Figure 39. X-ray crystal structure of sc-apCC-8**. **a,** Starting AlphaFold-Multimer[7] model for apCC-Oct-AIIA that was used to rationally seed computational design of sc-apCC-8-AIIA (chains colored in chainbows from *N* (blue) to *C* (red) termini, surface in magenta). **b-c,** 2.0 Å X-ray crystal structure of sc-apCC-8-AIIA (PDB id, 8qaf). Coiled-coil regions identified by Socket2[41] (packing-cutoff 7.5 Å ) are coloured as chainbows. **(c),** Slice through the structure for a heptad repeat showing the knobs-into-holes packing: *a* knobs, red; *d*, green; *g*, magenta; *e*, cyan. Socket2[41] identifies KIH for the full αHB assembly. **d-e,** Alignment of sc-apCC-8-AIIA crystal structure (green) and AlphaFold2 (magenta) model[5,6] (backbone RMSD = 0.413 Å). **(e)** Heptad-repeat slice through crystal structure and its respective models, colors same as in panel **d**.

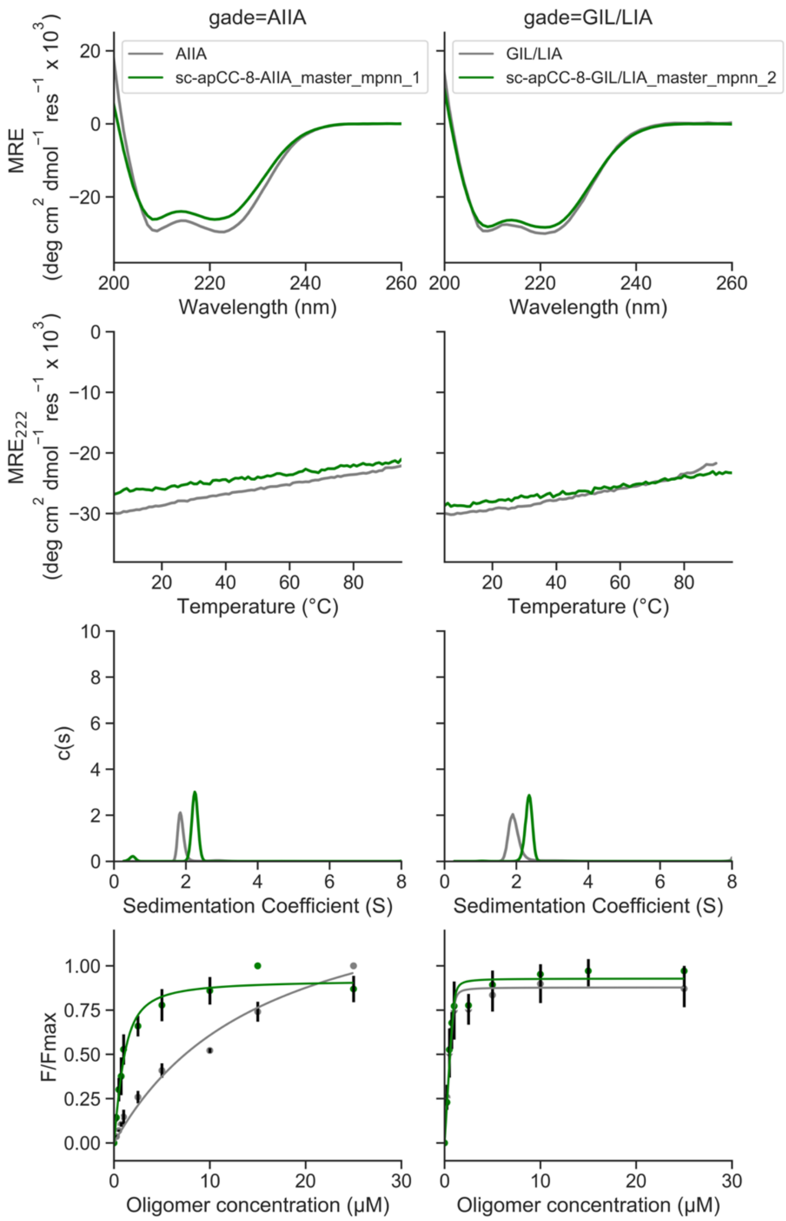

**Supplementary Figure 40. Direct peptide to protein biophysical characterization comparisons for sc-apCC-8 designs.** Peptide data are shown in grey, protein in green. Summary of biophysical data can be found in Supplementary Tables 3 and 18. Conditions: same as mentioned above for respective experiment.

##### 2.2.8 Computational analysis of sc-CC-5, 6, and 8

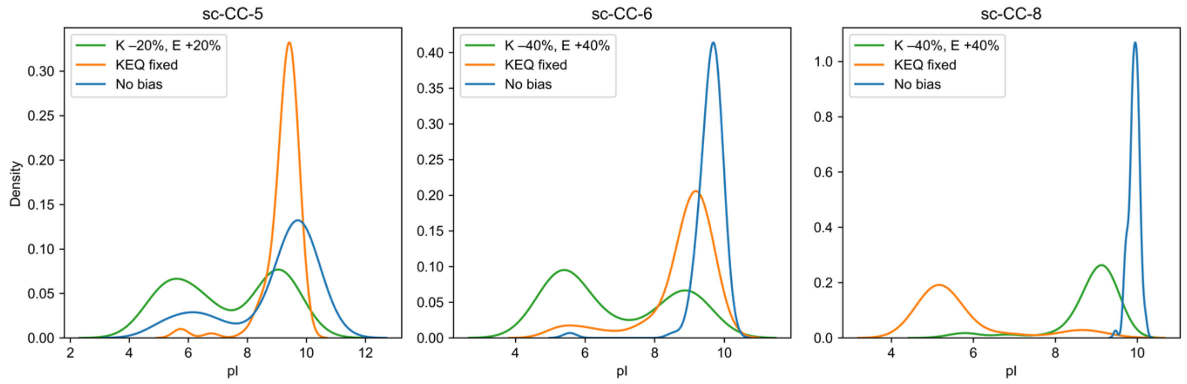

**Supplementary Figure 41. pI distribution after ProteinMPNN design with different sequence constraints.**

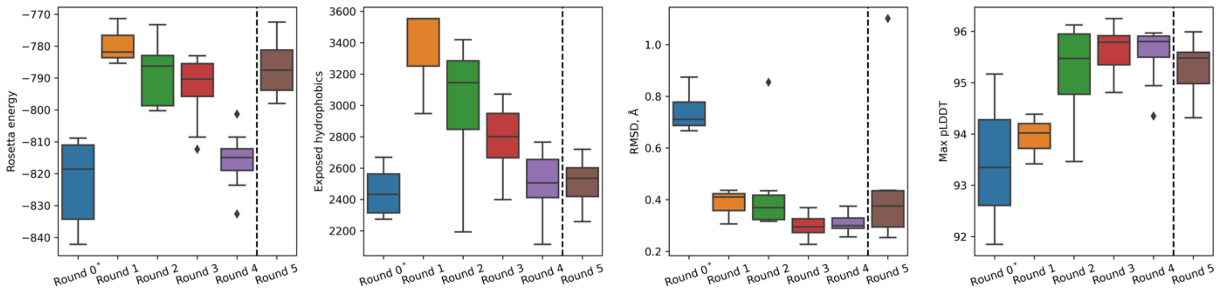

**Supplementary Figure 42. Design metrics after filtering at each iteration for sc-CC-5.** RMSD is calculated using the input backbone at each iteration. *External b-c-f fixed to Lys-Glu-Gln. Dotted lines show where the iterative design was stopped due to Rosetta energy convergence.

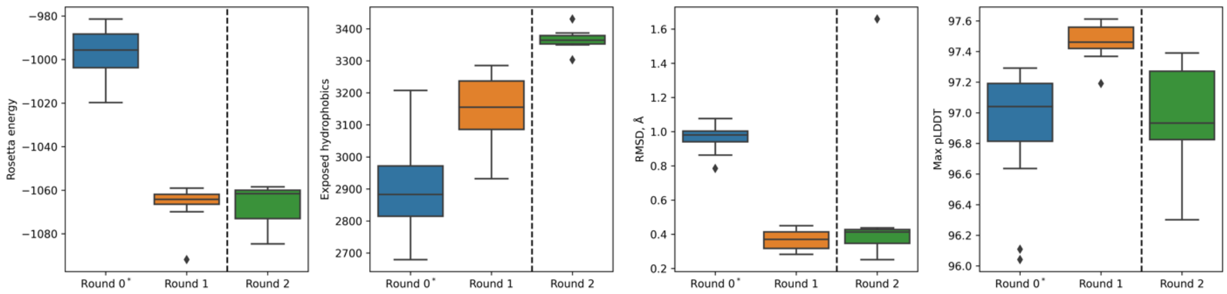

**Supplementary Figure 43. Design metrics after filtering at each iteration for sc-CC-6.** RMSD is calculated using the input backbone at each iteration. *External b-c-f fixed to Lys-Glu-Gln. Dotted lines show where the iterative design was stopped due to Rosetta energy convergence.

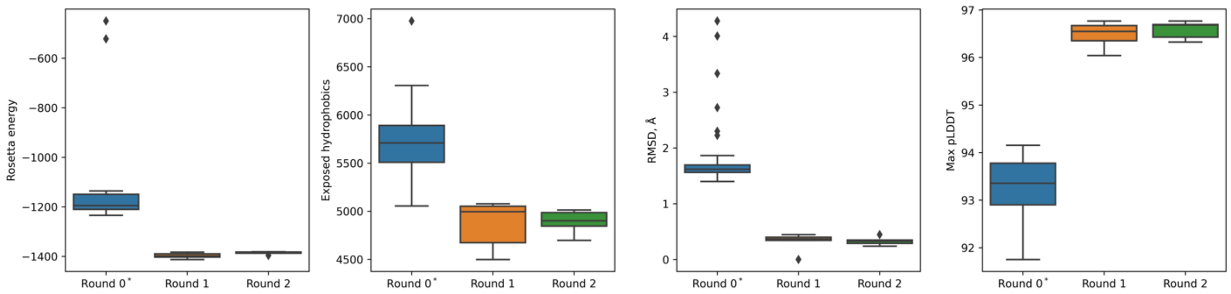

**Supplementary Figure 44. Design metrics after filtering at each iteration for sc-CC-8.** RMSD is calculated using the input backbone at each iteration. *External b-c-f fixed to Lys-Glu-Gln. 2 sequences were experimentally characterised from rounds 1 and 2.

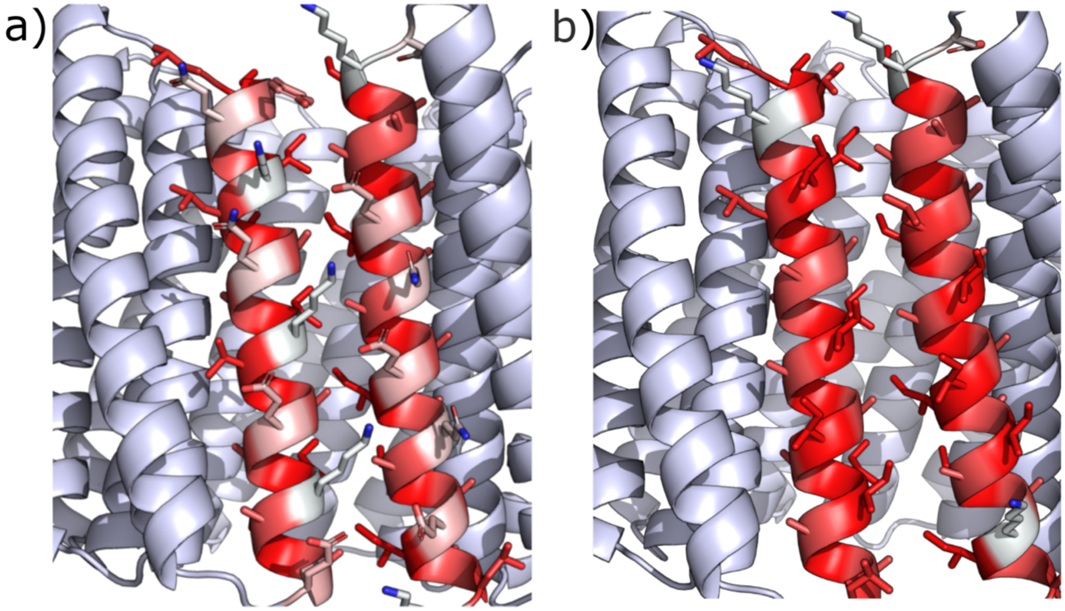

**Supplementary Figure 45. Extra *b-c-f* constraint for water exposed *N*- and *C*-terminal helices of sc-CC-8.** b-c-f positions were fixed to Lys-Glu-Gln to reduce the exposed hydrophobic surface as ProteinMPNN preferred hydrophobic amino acids at these positions. a) sc-CC-8-58, b) a representative design with no constraint at this interface. White, polar amino acids; red, hydrophobic amino acids.

#### 2.2.9 Biophysical and structural characterization of sc-CC-5, 6, and 8

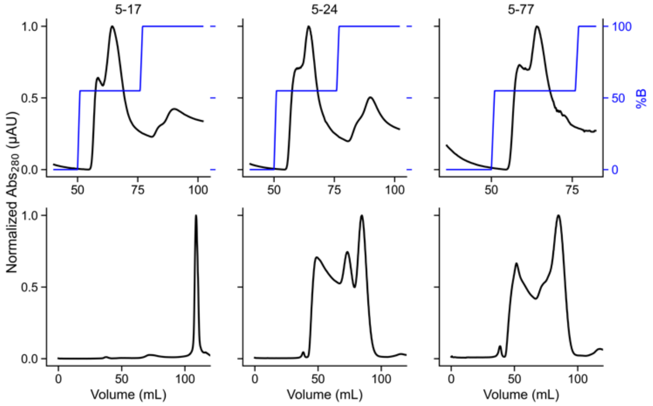

**Supplementary Figure 46. FPLC of sc-CC-5 variants purification.** Sequences for individual proteins can be found in Supplementary Table 20. Top: Ni-affinity chromatography. Fractions eluting at 55% B were pulled and concentrated then further purified by size exclusion chromatography. Conditions: Buffer A: 50 mM Tris, pH 7.4, 150 mM NaCl, 30 mM imidazole, B: 300 mM imidazole. Bottom: Size exclusion chromatography. Fractions eluting at 85 mL were pooled and concentrated to be analysed in further experiments. Conditions: 50 mM sodium phosphate, pH 7.4, 150 mM NaCl.

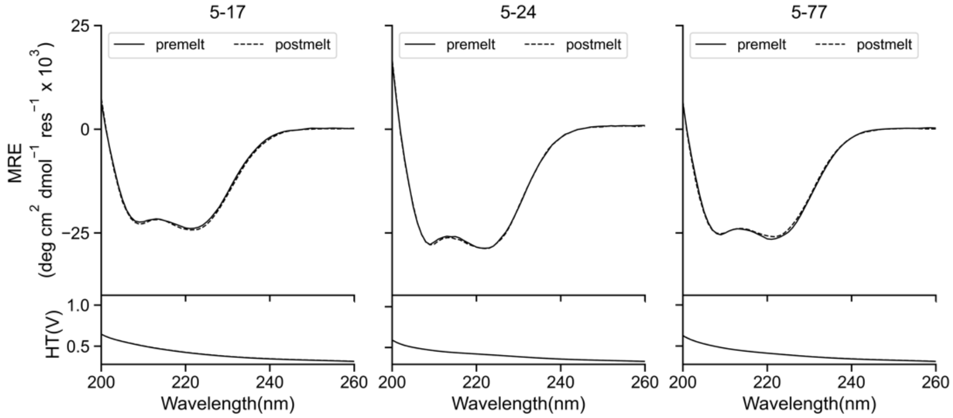

**Supplementary Figure 47. CD spectra of sc-CC-5 proteins designed for this study.** Sequences for individual proteins can be found in Supplementary Table 20. Summary of biophysical data can be found in Supplementary Table 27. Conditions: 5 μM protein, 50 mM sodium phosphate, pH 7.4, 150 mM NaCl, 5 °C. CD scans before thermal ramps up (solid line) and after thermal ramps down (dashed line).

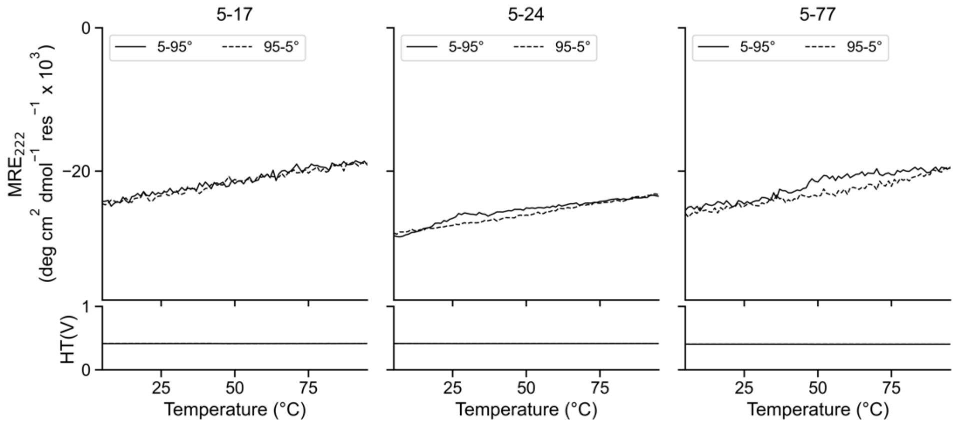

**Supplementary Figure 48. CD thermal-response profiles of sc-CC-5 proteins designed for this study.** Sequences for individual proteins can be found in Supplementary Table 20. Summary of biophysical data can be found in Supplementary Table 27. Conditions: 5 μM protein, 50 mM sodium phosphate, pH 7.4, 150 mM NaCl, 5 – 95 °C. CD scans before ramping up (solid line) and ramping down (dashed line).

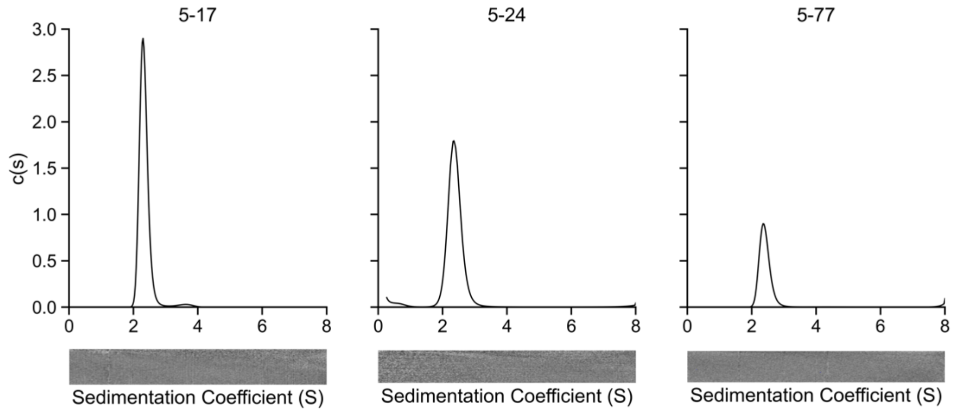

**Supplementary Figure 49. Sedimentation velocity (SV) AUC traces of sc-CC-5 proteins designed for this study.** Sequences for individual proteins can be found in Supplementary Table 20. Fit analysis can be found in Supplementary Table 26. Summary of biophysical data can be found in Supplementary Table 27. Residuals are shown as a bitmap below the fitted data. Conditions: 25 μM protein, 50 mM sodium phosphate, pH 7.4, 150 mM NaCl, 20 °C.

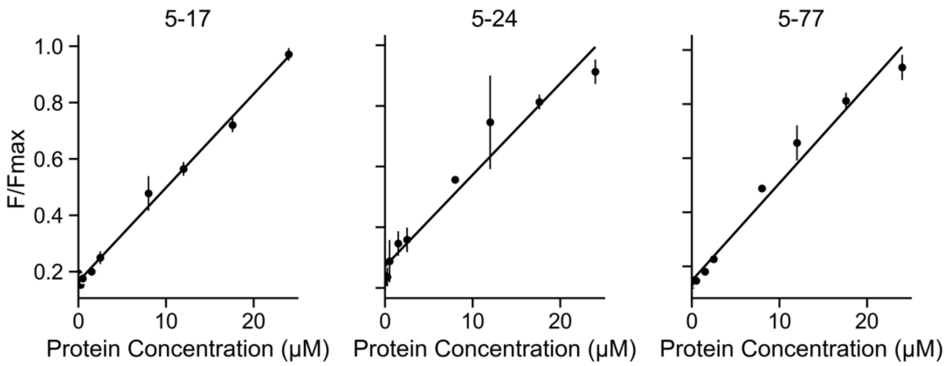

**Supplementary Figure 50. Non-specific binding curves with DPH of sc-CC-5 proteins designed for this study.** Sequences for individual proteins can be found in Supplementary Table 20. Summary of biophysical data can be found in Supplementary Table 27. Conditions: Oligomer concentration 0 – 24 μM protein, 50 mM sodium phosphate, pH 7.4, 150 mM NaCl, 20 °C. Final concentration 0.5 μM DPH (5% v/v DMSO).

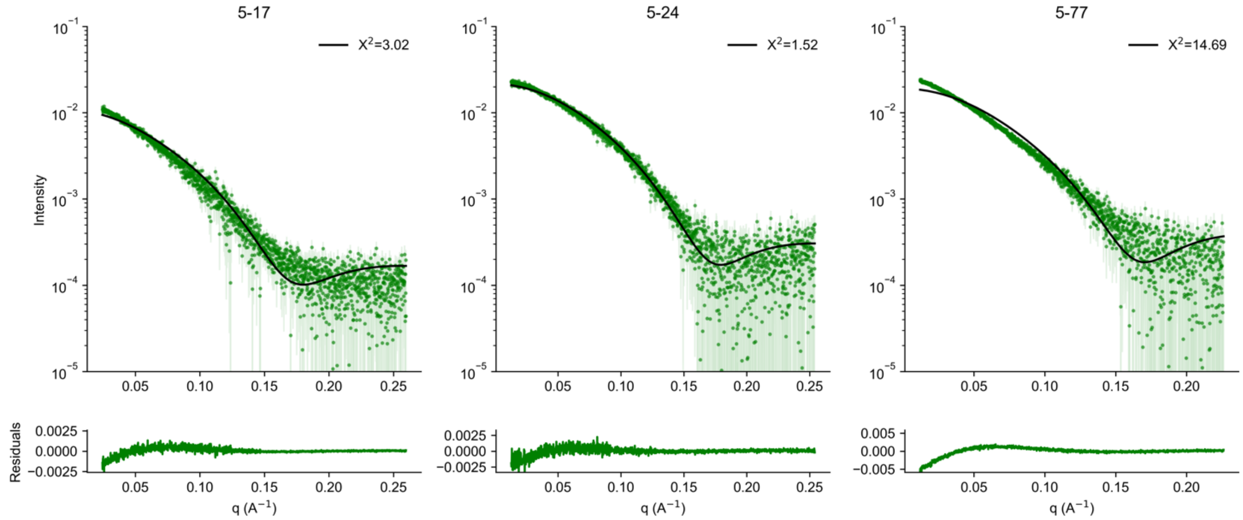

**Supplementary Figure 51. FoXs SAXS profile with fit (top) and residuals (bottom) to AlphaFold2 model of sc-CC-5 proteins designed for this study.** Sequences for individual proteins can be found in Supplementary Table 20. Summary of biophysical data can be found in Supplementary Table 27. Summary of SAXS metrics can be found in Supplementary Table 28. Conditions: 10 mg/mL, 50 mM sodium phosphate, pH 7.4, 150 mM NaCl. *5-77 aggregated at 10 mg/mL, which is observed in the SEC-SAXS collection.

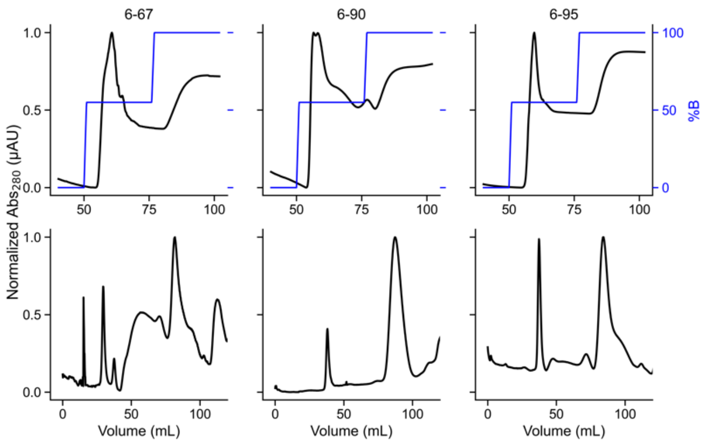

**Supplementary Figure 52. FPLC of sc-CC-6 variants purification.** Sequences for individual proteins can be found in Supplementary Table 22. Top: Ni-affinity chromatography. Fractions eluting at 55% B were pulled and concentrated then further purified by size exclusion chromatography. Conditions: Buffer A: 50 mM Tris, pH 7.4, 150 mM NaCl, 30 mM imidazole, B: 300 mM imidazole. Bottom: Size exclusion chromatography. Fractions eluting at 85 mL were pooled and concentrated to be analysed in further experiments. Conditions: 50 mM sodium phosphate, pH 7.4, 150 mM NaCl.

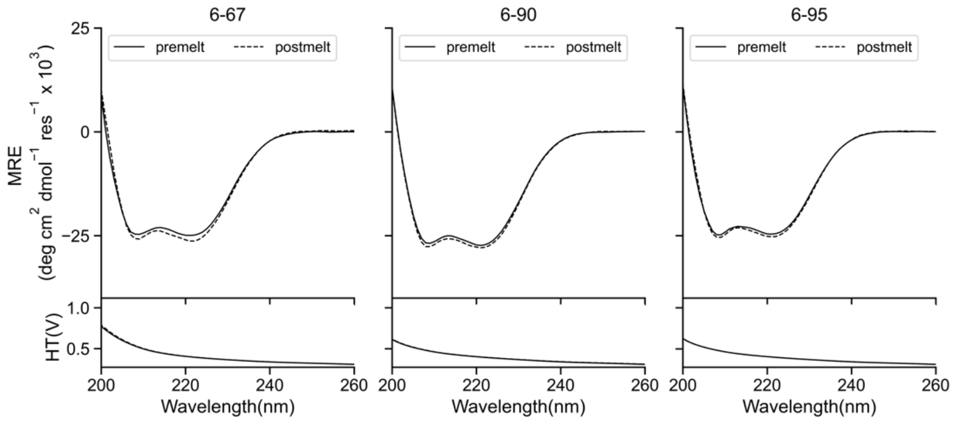

**Supplementary Figure 53. CD spectra of sc-CC-6 proteins designed for this study.** Sequences for individual proteins can be found in Supplementary Table 22. Summary of biophysical data can be found in Supplementary Table 27. Conditions: 5 µM protein, 50 mM sodium phosphate, pH 7.4, 150 mM NaCl, 5 °C. CD scans before thermal ramps up (solid line) and after thermal ramps down (dashed line).

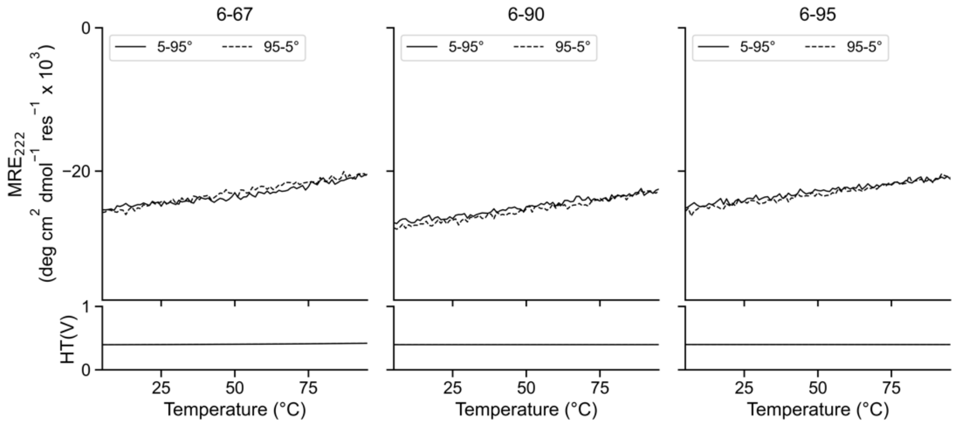

**Supplementary Figure 54. CD thermal-response profiles of sc-CC-6 proteins designed for this study.** Sequences for individual proteins can be found in Supplementary Table 22. Summary of biophysical data can be found in Supplementary Table 27. Sequences for individual proteins can be found in Supplementary Table 1. Conditions: 5 μM protein, 50 mM sodium phosphate, pH 7.4, 150 mM NaCl, 5 – 95 °C. CD scans before ramping up (solid line) and ramping down (dashed line).

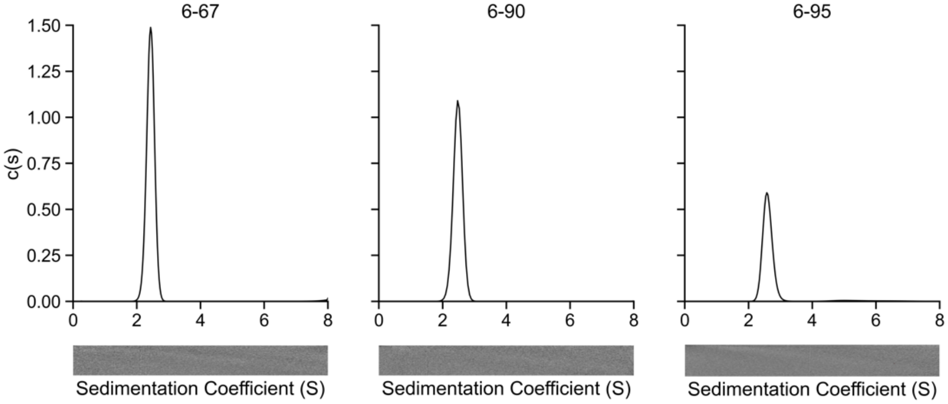

**Supplementary Figure 55. Sedimentation velocity (SV) AUC traces of sc-CC-6 proteins designed for this study.** Sequences for individual proteins can be found in Supplementary Table 22. Fit analysis can be found in Supplementary Table 26. Summary of biophysical data can be found in Supplementary Table 27. Residuals are shown as a bitmap below the fitted data. Conditions: 25 μM protein, 50 mM sodium phosphate, pH 7.4, 150 mM NaCl, 20 °C.

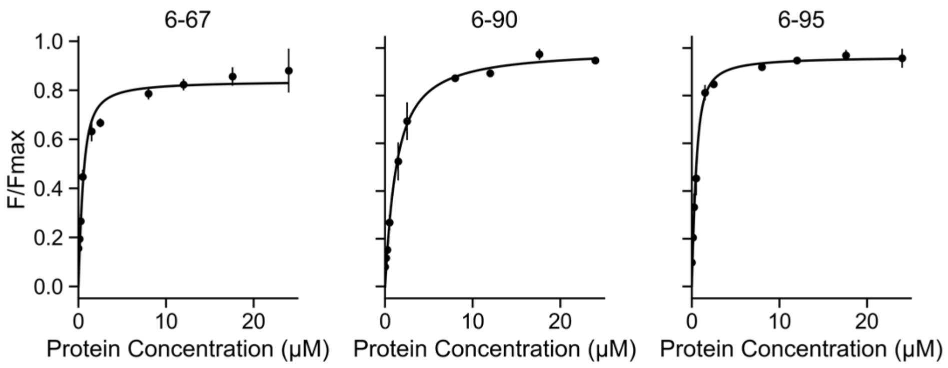

**Supplementary Figure 56. Saturation binding curves with DPH of sc-CC-6 proteins designed for this study.** Sequences for individual proteins can be found in Supplementary Table 22. Summary of biophysical data can be found in Supplementary Table 27. Conditions: Oligomer concentration 0 – 24 μM protein, 50 mM sodium phosphate, pH 7.4, 150 mM NaCl, 20 °C. Final concentration 0.5 μM DPH (5% v/v DMSO).

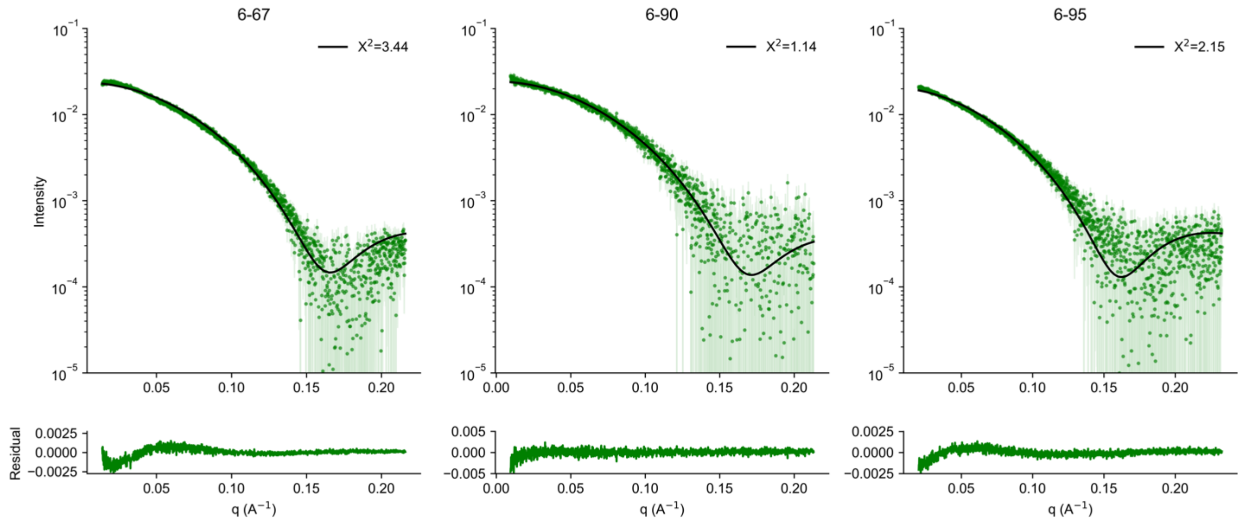

**Supplementary Figure 57. FoXs SAXS profile with fit (top) and residuals (bottom) to AlphaFold2 model of sc-CC-6 proteins designed for this study.** Sequences for individual proteins can be found in Supplementary Table 22. Summary of biophysical data can be found in Supplementary Table 27. Summary of SAXS metrics can be found in Supplementary Table 28. Conditions: 10 mg/mL, 50 mM sodium phosphate, pH 7.4, 150 mM NaCl.

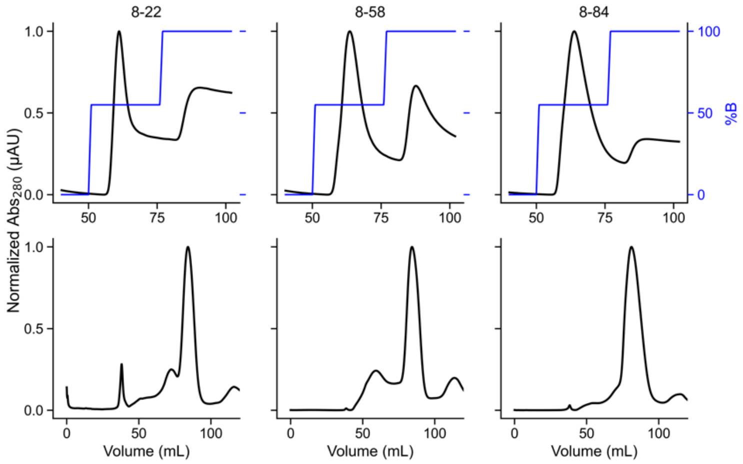

**Supplementary Figure 58. FPLC of sc-CC-8 variants purification.** Sequences for individual proteins can be found in Supplementary Table 24. Top: Ni-affinity chromatography. Fractions eluting at 55% B were pulled and concentrated then further purified by size exclusion chromatography. Conditions: Buffer A: 50 mM Tris, pH 7.4, 150 mM NaCl, 30 mM imidazole, B: 300 mM imidazole. Bottom: Size exclusion chromatography. Fractions eluting at 85 mL were pooled and concentrated to be analysed in further experiments. Conditions: 50 mM sodium phosphate, pH 7.4, 150 mM NaCl.

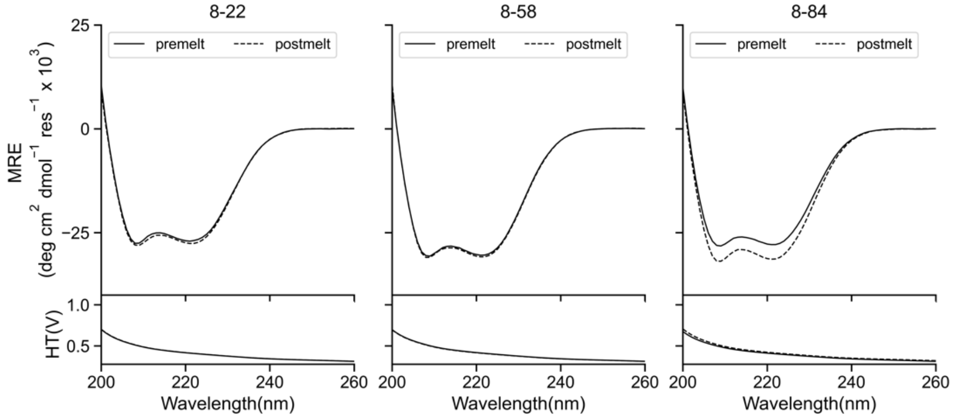

**Supplementary Figure 59. CD spectra of sc-CC-8 proteins designed for this study.** Sequences for individual proteins can be found in Supplementary Table 24. Summary of biophysical data can be found in Supplementary Table 27. Conditions: 5 μM protein, 50 mM sodium phosphate, pH 7.4, 150 mM NaCl, 5 °C. CD scans before thermal ramps up (solid line) and after thermal ramps down (dashed line).

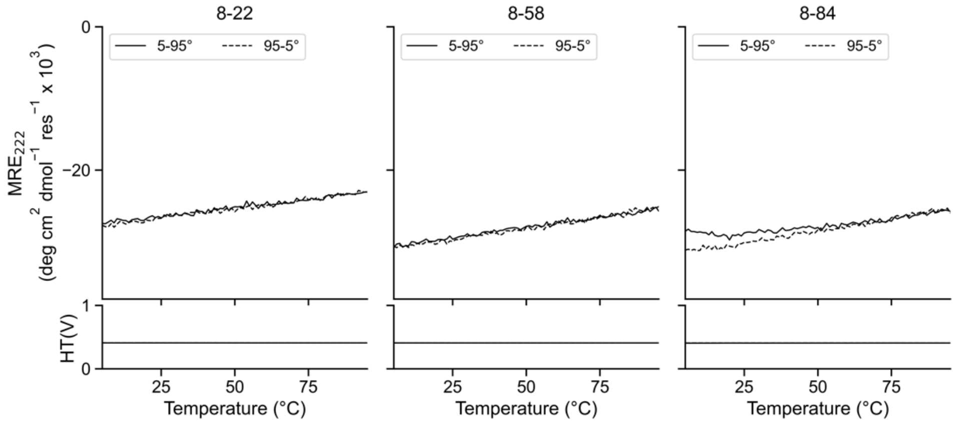

**Supplementary Figure 60. CD thermal-response profiles of sc-CC-8 proteins designed for this study.** Sequences for individual proteins can be found in Supplementary Table 24. Summary of biophysical data can be found in Supplementary Table 27. Conditions: 5 µM protein, 50 mM sodium phosphate, pH 7.4, 150 mM NaCl, 5 – 95 °C. CD scans before ramping up (solid line) and ramping down (dashed line).

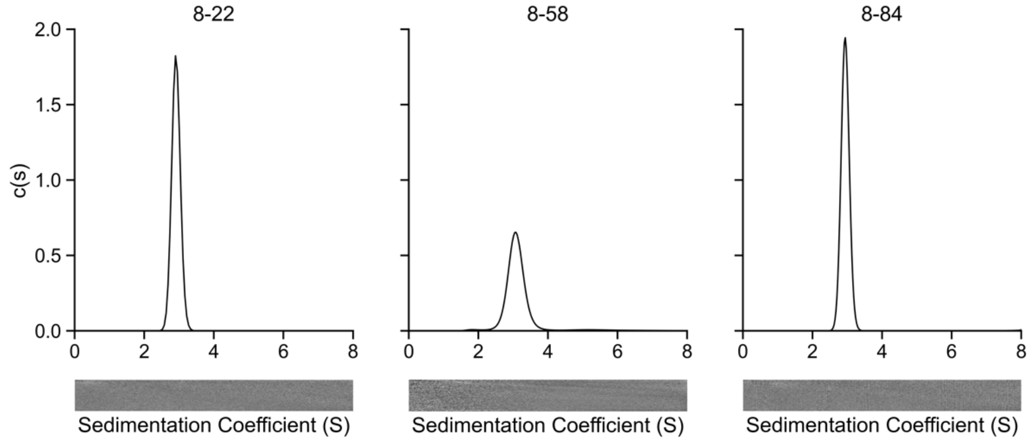

**Supplementary Figure 61. Sedimentation velocity (SV) AUC traces of sc-CC-8 proteins designed for this study.** Sequences for individual proteins can be found in Supplementary Table 24. Fit analysis can be found in Supplementary Table 26. Summary of biophysical data can be found in Supplementary Table 27. Residuals are shown as a bitmap below the fitted data. Conditions: 25 µM protein, 50 mM sodium phosphate, pH 7.4, 150 mM NaCl, 20 °C.

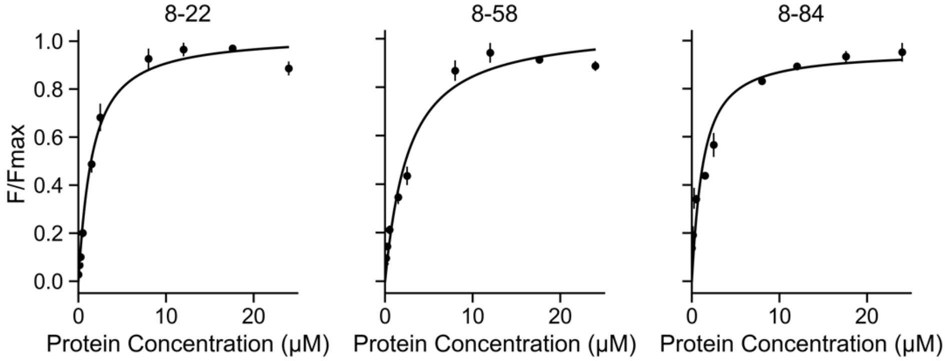

**Supplementary Figure 62. Saturation binding curves with DPH of sc-CC-8 proteins designed for this study**. Sequences for individual proteins can be found in Supplementary Table 24. Summary of biophysical data can be found in Supplementary Table 27. Conditions: Oligomer concentration 0 – 24 µM protein, 50 mM sodium phosphate, pH 7.4, 150 mM NaCl, 20 °C. Final concentration 0.5 µM DPH (5% v/v

DMSO).

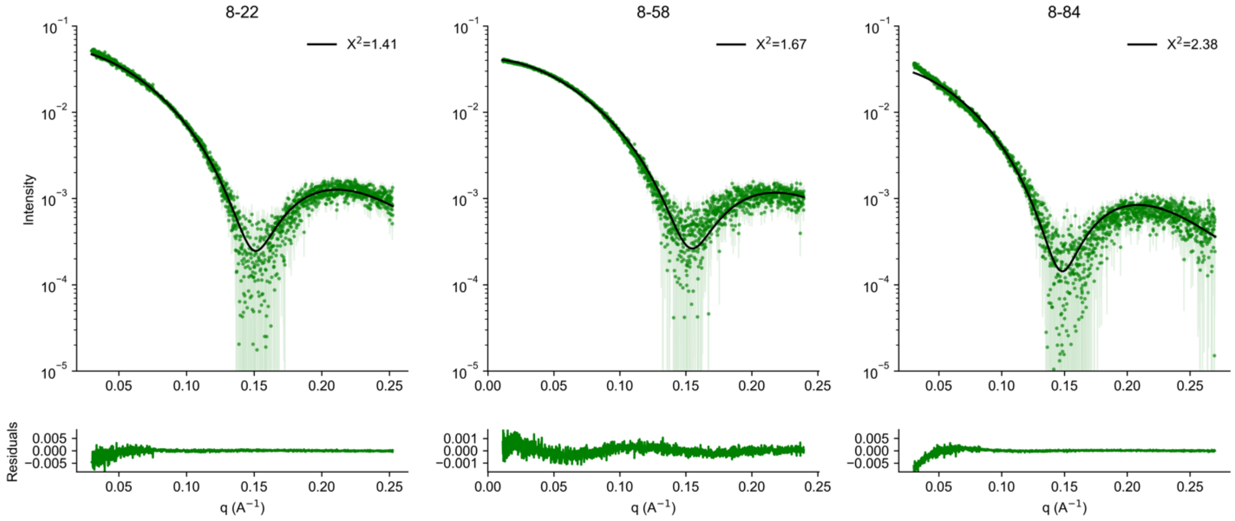

**Supplementary Figure 63. FoXs SAXS profile with fit (top) and residuals (bottom) to AlphaFold2 model of sc-CC-8 proteins designed for this study.** Sequences for individual proteins can be found in Supplementary Table 24. Summary of biophysical data can be found in Supplementary Table 27. Summary of SAXS metrics can be found in Supplementary Table 28. Conditions: 10 mg/mL, 50 mM sodium phosphate, pH 7.4, 150 mM NaCl.

#### 2.2.10 Structural analysis of antiparallel and parallel single-chain proteins

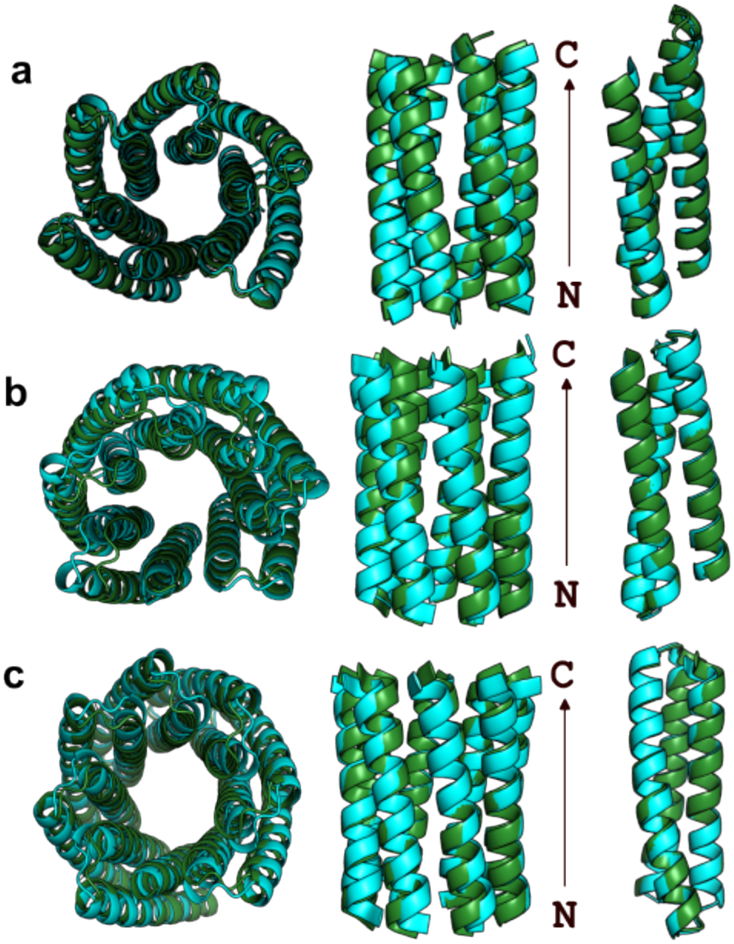

**Supplementary Figure 64. Overlays showing the starting backbone model (green) and the X-ray structure for (a) sc-CC-6-95 (PDB id, 8qag), (b) sc-CC-7-LI (PDB id, 8qai) and (c) sc-CC-8-58 (PDB id, 8qah).** Backbone Cα RMSD values available in Supplementary Table 31.

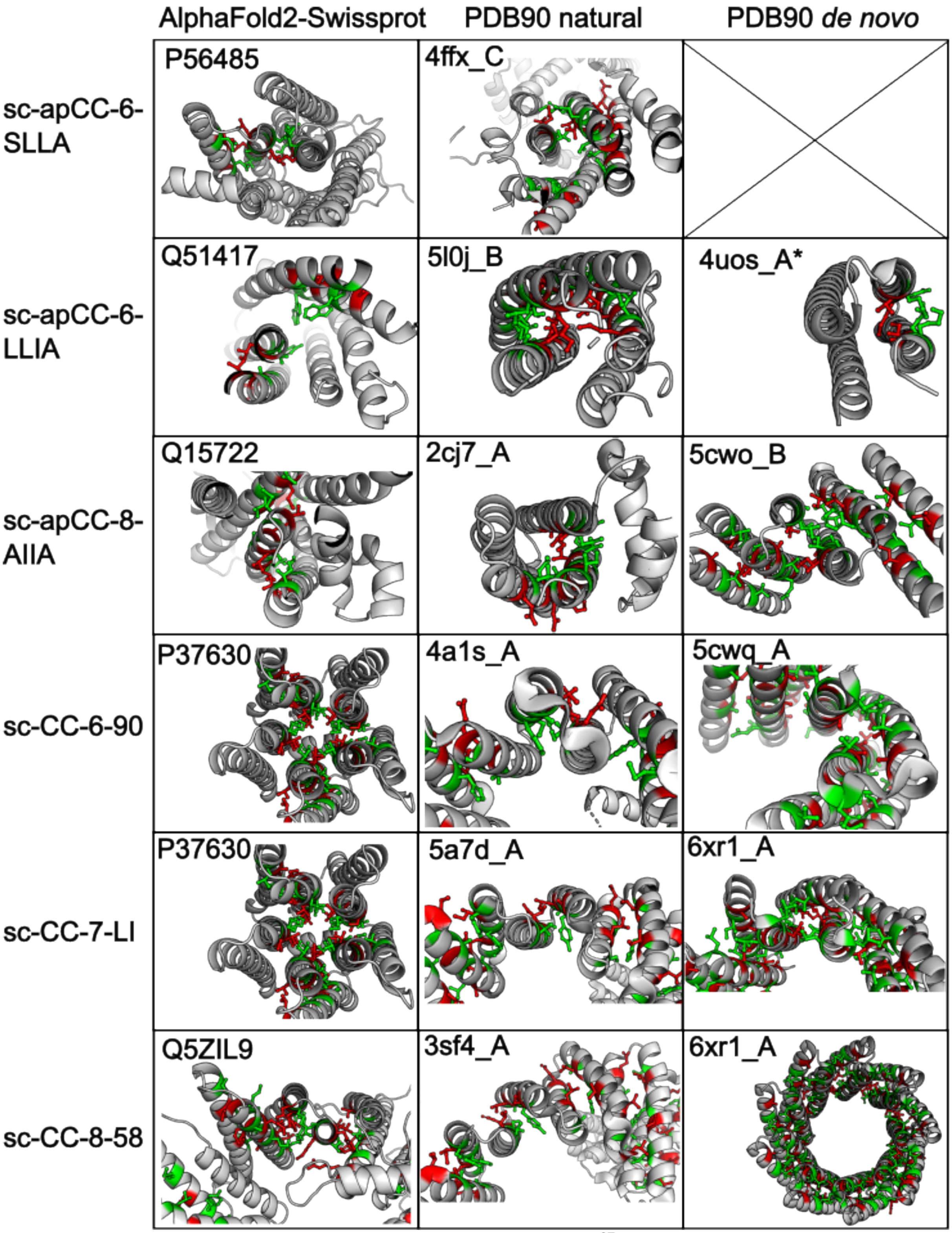

**Supplementary Figure 65. Socket2 analysis of top Foldseek[37] structural alignments against PDB90[39,40] and AlphaFold2 - Swiss-Prot databases[5,38].** 7.5 Å Socket2[41] packing cutoff; a-knobs in red, d-knobs in green. *4uos[42] contains non-canonical coiled-coil repeats of designed KIH contacts and hydrogen-bonding networks which are not detected by Socket2[41]. P37630 is a 5-helix αHB, other structures contain only 2- or 3-helix bundles.

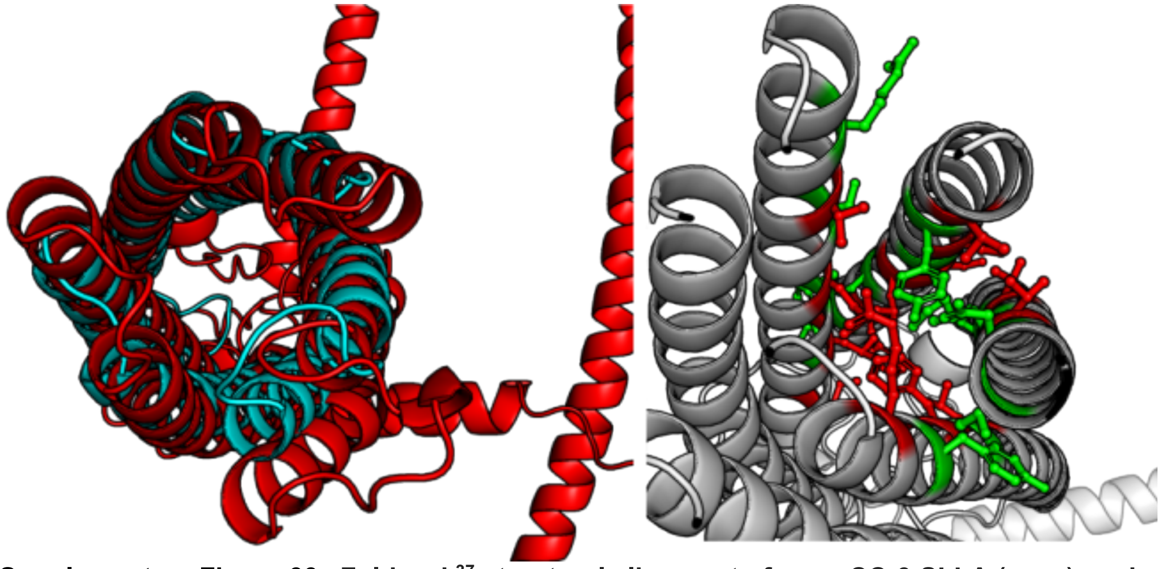

**Supplementary Figure 66. Foldseek[37] structural alignment of sc-apCC-6-SLLA (cyan) against an uncharacterised protein from *A. japonicus* (red) (UniProt id, A0A2G8LCW8).** TM-score = 0.402, RMSD = 8.26 Å, 14.7% sequence identity. Socket2**[41]** with 7.5 Å packing cutoff identifies repeating 2-helix bundles (a-knobs in red, d-knobs in green).

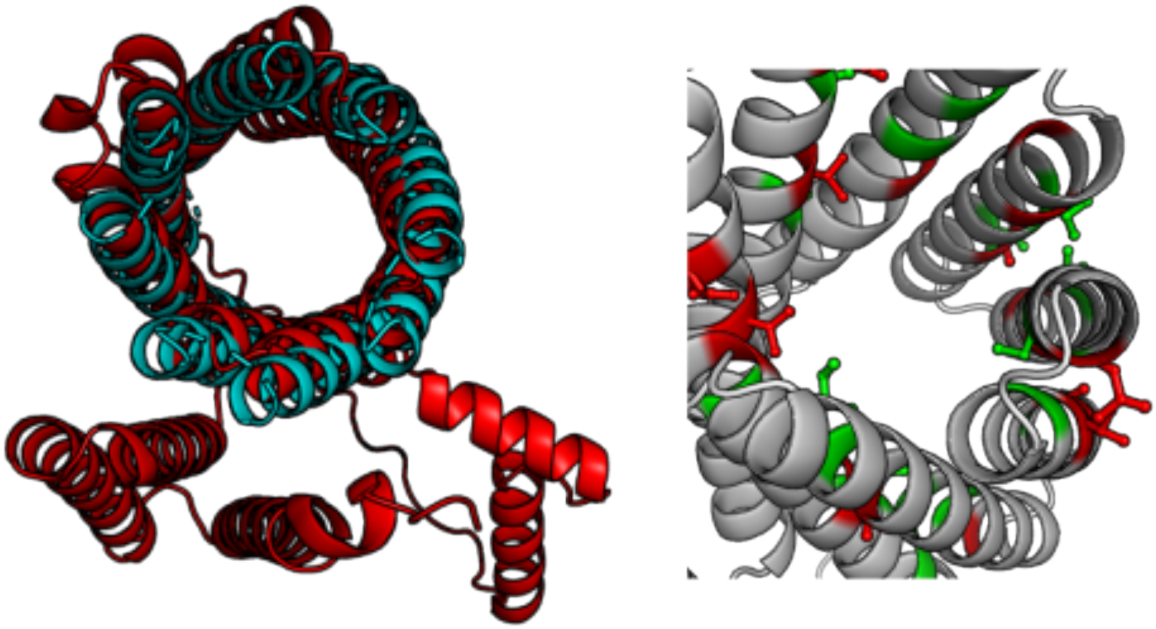

**Supplementary Figure 67. Foldseek[37] structural alignment of sc-apCC-8-AIIA (cyan) against an uncharacterised protein from *C. caeruleus* (red) (UniProt id, A0A3N1FT86).** TM-score = 0.626, RMSD = 5.14 Å, 7% sequence identity. Socket2**[41]** with 7.5 Å packing cutoff identifies repeating 2-helix bundles (a-knobs in red, d-knobs in green).
